# Reconstructing a Missing Link of HIV-1 Assembly: HIV-1 Envelope-Matrix Interactions in a Native Viral Context

**DOI:** 10.64898/2026.01.15.699503

**Authors:** Jacob T. Croft, Hung N. Do, Daniel P. Leaman, Klaus N. Lovendahl, Pooja Ralli-Jain, Katelyn J. Chase, Cynthia A. Derdeyn, Michael B. Zwick, S. Gnanakaran, Kelly K. Lee

**Affiliations:** Department of Medicinal Chemistry, University of Washington, Seattle, WA, USA; Theoretical Biology and Biophysics Group, Theoretical Division, Los Alamos National Laboratory, Los Alamos, NM, USA 87545; Department of Immunology and Microbiology, The Scripps Research Institute, La Jolla, CA 92037, USA; Department of Laboratory Medicine & Pathology, University of Washington School of Medicine, Seattle, Washington 98195, USA

**Author notes:** These authors contributed equally (Jacob T. Croft, Hung N. Do).

**Keywords:** HIV-1, maturation, cryo-electron tomography, subtomogram averaging, Env, glycoprotein, Gag, matrix protein, retrovirus, cytoplasmic tail, protein dynamics, molecular dynamics, virus structure, membrane

## Abstract

The envelope surface glycoprotein (Env) on HIV-1 drives cell entry and genome delivery through its receptor binding and membrane fusion activities. Incorporation of Env onto assembling virions is governed by its cytoplasmic tail (Env-CT) and the matrix (MA) domain of the PR55^Gag^ (Gag) polyprotein. To better understand how Gag recruits Env onto virions while reducing its antigenic profile by restricting Env copy number to low levels, we used cryo-electron tomography (cryo-ET) and subtomogram averaging combined with molecular dynamics simulations to investigate Env-MA association in intact viral particles. Full-length Env-CT was resolved directly over individual MA trimers in regions of MA lattice discontinuity. We observed that the conserved but enigmatic Kennedy-sequence motif in Env-CT forms a key linkage between Env and MA. Mutational analysis confirmed the impact of specific CT and MA interactions on Env incorporation. Gag maturation released MA lattice restraints on Env, facilitating its clustering and promoting fusion activity.

## Introduction

Assembly of human immunodeficiency virus type 1 (HIV-1) virions is a dynamic process involving the interaction of Gag and Gag-Pol polyproteins, viral genomic RNA, membrane, and the cytoplasmic tail domain (CT) of the Envelope (Env) glycoprotein. While Gag assembly has been characterized in detail ^1,2^, a key missing link in our understanding of HIV-1 assembly centers around the interactions between Env and Gag that lead to incorporation of Env, which drives receptor binding and membrane fusion and is thus essential for production of infectious virions.

The N-terminal matrix (MA) subunit of the structural polyprotein PR55^Gag^ (Gag) plays an essential role in particle formation as it targets Gag to the plasma membrane through binding to the negatively charged lipid phosphatidylinositol 4,5-bisphosphate (PIP2), and insertion of its hydrophobic myristoyl chain into the membrane ^3–6^. Furthermore, MA also mediates Env recruitment onto viral particles through poorly understood interactions with its long cytoplasmic tail (CT) ^7–18^, and seemingly paradoxically, is responsible for restricting Env incorporation to approximately only 7-14 copies per virion ^19,20^. The low level of Env incorporation is a defining characteristic of HIV-1 that is believed to have evolved as a means of reducing HIV-1’s antigenic profile by hampering bivalent antibody binding ^21^.

Following viral budding, the viral protease cleaves Gag into matrix (MA), capsid (CA), nucleocapsid (NC), and p6 subunits, along with two spacer peptides (SP1 and SP2) in a process termed maturation ^1^. HIV-1 maturation enables condensation of CA subunits into a protective conical core, which shields the viral RNA dimer from detection by innate immunity surveillance and facilitates its trafficking into the nucleus (Fig 1A) ^22–26^. Maturation also leads to reorganization of membrane-associated MA ^27,28^, and through still poorly understood mechanisms, has been reported to enhance the ability of Env to efficiently mediate membrane fusion and cell entry ^29–31^.

**Figure 1.**
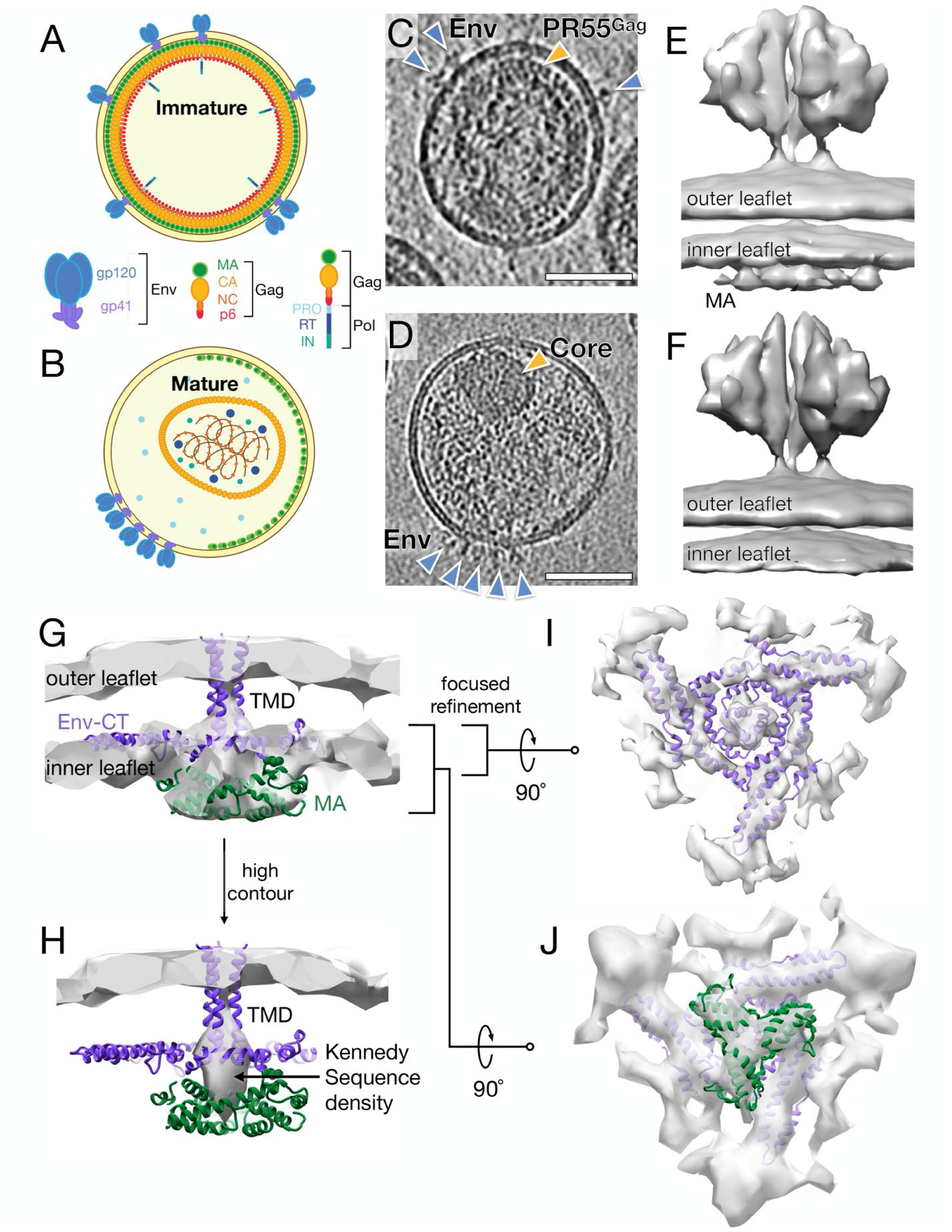
Cryo-ET of HIV-1 maturation. **(A)** Cartoon depicting immature and **(B)** mature HIV-1 virions. The immature virion contains a membrane associated PR55^Gag^ lattice, in which Env is dispersed on the surface. In the mature HIV-1 virion, the PR55^Gag^ has been cleaved into its component domains, causing rearrangement of the CA subunits to form a conical capsid and rearrangement of the MA lattice at the viral membrane. On the surface, Env forms clusters. **(C-D)** Representative 10 nm tomographic slices of a darunavir-treated immature **(C)** and mature **(D)** HIV-1 VLPs bearing CH505.N197D are shown. Env is indicated by blue arrowheads and the PR55^Gag^ lattice is indicated by the yellow arrowhead in the immature VLP. Scale bars are 50 nm. **(E, F)** Initial subtomogram averages, produced from global alignment of binned particles with a cylindrical mask, encompassing the ectodomain and membrane for Env on immature **(E)** and mature **(F)** virions. In panel E, additional density is indicated corresponding to the MA layer that is not present in the mature average. **(G-J)** Focused refinement was performed on the inner membrane beneath Env from immature virions. Panels G and H depict side views with different electron density thresholds. Panel I depicts the inner membrane leaflet viewed from the outside. Panel J depicts the inner leaflet and MA viewed from inside the virion. The model of the Env CT (PDB:7LOH) is colored purple and fit into the maps. A single MA trimer (green) is fit into the immature map (PDB:7OVQ).

Interaction between the Env-CT and MA in immature particles has been proposed to affect Env ectodomain conformation, activity ^32–36^, and clustering of Env on viral particles ^36–39^. It is also conceivable that changes in the MA and Gag lattices could impact Env stability and activity indirectly by modulating membrane order and plasticity. Env-MA and membrane interactions are therefore key determinants of viral infectivity, however a mechanistic understanding of Env-MA coupling and the resulting downstream functional effects have been limited by a lack of structural information for Env and MA *in situ* in virions.

HIV-1 Env is synthesized in the endoplasmic reticulum as an immature precursor (gp160), which undergoes trimerization, glycosylation, and processing into gp120 and gp41 subunits in the Golgi ^1^. Env then traffics as a trimer to the cell surface for incorporation into virions. Structural and functional studies have been heavily focused on the Env ectodomain, while a detailed characterization of membrane-interactive portions of Env on the exterior as well as the transmembrane domain (TM) and long, ∼150 amino acid-long CT have been critically lacking, particularly in the context of intact viral particles with biological lipid membranes. Env-CT contains interspersed highly conserved regions that point to functional roles in viral replication. A negatively charged segment known as the Kennedy sequence (KS; also referred to as “Kennedy epitope” or “Conserved Hydrophilic Domain”) has been proposed to be externalized on the host cell membrane where it could present immunogenic epitopes ^40,41^, while other studies placed this segment on the virus interior ^42–45^. Additionally, three α-helical rich regions located in the C-terminal part of the Env-CT, known as lentiviral lytic peptide (LLP) 1-3 have been shown to form membrane-associated amphipathic helices ^43,46,47^. Recent NMR studies have begun to provide structural insights using detergent or bicelle-solubilized, truncated CT constructs lacking the large trimeric ectodomain, which indicate that the LLP helices form a flat, compact “baseplate” structure beneath the TM ^44,45^. It remains to be determined, however, whether the fragmentary NMR structural models reflect the organization and interactions of native CT in a virus particle where the tail is embedded in a distinct lipid environment with raft-like composition and notable bilayer asymmetry ^48–50^, as well as being constrained by its connection to the TM, ectodomain, and interactions with MA on the virus interior.

Biochemical and mutagenesis studies have pointed to a direct engagement between Env and MA ^8,9,11,13,15,16,18,51–60^. For example, it has been reported that when the viral membrane is removed by detergent treatment, Env is largely retained with immature viral cores following sedimentation, in a manner dependent on the presence of the CT ^16,18^. Additionally, mutations designed to disrupt the Env-CT baseplate resulted in defects in Env incorporation in certain cell types ^60^. A structural framework is needed to connect the observations and elucidate the mechanism of engagement between the Env-CT, MA, and the viral membrane that mediate specific but apparently restrained Env incorporation onto virions.

Here, to investigate the molecular interactions between the Env-CT and Gag-MA *in-situ*, we performed cryo-electron tomography (cryo-ET) on both mature and immature viral particles. Using sub-tomogram averaging, we reconstructed full-length Env-CT in complex with MA *in situ*. Aided by MD simulations and mutagenesis of residues predicted to impact interaction between Env-CT and Gag-MA, we mapped key interactions between the two proteins. Together, this integrative analysis of immature and mature HIV-1 particles provides new insights into the interactions between essential viral components and the changes resulting from maturation that transform them into infectious particles. The structural insights inform future targeting of therapeutics to disrupt Env incorporation and virus assembly by blocking these essential interactions.

## Results

### Env-CT is located directly above an MA trimer in immature HIV-1 VLPs

We first performed cryo-ET on mature and immature HIV-1 virus-like particles (VLPs) bearing the full-length CH505.N197D Env, which is derived from the transmitted/founder primary isolate CH505 (with N197 glycosylation site near the CD4 binding site removed) ^61^. Immature and mature VLPs (Fig 1A-D) were cultured and harvested in identical fashion, with the exception that the protease-inhibitor darunavir was used to generate immature VLPs (Fig S1A). In raw tomograms, darunavir-treated VLPs can clearly be seen with an intact Gag lattice beneath the membrane (Fig 1C). By contrast, in mature VLPs, an organized Gag lattice was not observed (Fig 1D), although the AT-2 treatment used to ensure the samples were non-infectious is anticipated to alter the internal architecture of the condensed CA/NC core through its modification of the NC-RNA interaction ^62^, and distinct conical cores were only identifiable in some instances.

In a previous study of immature particles, we observed that Env appears to be positioned directly over Gag ^63^, rather than residing over the holes in the lattice as had been presumed in some models of HIV-1 assembly ^12^. At the time, the electron-dense CA lattice was the most prominent feature in immature Gag particles, while the small, membrane-associated MA subunits posed a greater challenge to reliably visualize. Here, utilizing larger datasets and improved tomography analysis, we have been able to visualize Env in relation to Gag-MA in intact VLPs.

Subtomogram averaging was performed on CH505.N197D full-length Env from immature and mature VLPs (Fig S2). First, alignment of binned particles with a cylindrical mask was performed to obtain initial, low-resolution structures including the Env ectodomain and the viral membrane underneath (Fig 1E, F, Fig S2). The ectodomains for both maturation states appeared nearly identical at this resolution (Fig 1E-F). However, on the virus interior, density corresponding to MA was clearly visible beneath Env in close association with the inner membrane leaflet of immature VLPs (Fig 1E) but absent in mature VLPs despite applying the same masking strategy (Fig 1F).

We next performed local refinement focused on the membrane inner leaflet and internal region that is anticipated to contain MA in immature VLPs. Density consistent with an MA trimer was resolved directly underneath a triangular density feature that showed excellent agreement with the reported CT “baseplate” previously determined by NMR using truncated tail-based constructs ^64,65^ (Fig 1G-J). The density threshold shown in panel G, was chosen to show the density features for both the Env-CT and MA, but since the tail is positioned in a similar plane to the inner membrane leaflet phospholipid head group layer, at this contouring, Env-CT density is obscured by surrounding membrane density. A comparable focused refinement in the mature VLPs resolved a similar CT baseplate feature as in the immature reconstruction, however no equivalent MA density was observed beneath Env (Fig S2D).

Close up views of available isolated CT and MA models rigidly fit into the density are shown in Figs 1I and 1J respectively. The resolution of both a single MA trimer and the Env-CT in a single map indicates that a specific interaction and positioning exists between the two proteins. A single, central transmembrane density is seen crossing the membrane in this structure (Fig 1G), consistent with the trimeric, helical bundle TM conformation in the NMR model ^64,65^.

While our cryo-ET map does not directly resolve interacting residues between MA and CT, certain regions show stronger connecting density at higher thresholds. The strongest connection appears directly underneath the TM, where a solid plug of density extends from the CT to fill the central hole of the MA trimer (Fig 1H). We hypothesize that the polypeptide segment that follows the TM (residues 710-742) including the KS interacts as a loop with the MA trimer along its central axis. The strong connecting density, along with the positioning of the CT directly over the MA with robust density along the central 3-fold axis of an MA trimer, provides an explanation of why many studies have implicated a link between MA trimerization and Env incorporation ^13,51,66^ by direct interaction of an intact MA trimer with the Env-CT.

Notably, despite the presence of ordered Gag CA lattice in the immature particles, only a single central MA trimer was resolved interacting with the CT. This could indicate that Env with full-length tails may preferentially localize to areas of broken MA lattice or the tail itself may induce local lattice defects. This effect is further examined in molecular dynamics (MD) simulations described below.

### Env positioning relative to MA is conserved in ADA.CM.755* VLPs

Next, we performed subtomogram averaging on mature and immature high-Env copy number VLPs (hVLP) bearing the ADA.CombMut Env (ADA.CM.755*) ^67,68^, which has a partial CT truncated after residue 755. These VLPs incorporate higher levels of Env than typical HIV-1 particles, making them more amenable for higher resolution subtomogram averaging (Fig 2A, S3). The 755* tail truncation removes much of the “LLP” portions of the CT baseplate structure, while the KS and the N-terminal portion of the H1 helix of the LLP-2 region remain. We previously demonstrated that despite partial CT truncation, Env on these VLPs shows consistent positioning above the Gag lattice as full-length ADA.CM Env ^63^ (Fig S4), indicating that in the truncated form, at least a subset of key residues responsible for MA binding are retained, contrasting with studies that highlighted C-terminal segments of the tail as mediating Env incorporation ^7,51^.

**Figure 2.**
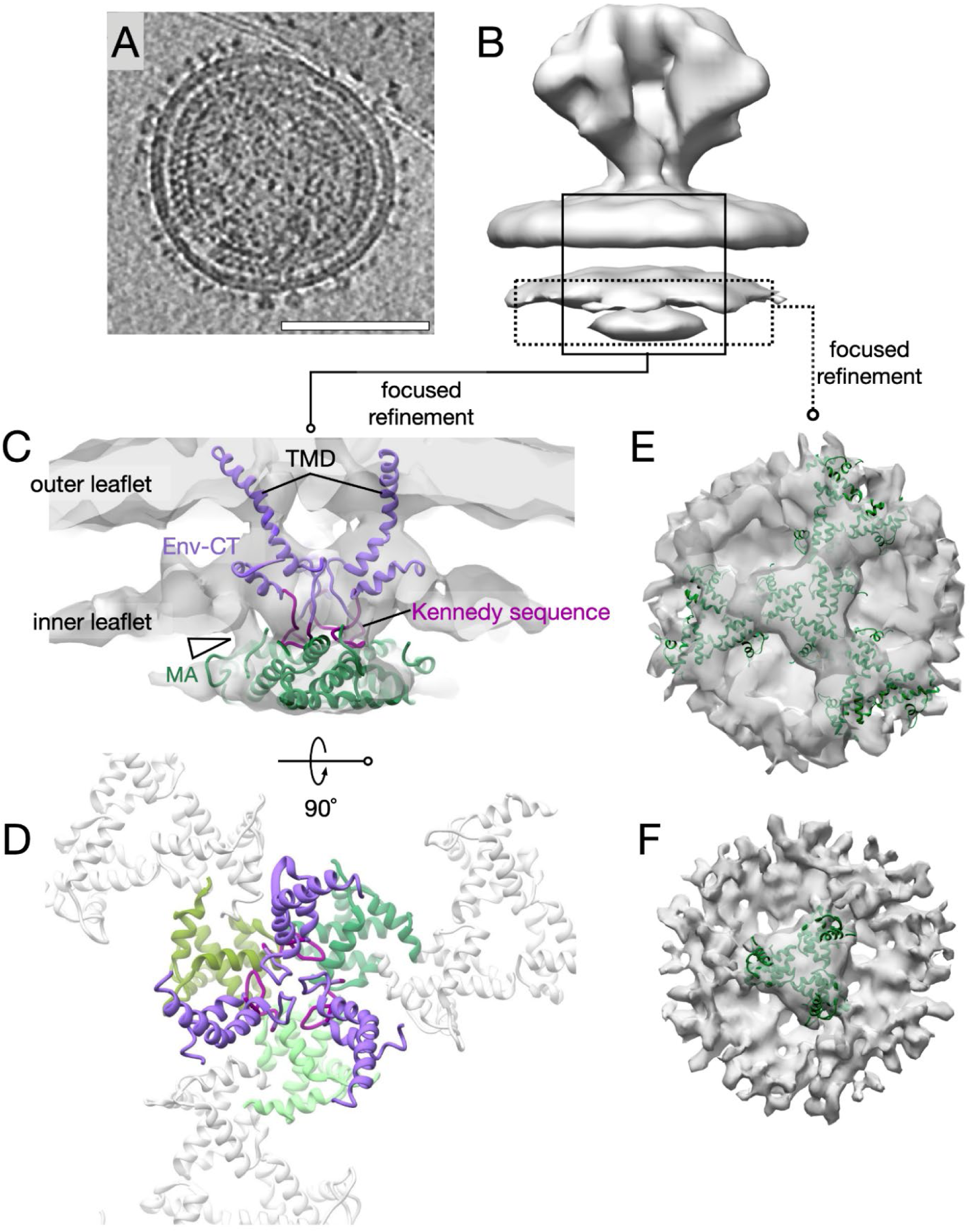
Subtomogram average of the immature ADA.CM.755* Env CT-MA interaction. **(A)** Representative 10 nm thick tomographic slice of an immature (darunavir-teated) VLP bearing ADA.CM.755* Env. Scale bar 100 nm. **(B)** Initial subtomogram average of immature ADA.CM.755* Env produced from global alignment of binned particles with a cylindrical mask encompassing the ectodomain and membranes. Areas masked for focused refinement are indicated in boxes. **(C-D)** Focused, masked refinement of both membranes resulted in a map in which 3 separate TM densities are resolved over an unclear MA density. Homology models of 3 subunits of the ADA.CM.755* Env CT are colored purple and fit to the map. The KS-region is colored magenta. **(E-F)** Focused, masked refinement on the inner membrane leaflet and MA layer resolved density consistent with the immature MA lattice (PDB: 7OVQ, colored green). The same map is shown in both panels, at different density thresholds.

According to a previous report ^18^, the Env-CT/MA interaction can withstand treatment with the detergent Triton X-100, and immature Gag particles pelleted after detergent treatment retained bound Env. We sought to further validate whether the observed positioning of Env over the MA trimer in the ADA.CM.755* sample is resistant to treatment of VLPs with Triton X-100. Cryo-ET revealed that Env indeed is retained on de-membranated Gag lattices (Fig S4C). Surprisingly, Env positioning over the lattice in these instances was consistent with the positioning observed for intact ADA.CM.755* ^63^ (Fig S4B) and CH505.N197D VLPs (Fig S4A), indicating persistent, robust Env-CT-MA interaction, even with the absence of membrane and truncation of nearly 100 C-terminal residues of Env-CT in the ADA.CM.755* case.

In accord with the CH505.N197D VLPs, subtomogram averages of Env focused on the membrane and internal membrane-associated region revealed the presence of the MA layer underneath Env in only the immature VLPs but not the mature VLPs (Fig 2B, Fig S3). With ADA.CM.755*, depending on the masking approach used during focused refinement, we either resolved 3 separate TM densities crossing the membrane over unclear MA layer density (Fig 2C-D, S3), or clearly defined density underneath Env that was consistent with the previously determined immature MA lattice (Fig 3E-F, S3). This contrasts with the full-length CH505.N197D Env, in which the positioning of the Env-TM/CT were in register with the single MA trimer and simultaneously resolved, indicating some flexibility in the interaction in the partially tail-truncated case. Additionally, the difference in TM ordering suggests that the presence of the intact CT baseplate may play a role in stabilizing the CH505.N197D transmembrane helices as a trimeric bundle.

**Figure 3.**
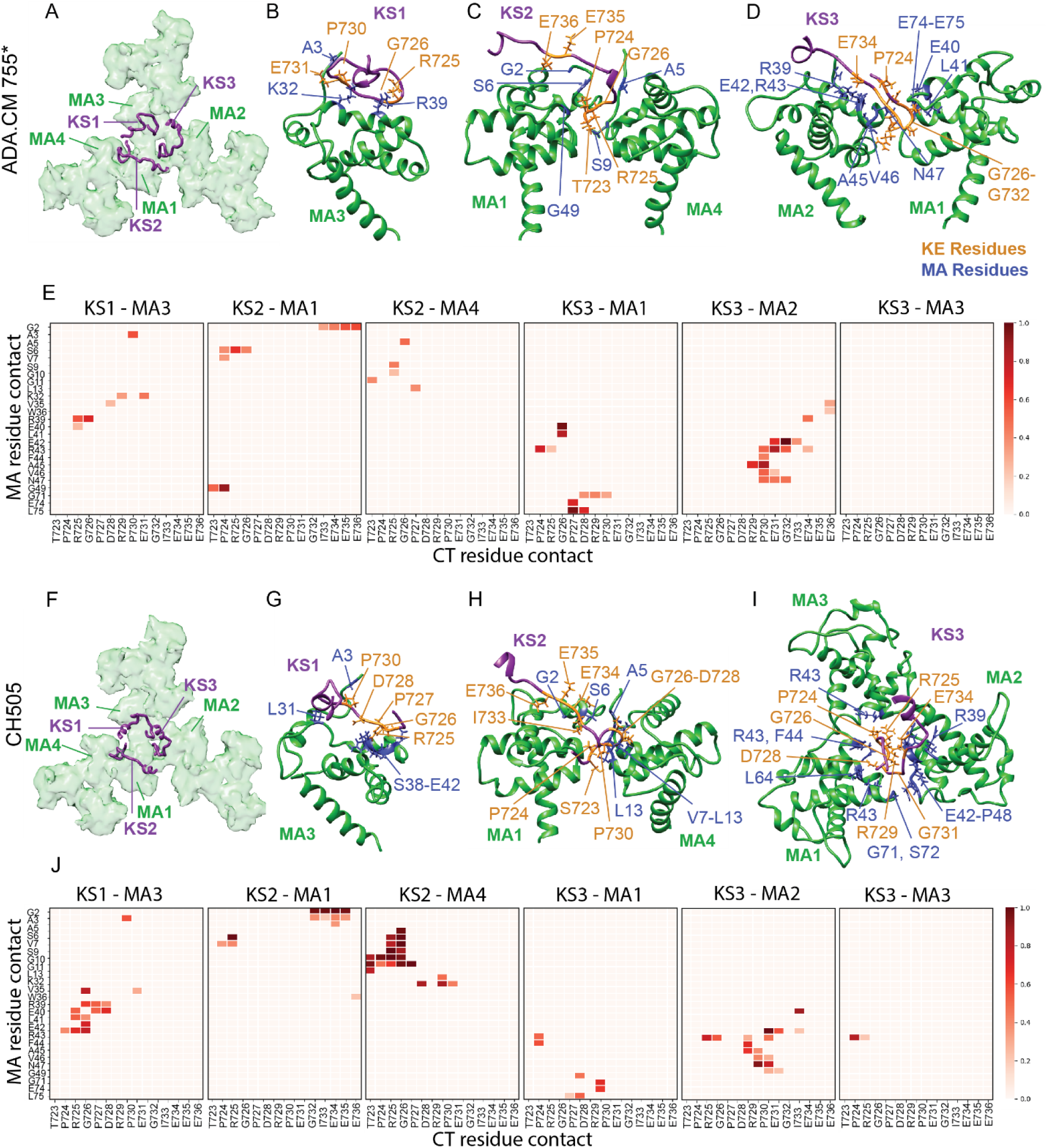
Interactions between the KSs and the immature psg53 Gag MA)protein captured from the density-guided MD simulations. **(A-D)** Representative KS positions relative to the restrained immature MA in the density-guided MD simulations of ADA.CM.755*. **(B)** Residues involved in the interactions between the first KS (KS1) and restrained MA monomer 3 (MA3) of the central MA trimer. **(C)** Important residues involved in the interactions between KS2, restrained MA1 of the central MA trimer, and MA4 of the perimeter MA trimer. **(D)** Residues involved in the interactions between KS3 and restrained MA1 and MA2 of the central MA trimer. **(E)** Heatmap displaying contact frequencies between pairs KS and MA subunits during the ADA.CM.755* simulation. Darker red squares indicate greater contact frequency. Only residue interactions with contact frequencies ≥ 0.2 are plotted. **(F-I)** Representative KS positions relative to the restrained immature MA in the density-guided MD simulations of CH505. **(G)** Residues involved in the interactions between KS1 and restrained MA3 of the central MA trimer. **(H)** Residues involved in the interactions between KS2 of CH505 and restrained MA1 of the central MA trimer and MA4 of the perimeter MA trimer. **(I)** Residues involved in the interactions between the KS3 and restrained MA monomers MA1, MA2, MA3 of the central MA trimer. Residue contacts shown in the figure had contact frequencies ≥ 0.5 during the MD simulations. The KS regions are colored purple, MA is green, residues of the CTs involved in CT-MA interactions are orange, and the residues of MA involved in CT-MA interactions are blue. Residue numbers are in accordance with the HXB2 numbering scheme. **(J)** Heatmap displaying contact frequencies between pairs of KS and MA subunits during the CH505 simulation. Darker red squares indicate greater contact frequency. Only residue interactions with contact frequencies ≥ 0.2 are plotted.

Taken together, the cryo-ET data suggest that in ADA.CM.755* Env-bearing immature VLP, the interaction between MA and the 755* CT exhibits a degree of flexibility, though they retain sufficient interactions to localize Env over an MA trimer. Notably, upon inspection of the region connecting Env to MA, in agreement with the CH505.N197D, a density we interpret as the KS extends from the TM towards the center of the MA trimer (Fig 2C). This lobe of density fits a truncated homology model for the TM and CT with the KS loop positioned towards MA, albeit with splayed rather than bundled TM. More peripherally, a second density reaches up from MA to the inner membrane leaflet, in fact this density connects the pocket on MA which has been shown by cryo-EM to bind the SP2 peptide in mature MA lattices with PIP2 or IP6 in free MA trimers by NMR ^69,70^. Alternatively, this bridging density may reflect a C-terminal segment of the KS loop fragment of LLP2. At the present resolution it is not possible to identify the composition of this bridge of density, however we note it is a robust density feature, which likewise is occupied with strong density in the case of the full-length CH505.N197D Env-CT.

To determine whether the MA lattice positioning beneath Env, observed following focused refinement on the membrane/MA layer, accurately reflects its position in the VLP, we also performed local refinement of the Env ectodomain (described in the accompanying paper) ^71^ and compared the position of each particle between refinement centered on the membrane and ectodomain regions for the ADA.CM particles (Fig S4D) and CH505 particles (Fig S4E). In both datasets, most particles only exhibited small (5-15 Å) shifts between Env and MA, indicating that the majority of Env is close to centered above MA trimers (Fig S4F). The small shift is presumably either the result of flexibility of the ectodomain-membrane connection or slight variability in the MA-CT interaction.

Additionally, we performed local refinement of both the ADA.CM.755* and CH505.N197D inner membrane regions with relaxation to C1 symmetry (Fig S5A-B). The resulting maps were noisier than the C3-symmetry imposed map, however no significant change was observed upon C1-symmetric reconstruction of the immature ADA.CM.755* MA lattice (Fig S5D-E). For CH505.N197D Env, the CT and MA could still be fit into the density (Fig. S5C). Compared to the fit into the C3 map, MA was shifted 4.6 Å from the center of the Env-CT baseplate, but otherwise the reconstructions were in excellent agreement. The agreement of reconstructions with and without symmetry imposed indicate that the trimeric nature of the MA and CT densities were not the result of artifacts at the axis of symmetry during reconstruction.

We conclude that at the resolution of our maps, Env is centered over the MA trimer for both ADA.CM.755* Env and CH505.N197D Env. However, the full-length CH505.N197D CT was resolved directly over a single MA trimer, while the 755* Env interacts with MA in an intact lattice with more variability in specific relative positioning of Env and the MA lattice. These findings point to both specific interactions between the CT and MA as well as a perturbing effect of the full-length tail on the immature MA lattice.

### Molecular dynamics simulations reveal asymmetric engagement of CT and MA trimers

To gain more detailed insight into the Env-CT-MA interaction, MD simulations were performed starting from the positioning mapped out by cryo-ET. Models of the full-length CH505 and ADA.CM.755* Envs were constructed and embedded in membranes that mimic the lipid composition and leaflet asymmetry of HIV-1 particles ^48–50^. Next, we docked an immature MA lattice composed of 4 MA trimers centered underneath the Env-CT using energy minimization and pulling simulations to facilitate insertion of the N-terminal myristoyl-groups of MA into the lower leaflet of the viral membrane (Fig S6). We then created composite cryo-ET maps based on the focused refinements of the membrane and ectodomain layers for each strain and performed all-atom MD simulations in which the peptide was fit to the density (Fig S7). In these simulations, the inter-trimer distances of the immature MA lattice were laterally restrained based on the expectation that it is more constrained in its native context via its C-terminal tethers to the immature CA lattice and the presence of neighboring MA trimers not included in the simulated system.

Based on our cryo-ET density map, we hypothesized that the KS loop may interact with MA near the trimer interface and possibly insert into the center of the MA trimer. Therefore, we analyzed the positioning and residue contacts of the KS in our simulations of first the simpler ADA.CM.755* Env and observed an asymmetric interaction with MA in which the three KS loops occupy different positions above the MA trimer, with one KS indeed inserting into the central cavity of MA and the other two positioned over MA monomers (Figure 3A-E, Table S2). KS-1 was centered over MA monomer 3 (MA-3), mostly forming contacts with helix-2 residues K32, V35, R39, and K40, as well as residue A3. KS-2 sat above MA-1 as well as over the immature lattice interface where it contacted residues G2, S6, V7, and G49 of MA-1 and A5, S9, G10, and G11 of MA-4. KS-3 occupied a central position we suspect accounted for the strong connecting density in our cryo-ET map, making contacts in or near the central axis of the MA trimer where it contacts MA-1 and MA-2. KS-3 interacted with MA1 helix-2 residues E40, L41, R43 of MA-1, G71 which connects the 3_10_-helix and helix-4, as well as E74 and E75 of helix 4. KS-3 also interacted with neighboring MA-2 residues V35, R39, F42, R43, F44, A45, V46, and N47.

A similar pattern of KS-MA interaction was observed in the simulation of the full-length CH505 tail (Figure 3F-J, Table S3). In general, many interacting residues were present in both simulations, with the list of MA residues contacted by the KS motifs in the CH505 simulation being more extensive.

In addition to the density-guided simulations, we ran unbiased all-atom simulations of both strains in parallel as controls to assess whether the conformations obtained in the density-guided simulations are favorable or artificially constrained. In support of the cryo-ET data and the density-guided simulations, we observed a similar pattern of KS positioning and residue contacts in the unbiased simulations, supporting the validity of our cryo-ET-based model (Fig S8, Tables S5 & S6). Comparing the contacts between simulations, a number of MA residues formed contacts with the KS across all simulations (ADA.CM and CH505, density-guided and unbiased) (Table S2-3, S5-6). Notably, mutations at several of these contact residues, such as L13, A45, and L75, have previously been shown to reduce Env incorporation and infectivity ^9,12^. Additionally, L31 formed contacts with the KS in three of the four simulations, supporting the L31E mutation, which has also been shown to affect Env incorporation ^9^. Of the KS residues that interacted with MA, residues 723-736 were consistently observed in contact with MA residues across simulations. Furthermore, we observed more contacts with higher contact frequencies between the KS’s and MA in the CH505 Env compared to the ADA.CM.755*, which indicated that the interaction was more constrained for CH505 than for ADA.CM.755*, consistent with differences in flexibility observed by cryo-ET (Fig 3E, J). The KS-MA residue contacts observed in ADA.CM.755* tend to be a subset of the contacts observed with the CH505 Env (Figure 3E, J) as the full-length tail in the latter system may lead to more tight packing.

In summary, the MD simulations were consistent with the hypothesis that the KS of an Env sitting directly over an MA trimer can facilitate interaction with MA via an asymmetric interaction in which one KS subunit occupies the central MA trimer cavity and the other two interact with other membrane facing residues of MA.

### Molecular dynamics reveals disruption of the MA lattice by the full CT baseplate

The CH505 MD simulations included the full-length tail, thus in addition to the analysis of KS-MA interactions, we examined the role of the LLP regions in mediating interactions with MA (Fig 4A-D, Tables S4, S7). Surprisingly, the LLPs did not form extensive contacts with MA. Most contacts involved residues 2-4 of MA, which are in close proximity to the membrane due to the insertion of the N-terminal myristoyl group, and therefore are likely to be in close contact with the membrane embedded amphipathic LLP helices. In addition, V35, where the V35E substitution was previously shown to eliminate Env incorporation ^9,55^, exhibited a high contact frequency with R846 of the CT-2 protomer (Fig 4C). This contact, as well as contacts with MA residues 2-4, were maintained in the unbiased simulation (Fig S8). The only additional MA residue contacts involving the LLPs in the unbiased simulation was between G10 of MA and R788 of the CT-1 protomer.

**Figure 4.**
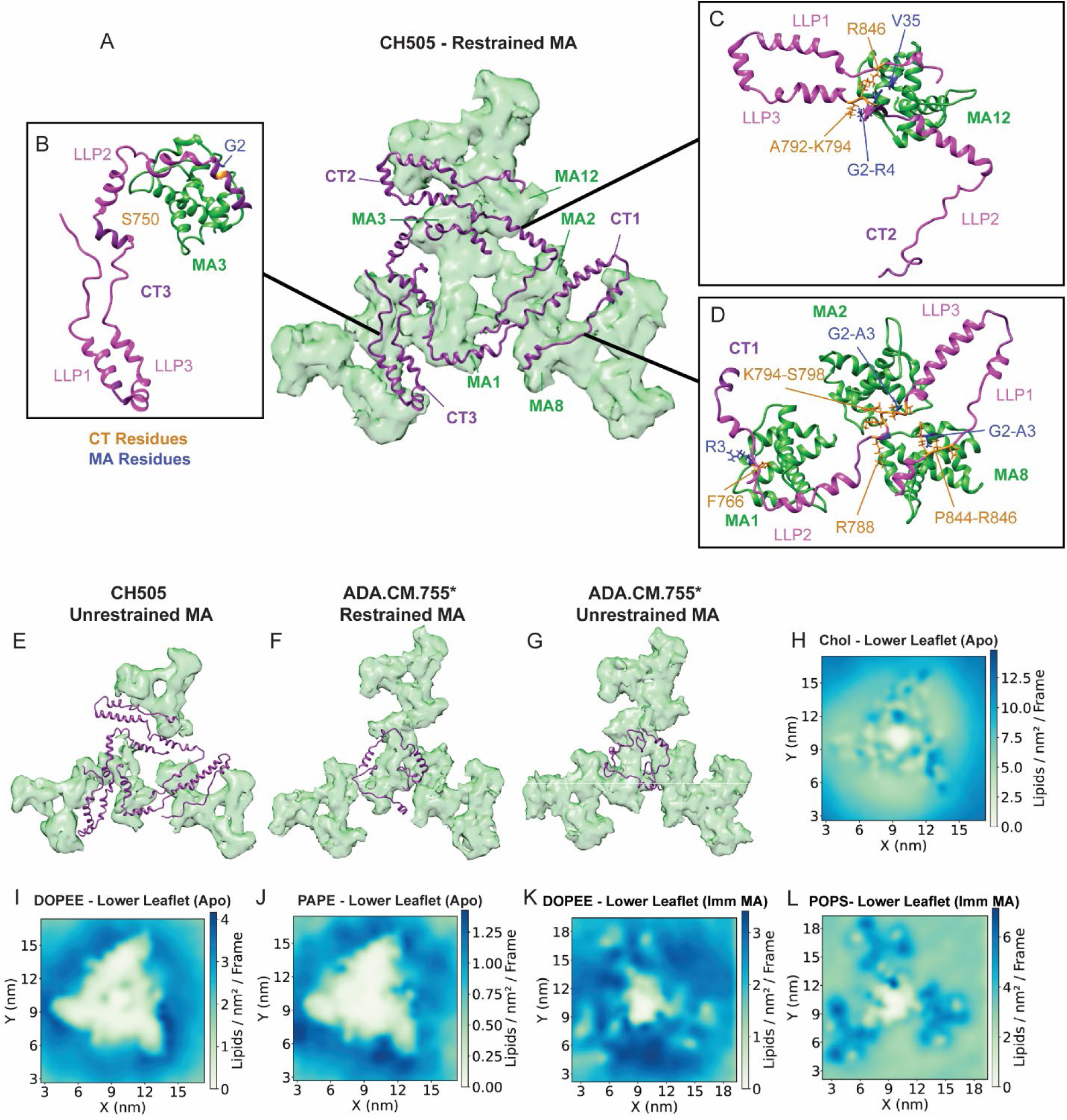
Interactions between the gp41 CT and the immature psg53 Gag MA protein captured from the MD simulations. **(A)** Representative CH505 Env CT positions relative to the restrained immature MA in the density-guided MD simulations. No distortion of MA trimer was observed under the restraints. **(B)** Residues involved in the interactions between the third gp41 CT (CT3) and restrained MA monomer 3 (MA3) of the central MA trimer. **(C)** Residues involved in the interactions between CT2 and restrained MA12 of the perimeter MA trimer. **(D)** Residues involved in the interactions between CT1 and restrained MA monomers MA1 and MA2 of the central trimer and the peripheral MA8 **(E)** Representative gp41 CT positions relative to the unrestrained immature MA in the density-guided MD simulations. The MA trimers were heavily distorted if no restraint was applied. **(E, F)** Representative gp41 CT positions relative to the restrained **(E)** and unrestrained **(G)** ADA.CM.755* density-guided MD simulations. No significant distortion to the MA is present in either simulation of ADA.CM.755* Env. Residue contacts shown in the figure had contact frequencies ≥ 0.5 during simulations. The Env CT regions are colored purple, the LLPs are pink, MA is green, residues of the CT regions involved in CT-MA interactions are orange, and residues of the MA involved in interactions are colored blue. Residue numbers are in accordance with the HXB2 numbering scheme. **(H-J)** Concentrations of CHOL, DOPEE, and PAPE, around the gp41 CT observed in the CG simulations of mature CH505. **(K-L)** Concentrations of DOPEE and POPS around the immature MA protein observed in the CG simulations of ADA.CM.755* with immature MA protein. A color scheme of green – blue was used to illustrate the lowest – highest average density of the lipid molecules in the lower leaflet in the simulations in panels H-L.

We noticed that in the simulations, the CT structure appeared to have exhibited some change compared to its original configuration based on the NMR model, however the LLP1, LLP2, and LLP3 regions remain predominantly helical throughout the simulations (Fig S9). We hypothesized that interaction with the MA lattice may influence the dynamics of the tail. We therefore ran another simulation in which the immature MA lattice distances were not restrained (Fig 4E). In this simulation, we observed substantial distortions of the MA lattice and the individual MA trimer positioned beneath the Env trimer. However, this distortion of the MA lattice did not change the behavior of of the Env-CT baseplate structure (Fig S9C) and the CT did not exhibit large differences in helical probability of the CT between the simulations with or without MA, with minor parts of the LLPs exhibiting slightly less helicity in the presence of MA (Fig S9D-E). To determine whether the distortion of the MA lattice was due to the full-length CT and not instability of the lattice in the unrestrained simulation, we compared the MA lattice in comparable restrained and unrestrained simulations containing the truncated ADA.CM.755* CT. In these cases, we did not observe distortion of the MA lattice (Fig 4F-G). Therefore, we conclude that the full-length CT baseplate structure can induce deformation of the MA lattice via steric incompatibility between the CT-bound and lattice MA forms. The immature MA lattice was observed to be fragmented ^27^, so we interpret the steric incompatibility between CT-bound and lattice MA forms as a feature of the virus that can facilitate specific incorporation of Env while limiting incorporation to low levels. These findings are consistent with the presence of only a single, clearly resolved MA trimer residing beneath the full-length CH505.N197D baseplate in our subtomogram average vs the presence of an immature lattice underneath ADA.CM.755*.

### Influence of CT and MA on lipid distributions and membrane order

The CT of CH505 was observed to embed into the inner leaflet of the HIV-1 model membrane during the MD simulations (Fig 4H-J, S10A, S11), starting from its original location lying below the inner leaflet membrane lipid head groups (Fig S6). Furthermore, we observed enriched concentrations of lipids with phosphatidylethanolamine (PE) headgroups, specifically DOPEE and PAPE surrounding the CT of mature CH505 (Fig 4I-J), while CHOL appeared enriched above the baseplate (Fig 4H, Fig S10B-D).

The specific residues from CH505 CT that interact with lipids are summarized in Figure S11. In particular, eight of the top 20 lipid-interacting residues were found in the LLP3 domain (including W803, V822, L818, L814, L815, L796, L807, and L793). Five were found in the LLP2 region (including C764, I775, F774, A779, and L776), while another five were in the LLP1 domain (including C837, R841, F851, V833, and R838). Based on the variations seen among the protomers, we expect these interactions to be dynamic in nature.

MA also influences lipid distributions. In simulations including the immature MA lattice, lipids consistently interacted with two regions in MA, N-terminal myristoylated segment and a region between K26 and E42 (Fig S12). Specifically PAPE, and POPE concentrated around the trimer (Fig 4K, S13D-E) and POPS concentrated above the MA trimer (Fig 4L). CHOL exhibited more punctate localization near the N-terminal residues of MA (Fig S12, S13).

### Testing residue specific effects on Env incorporation and infectivity

Based on the cryo-ET structures and contact residues in our MD simulations, we identified several residue substitutions that we hypothesized would affect CT-MA interactions. First, we sought to validate a set of previously reported mutations that affect levels of Env incorporation in the NL4-3 strain ^9,11,51,55,66^ using the CH505 HIV-1 molecular clone. Here, changes in Env incorporation were quantified by measuring relative intensity of Env gp41 vs p24 bands in SDS-PAGE (Fig 5A-B, Fig S1B).

**Figure 5.**
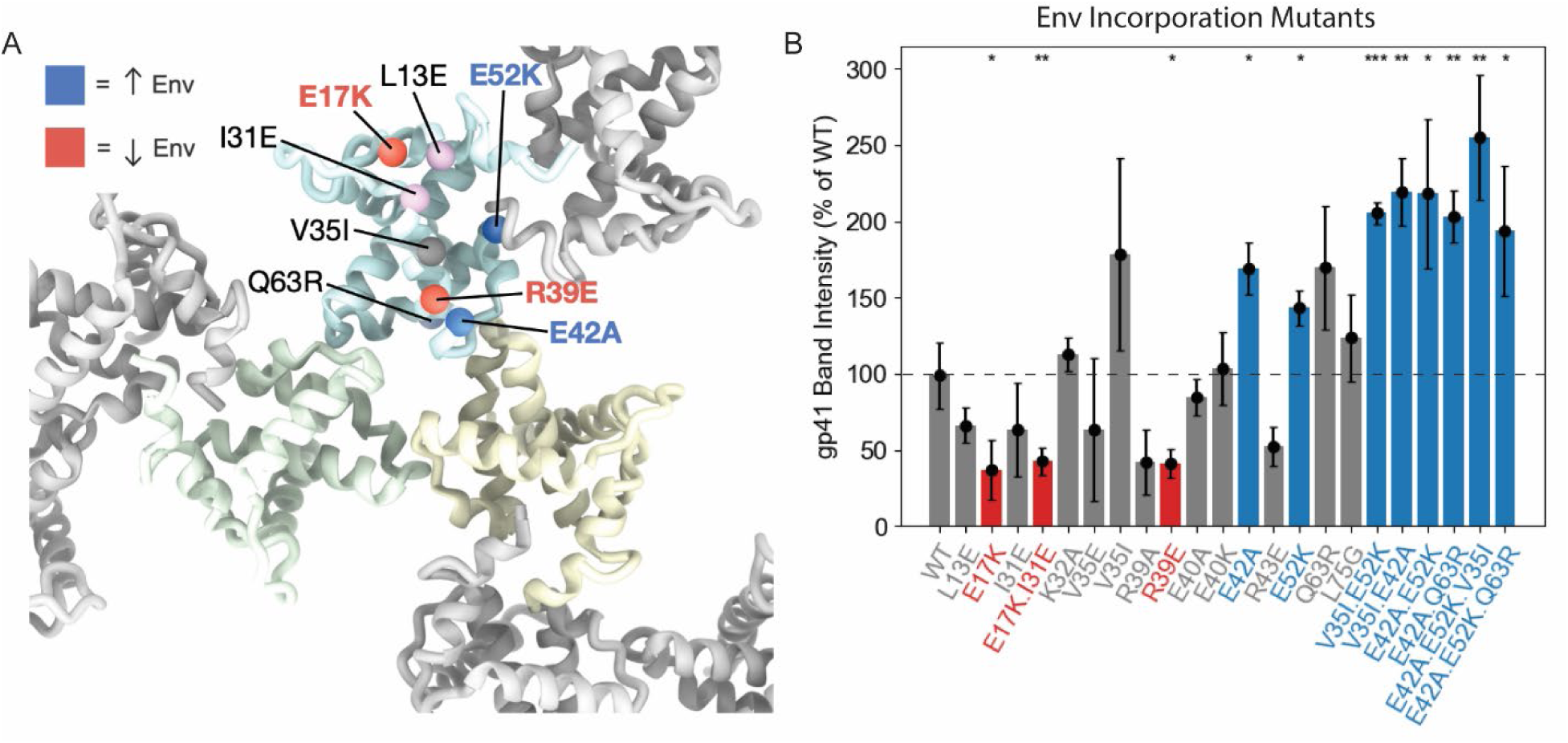
Effect of MA mutants on Env incorporation. **(A)** Model of the immature MA lattice (PDB: 7OVQ) in which the locations of mutants that affected Env incorporation are indicated on a single monomer. Mutants that had a statistically significant effect on Env incorporation in CH505 (P < 0.05) are colored blue or red, for increase or decrease respectively. Mutants nearing statistical significance (P < 0.1) that were shown to have an effect in other studies are colored light blue or light red. Mutants showing effects in other studies that did not have an effect in CH505 are colored grey. **(B)** Histogram depicting the effect of MA mutants on Env incorporation, assessed by comparing gp41 band intensity by SDS page. The height of the bar indicates the average of two replicates. Significance was assessed using one way ANOVA, with asterisks above bars marking comparison to WT. * = P < 0.05, ** = P < 0.01, *** = P < 0.001. Mutants that increased incorporation are colored blue, decreased is colored red.

Consistent with effects previously reported in NL4-3 strain of HIV-1 ^9,11^, MA mutants L13E (P = 0.063) and E17K (P = 0.022) resulted in decreased Env incorporation (Fig 5B), although the P value for L13E was just above the level to indicate significance. We examined mutants I31E (P = 0.081), V35E (P = 0.38), and L75G (P = 0.39) which previously reduced Env incorporation in NL4-3 ^9,11,51,55^. None of these mutants significantly affected Env incorporation in CH505, although I31E exhibited a modest decrease. The E17K.I31E double mutant (P = 0.008) decreased Env incorporation to a level similar to E17K alone (Fig 5B). The mutation Q63R that was previously shown to compensate for the effect of V35E ^55^ resulted in increased Env incorporation in CH505 to a level just above the threshold for significance (P = 0.089) (Fig 5B). Similarly, we observed a modest but not significant increase in V35I (P = 0.17) and a significant increase in E52K (P = 0.025) alone (Fig 5B). Both of these mutations were previously shown to compensate for defects imposed by L13E and C87S mutations, respectively ^10,66,66^.

We also tested the role of residues implicated in CT-MA interactions based on the MD simulations and cryo-ET data. Mutation R39E resulted in a significant decrease in Env incorporation (P = 0.02) and R43E (P = 0.12) resulted in a modest but not significant decrease, while E42A (P = 0.01) resulted increased Env incorporation (Fig 5B). These MA residues are located close to the center of the MA trimer, where we observed a plug of density reaching down in the cryo-ET data and observed interactions with the KS in MD simulations. These mutants follow a common trend, in which placement of glutamate (E) in this region decreases Env incorporation, possibly due to repelling like charges on the KS, which contains several acidic residues. We also tested the mutants K32A (P = 0.50), E40A (P = 0.33) and E40K (P = 0.83) based on interactions observed in the MD simulations, however these did affect Env incorporation (Fig 5B).

Finally, we tested the double and triple mutants of MA V35I.E52K (P = 0.0005), V35I.E42A (P = 0.003), E42A.E52K (P = 0.034), E42A.Q63R (P = 0.003), E42A.E52K.V35I (P = 0.008), E42A.E52K.Q63R (P = 0.047) to determine whether the positive Env incorporation effects could be additive (Fig 5B). All combinations of mutants resulted in statistically significant increases Env incorporation relative to wild-type, and in general showed modest increase in comparison to single mutants. The triple mutant E42A.E52K.V35I showed the greatest increase in Env incorporation compared to all mutants tested, with a ∼2.5x increase compared to WT.

### Maturation-dependent Env clustering is observed by cryo-ET

In many of the tomograms of CH505.N197D VLPs, increased Env clustering was visually apparent in mature vs. immature VLPs (Fig 6A-B, S14A-B). Therefore, Env positions on immature and mature viral particles were identified in the tomograms to quantify differences in distribution across the virus surface depending on maturation state. Only trimers that were retained from particle filtering and classification during subtomogram averaging were included in the clustering analysis. Spatial clustering analysis revealed an increase in the proportion of Env that could be contained within smaller radii in the mature sample (Fig 6C). Complementary pairwise distance analysis confirmed the trends, with a larger fraction of the Env in mature VLPs located within a smaller distance (Fig S14C). However, the trend was not as evident when only plotting nearest neighbor distances (Fig S14D). This was consistent with the Env positioning in raw tomograms, in which pairs or small clusters were often observed in immature VLPs, with larger clusters (>4 Env) more often observed in mature VLPs.

**Figure 6.**
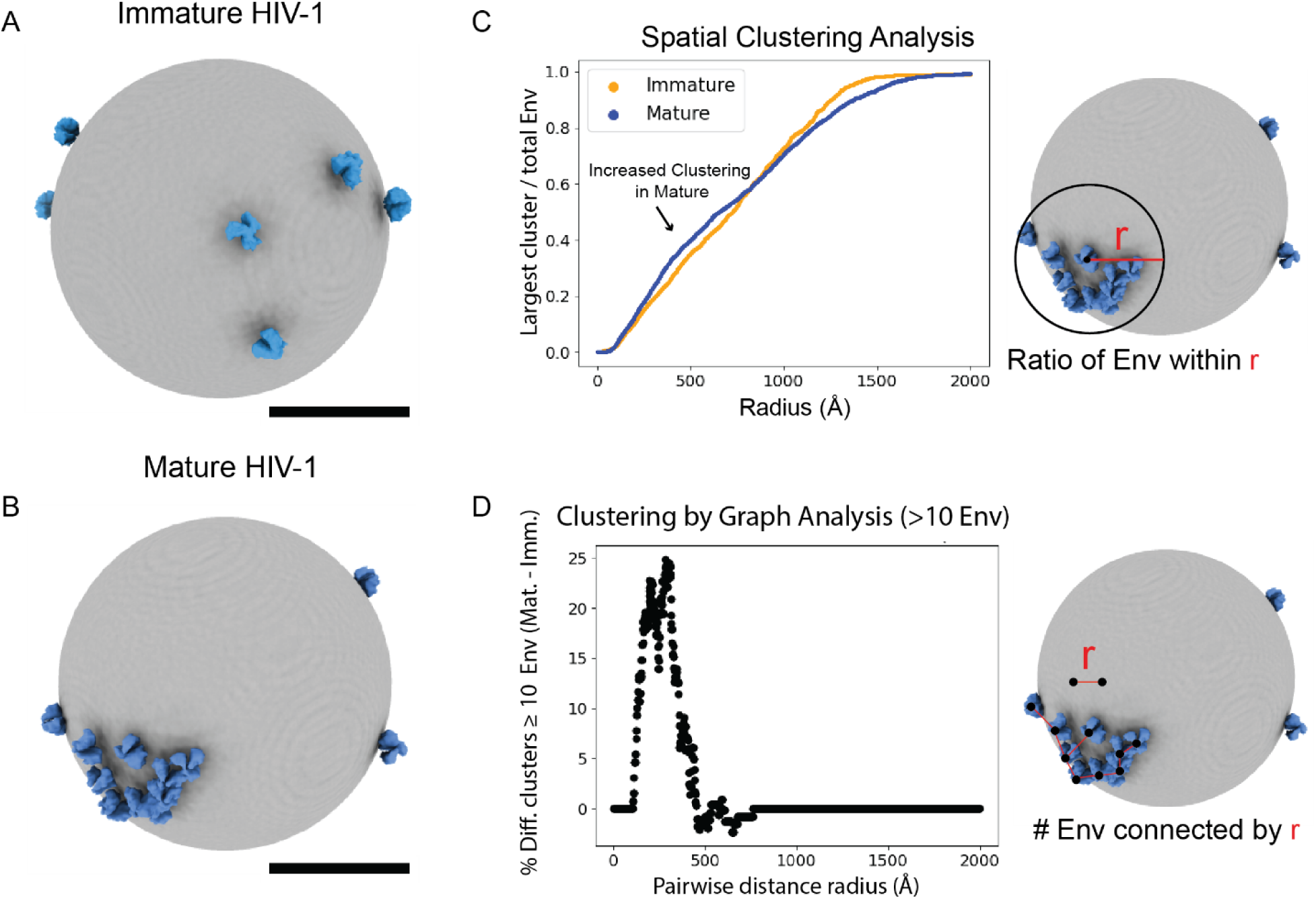
Maturation induces Env clustering. Env positions determined by subtomogram averaging were reprojected back into their location in tomograms of **(A)** an immature CH505 VLP and **(B)** a mature CH505 VLP. Env is depicted as blue density and the viral membrane is grey. **(C)** Spatial clustering analysis in which the average ratio of the largest number of Env trimers that can be contained within a sphere is plotted as a function of radius, as depicted by the diagram on the right. The blue curve corresponds to mature VLPs, the orange immature. **(D)** Env clustering was analyzed by graph analysis, where Env within a given distance threshold (x axis) formed connected nodes on a graph. A connected set of nodes was considered clustered. Difference plot (mature – immature) of percentage of VLPs with a largest cluster of at least 10 Env trimers is plotted against distance threshold. Only VLPs with at least 10 Env trimers were considered in the analysis. At small distances, mature virions displayed greater Env clustering.

The increase in clusters of 4 or more Env in mature VLPs was confirmed using graph analysis (Fig S14E) and was stronger when plotting difference in clusters of 10 or more Env (Fig 6D). What appeared to be a contradictory increase in apparent clustering in immature VLP at distances nearing the size of whole VLP (>700 Å) (Fig S14E), was determined to be the result of greater average Env copy number on immature CH505 VLPs (7.5 per immature VLP vs 5.2 per mature VLP) (Fig S14G-H) and was not present when considering only VLPs with 10 or more Envs (Fig 6D). Thus, the enhancement in Env clustering in mature VLP would likely be even more prominent if both particle samples exhibited the same number of Envs per virion. Cumulatively, our clustering analysis is consistent with previous reports ^37^, and we confirm that Env clustering is facilitated by proteolytic maturation of the Gag lattice.

## Discussion

Here, using cryo-ET, MD simulations, and mutagenesis, we have mapped out the interactions between the HIV-1 Env-CT, the MA subunit of the Gag polyprotein, and biologically realistic membranes in viral particles (Fig 7A-B). Dissection of these interactions and the structural consequences on the MA lattice in assembling immature particles, particularly resulting from the presence of E CT, help to explain the selectivity and restriction of high levels of Env incorporation onto HIV-1 particles.

**Figure 7.**
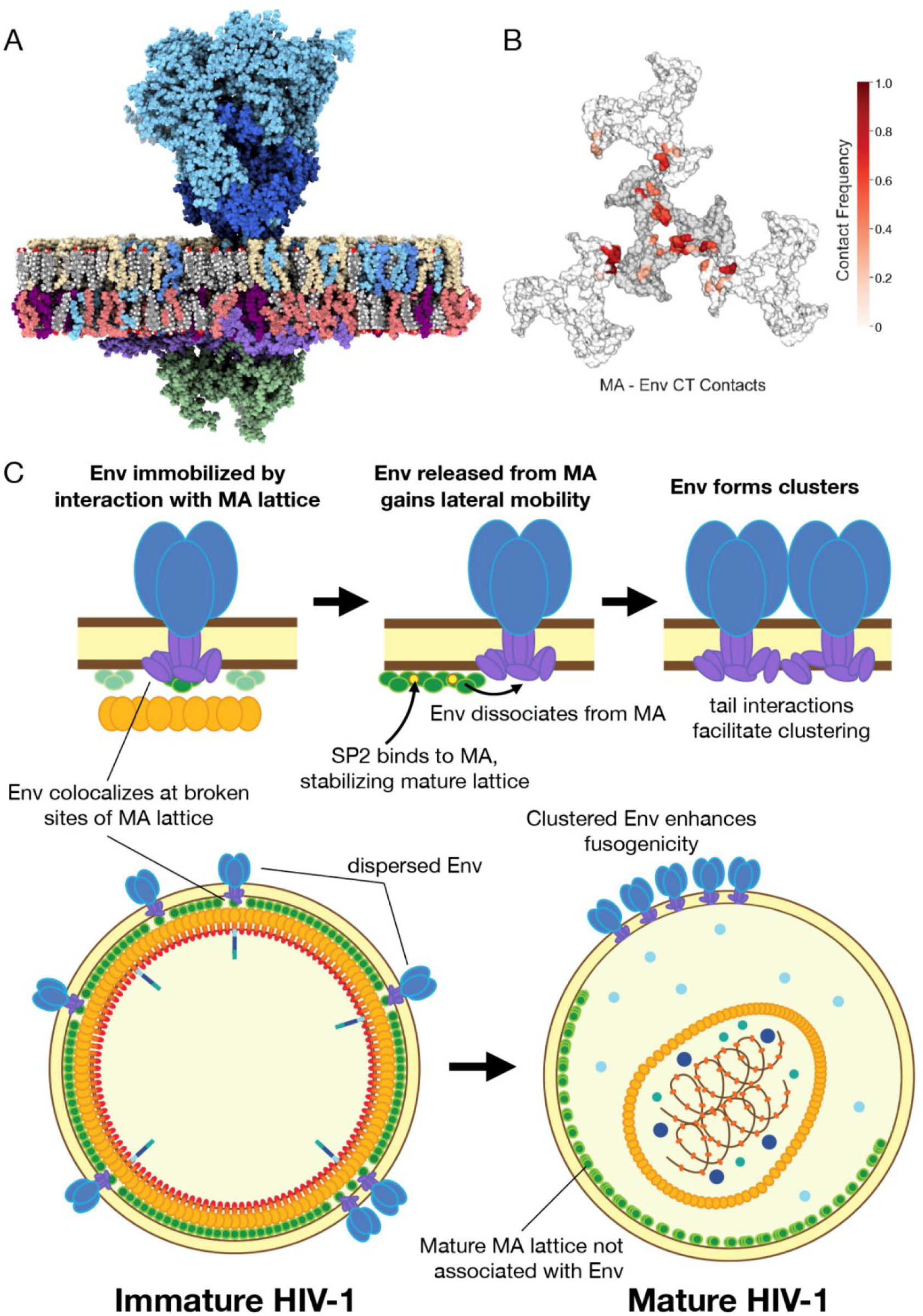
Model of HIV-1 maturation. **(A)** Final conformation of one replicate of the unbiased MD simulation of CH505 Env in complex with immature MA. gp120 is colored cyan, gp41ectodomain is blue, gp41 CT is light purple, MA is green, PSM is yellow, PC lipids are steel blue, POPS is purple, PE lipids are red, CHOL is grey. **(B)** Residues of the immature MA lattice are colored red by contact frequency with the Env CT in the density-guided MD simulations of CH505 in complex with the immature MA. The central MA trimer is highlighted by coloring the residues that did not contact Env darker grey than the adjacent lattice trimers. **(C)** Cartoon model of HIV-1 maturation. Env is in a dispersed pattern on immature VLPs, immobilized through interaction with MA. Env CT interacts with MA by binding directly above MA trimers in areas of fragmented lattice. After MA lattice maturation, the SP2-stabilized mature MA lattice no longer binds Env, and Env becomes mobile on the surface of the virion where it diffuses and forms clusters with other Env trimers in a CT- and Chol-dependent manner. We propose the difference in clustering contributes to difference in Env fusion activity between maturation states.

We observed density bridging from the center of the Env TM/CT to the central hole of the MA trimer which we hypothesize involves the hydrophilic KS region that connects the TM to the amphipathic CT LLPs. This was confirmed by our MD simulations, in which residues 723-736 of the KS made frequent contacts with MA, and one KS even inserted into the central cavity of the MA trimer.

The role of the C-terminal segment spanning LLP1 and LLP3 in particular is complex. In addition to forming limited contacts with MA, the bulk of the full-length CT baseplate appears to compete with and perturb immature MA lattice formation. Indeed, it has previously been observed that the immature MA lattice is less well-ordered than the mature lattice ^72^. Together, these observations lead us to propose that full-length Env-CT has the tendency to incorporate at fractures in the MA lattice. This mode of interaction may serve as a mechanism to limit the number of Env displayed on viral particles, since areas of fragmented lattice are limited on the forming particle and intact lattice could sterically hinder Env binding. We do not observe Env binding to the more condensed, ordered mature MA lattice, which we hypothesize is sterically incompatible with bound Env-CT.

A recent NMR spectroscopy study detected two clusters of residues in the CT that interacted with spin-labeled MA; of note, the constructs used in the NMR study removed significant portions of the KS loop (residues 726-736), while retaining the rest of the tail that forms a baseplate structure ^73^. The first cluster of residues (residues 748-753), in LLP2, is consistent with the positioning of the CT centered directly over a single MA monomer in our model. The second cluster of residues (res 802-806, 817, 826, 837, 850-851, and 856), located at more distal regions in LLP1 and LLP3 is not located above the central MA trimer in our model, indeed the span of the baseplate extends well-beyond the dimensions of a single MA trimer. Instead, in our current model, these residues would be positioned to interact with adjacent MA trimers, if the lattice were intact. Although we did not resolve an intact MA lattice clearly under the full-length CT in the CH505.N197D Env construct, these distal regions of Env-CT may interact with neighboring MA trimers while disrupting the MA lattice to an extent that it is heterogenous and not resolved in our sub-tomogram averaged structures. Additionally, in the NMR study the residues immediately preceding the KS deletion were identified as forming strong interactions, and separately, mutation of E735K/E736K in the study was observed to significantly reduce infectivity ^73^, consistent with our findings that implicate the KS as a motif where key MA-contact residues are concentrated.

Another study, in which GST-tagged CT fragments were used to pull down soluble MA, identified residues 796-812 of the Env-CT LLP3 region as interacting with MA ^51^. In our MD simulations, CT residues 793-799 did interact with N-terminal residues G2, A3, and R4 of MA, but did not make extensive contacts with the bulk of MA. The disposition of these N-terminal residues likely depends heavily on the N-terminal myristoyl group coupling the polypeptide to the viral membrane and likely differ in the virus and intact Env-CT/MA context versus how they would interact in the CT fragment-soluble MA scenario.

To explain the structural basis of previously characterized mutants affecting Env incorporation, these were mapped to the structure of MA (Fig 5A). Many of these overlap with regions that interacted with the Env-CT in our MD simulations (Fig 7B) As noted by others ^12,13^, mutants that increase Env incorporation tend to cluster near the center of the MA trimer, where we observe a plug of density that we attribute to the KS reaching down to fill the central cavity of MA. Several of these residues appear to interact with Env-CT in our simulations in which the MA trimer interface is a site of frequent interaction (Fig 3, 7B). Indeed, mutants that decrease MA trimerization have been shown to negatively affect Env incorporation, while mutants that restore MA trimerization can rescue these defects ^12,13^. Furthermore, we identified several new mutations near the density we observed at the trimer interface, including E42A, which resulted in increased Env incorporation, and R39E which reduced Env incorporation. We identified a triple mutant E42A.E52K.V35I that resulted in increased Env incorporation by over twofold (Fig 5B). This mutant could aid future structural studies or VLP-based vaccines in which increasing copy number of full-length, unmodified Env is desirable. Our findings support the conclusion that the MA trimer interface itself is a crucial site of interaction with the Env-CT.

Collectively, our ultrastructural analysis of immature and mature HIV-1 particles provide support for an integrative model of the interactions between Env and Gag during maturation (Fig 7C). During assembly, Env interacts with MA either at regions of fragmented lattice or induces defects in the MA lattice upon binding, and is dispersed across the immature virion ^39^ (Fig 6A, 7D). This interaction is maintained in the immature VLP, in which we visualize the Env-CT binding directly above a single MA trimer by cryo-ET (Fig 1G-J). Our MD simulations suggest this interaction is facilitated by the KS binding near the MA trimer interface and inserting into the central cavity of the MA trimer (Fig 3), as well as by interactions of the LLPs (Fig 4). We do not observe immature MA lattice under the full-length CT, however, disordered density from neighboring MA trimers was present in our averages and suspect that the LLP domains interact with neighboring MA as observed in our MD simulations (Fig 4).

Upon maturation, sequential cleavage of Gag results in release of the SP2 peptide, which has recently been shown to bind MA and stabilize the mature MA lattice ^28^. We did not observe MA underneath Env in mature particles in either strain, therefore, we propose that the mature MA lattice is incompatible with Env-CT binding, and after maturation, Env is competed away and freed from its interaction with MA. We suspect steric clashes between the CT and tightly packed, well-ordered mature MA lattice are responsible for disruption of the CT-MA interaction following maturation.

Released from the MA lattice, Env is free to diffuse across the viral membrane and form clusters mediated by the Env-CT ^36–39^ (Fig 6, 7C). In the accompanying manuscript ^71^ we report that maturation did not give rise to major differences in Env ectodomain conformation or dynamics between maturation states. The previously reported increased fusion activity of mature particles is thus likely to reflect Env clustering and cooperativity of Env-mediated receptor binding and membrane fusion activities that can occur more frequently in mature particles ^38^. It has recently been shown that the mature MA lattice is not required for infectivity ^74^. Rather, we propose that by restraining Env’s lateral mobility and clustering, the immature MA lattice inhibits Env-mediated fusion, thus preventing premature fusion before a mature capsid has formed to ensure protection and trafficking of the genome to the nucleus in the infected cell.

The function of the mature MA lattice itself remains to be elucidated. One possibility is the tendency for MA to form a more stable and compact lattice in mature virus, may help sequester this membrane-buttressing structural lattice on one face of the virus particle, while Env in mature particles forms its own clusters in largely MA-free regions where the lipid membrane will be more pliable for Env-mediated membrane fusion without associated MA. It has also been speculated that the mature MA lattice may perform a post-fusion function within the infected cell ^28^.

In conclusion, we present a detailed structural model of the interaction between MA and the Env-CT and shed light on the mechanism of assembly and coordination of Env fusion activity with the internal state of Gag. These studies point to possible sites for potential intervention by antiretrovial agents that act by disrupting the Env-CT/MA interactions.

## Limitations of the Study

Due to biosafety concerns, mature ADA.CM.755* and CH505.N197D VLPs were treated with AT-2. In our cryo-ET and MD simulations, VLPs contained a mismatch of Pr55^Gag^ from strain SG3 and Env from either strain CH505.N197D, CH505, or ADA.CM.755*. We note this mismatch could contribute to the incompatibility of the stable immature lattice with the full-length Env-CT. Between collection of mature and immature ADA.CM.755* datasets, the 300 kV Titan Krios used for imaging was upgraded from a Gatan K2 to a Gatan K3 direct electron detector. Therefore, different imaging conditions were used which may have resulted in subtle differences in quality and resolution of the maps. However, this should not affect the main findings that the MA lattice was resolved beneath Env in only the immature map or that the ectodomain displays a similar prefusion conformation in both maturation states. We employed extensive density-guided and unbiased multi-scale (AA and CG) MD simulations to explore the interactions between the HIV-1 Env gp41 KS and CT with the immature MA protein. Similar limitations in the study with AA MD simulations, as provided in the accompanying paper ^71^, are applicable here. As noted in the accompanying paper, we have applied the same three rigorous measures to address these limitations. Similar considerations in CG simulations have also been incorporated in this study to address the limitations of the Martini 3 force field. In the AA MD simulations carried out in this study, we kept the MA trimers laterally restrained to mimic immature HIV-1 particles. Finally, the HIV-1 viral membrane model used in simulations incorporated all dominant lipid types present at >5% in the experimentally determined lipidomic studies. Therefore, the inner leaflet wasn’t populated with the PIP2 and other lipid molecules with compositions of less than 5%.

## Acknowledgements

This work was supported by National Institutes of Health grants R01-AI140868 (KKL) and R01-AI179697 (CAD, SG, KKL), R01-AI186650 (CAD, SG), Duke Center for HIV Structural Biology (U54-AI170752-01), and S10-OD032290.. A portion of this work was also supported by grants from the Gates Foundation (INV-084290 and INV-010646) (KKL). We thank the University of Washington Arnold and Mabel Beckman Cryo-EM Center and the School of Pharmacy Mass Spectrometry Center for data collection time and support. We would also like to thank Max Crispin for providing unpublished data on the glycosylation of CH505 Env. Anju Yadav, Cesar Lopez, and Mingfei Zhao for insights on setting up glycans and the complex membrane in the simulations. The authors acknowledge the support from the Los Alamos National Laboratory Institutional Computing for providing extensive computational resources.

## Author Contributions

Conceptualization, J.T.C., H.N.D., K.K.L. S.G.; Formal Analysis, J.T.C, H.N.D., K.N.L.; Funding Acquisition, K.K.L., S.G., C.A.D., M.B.Z.; Investigation, J.T.C., H.N.D., D.P.L., K.N.L., P.R.J.,K.J.C.; Project Administration, K.K.L., S.G., C.A.D., M.B.Z.; Resources, M.B.Z; Software, J.T.C, H.N.D., K.N.L.; Supervision, K.K.L., S.G., C.A.D., M.B.Z.; Validation, J.T.C., H.N.D.; Visualization, J.T.C, H.N.D., K.N.L.; Writing, J.T.C., H.N.D., K.K.L. S.G.; Writing Review, C.A.D., M.B.Z., D.P.L., P.R.J., V.M.P., K.N.L.

## Declaration of Interests

The authors declare no competing interests.

## STAR Methods

### Key resources table

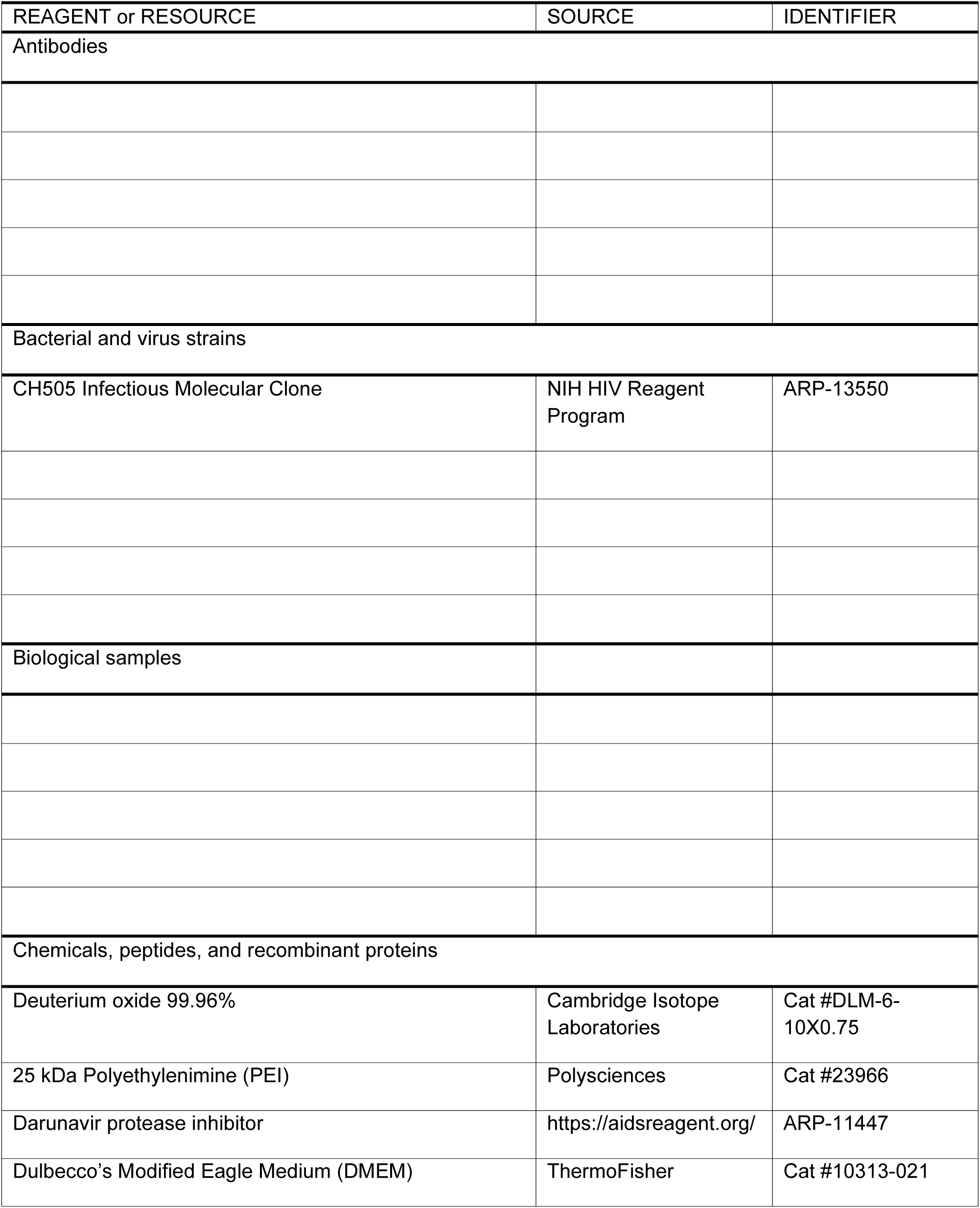
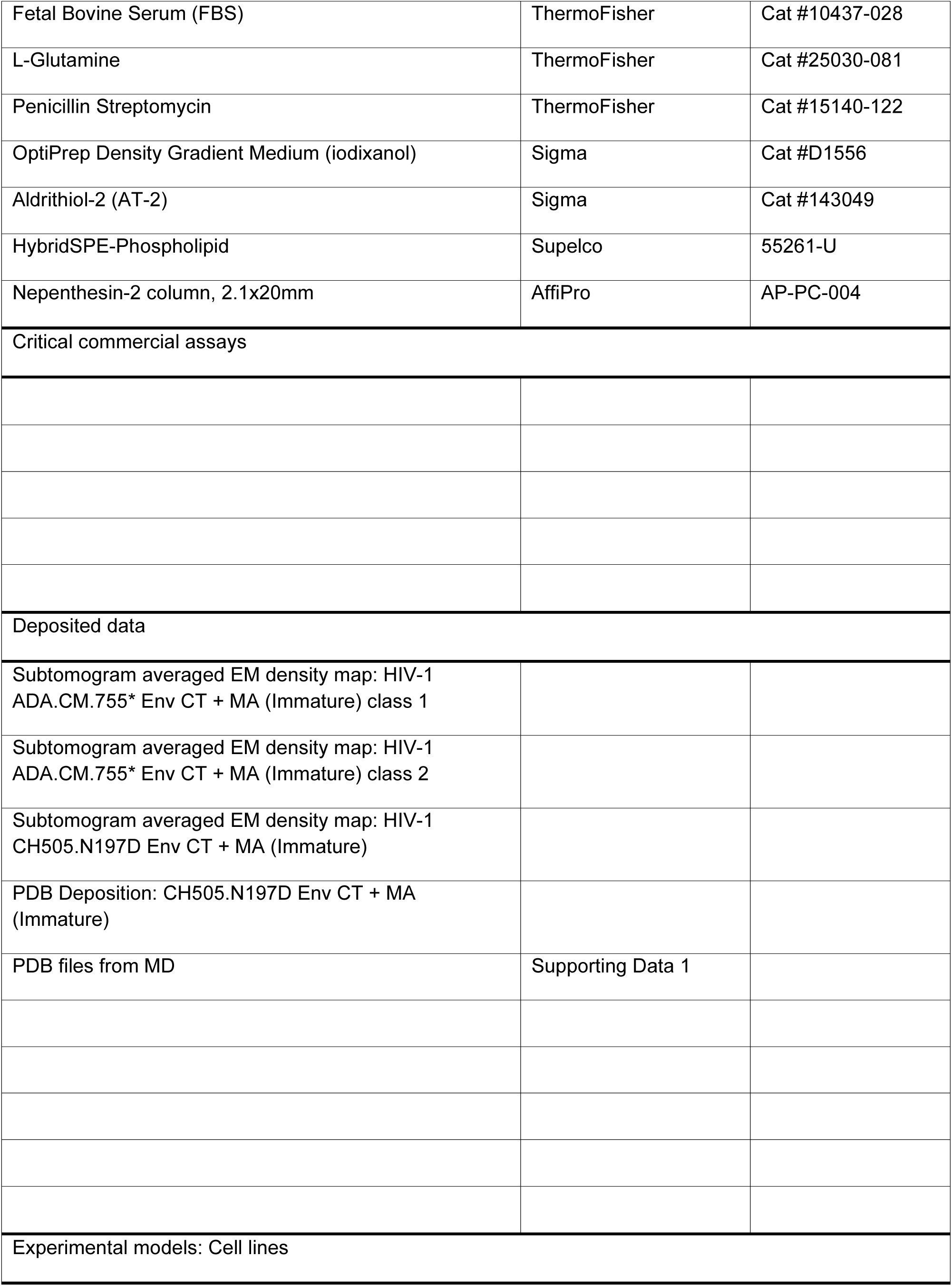
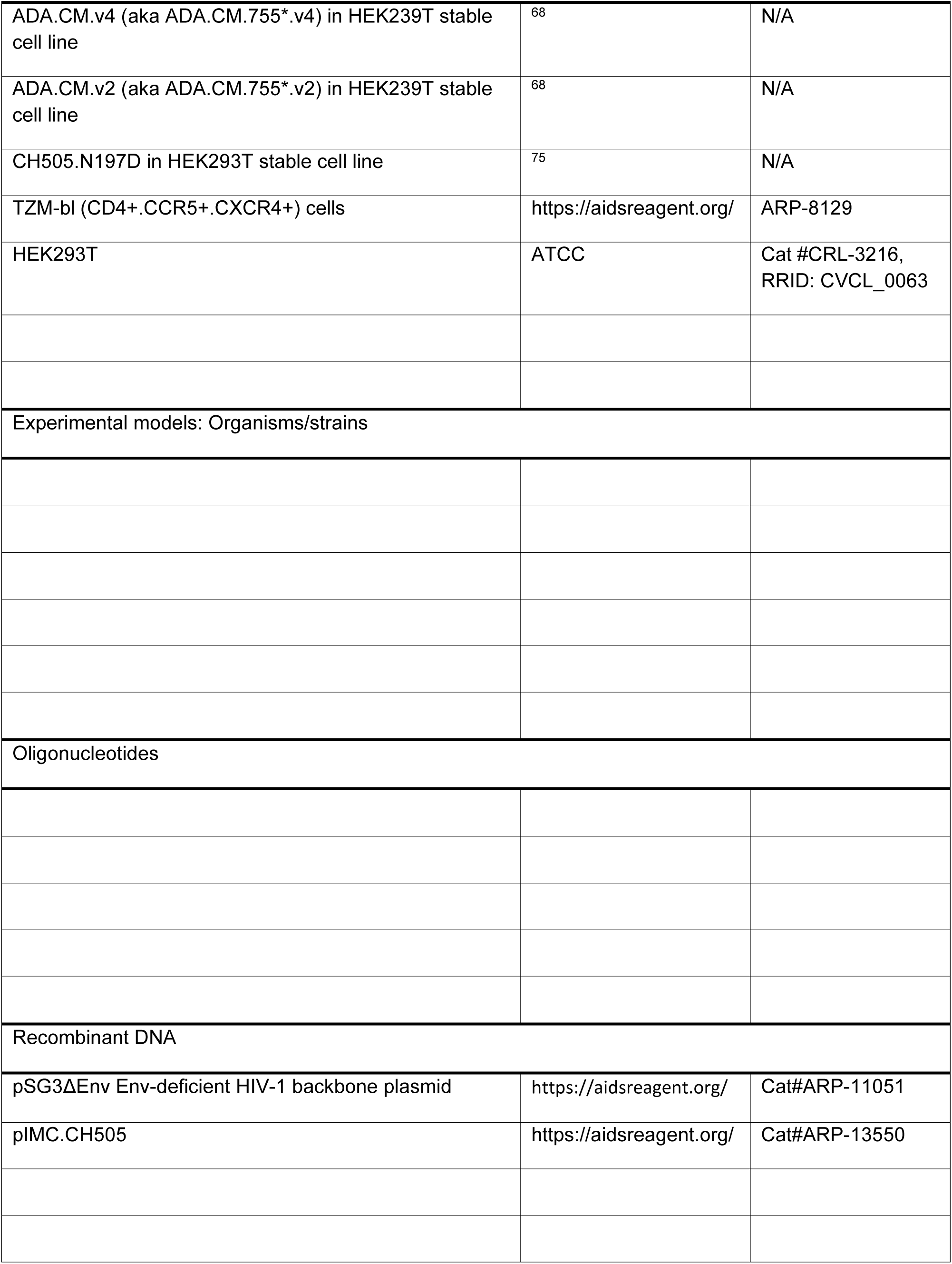
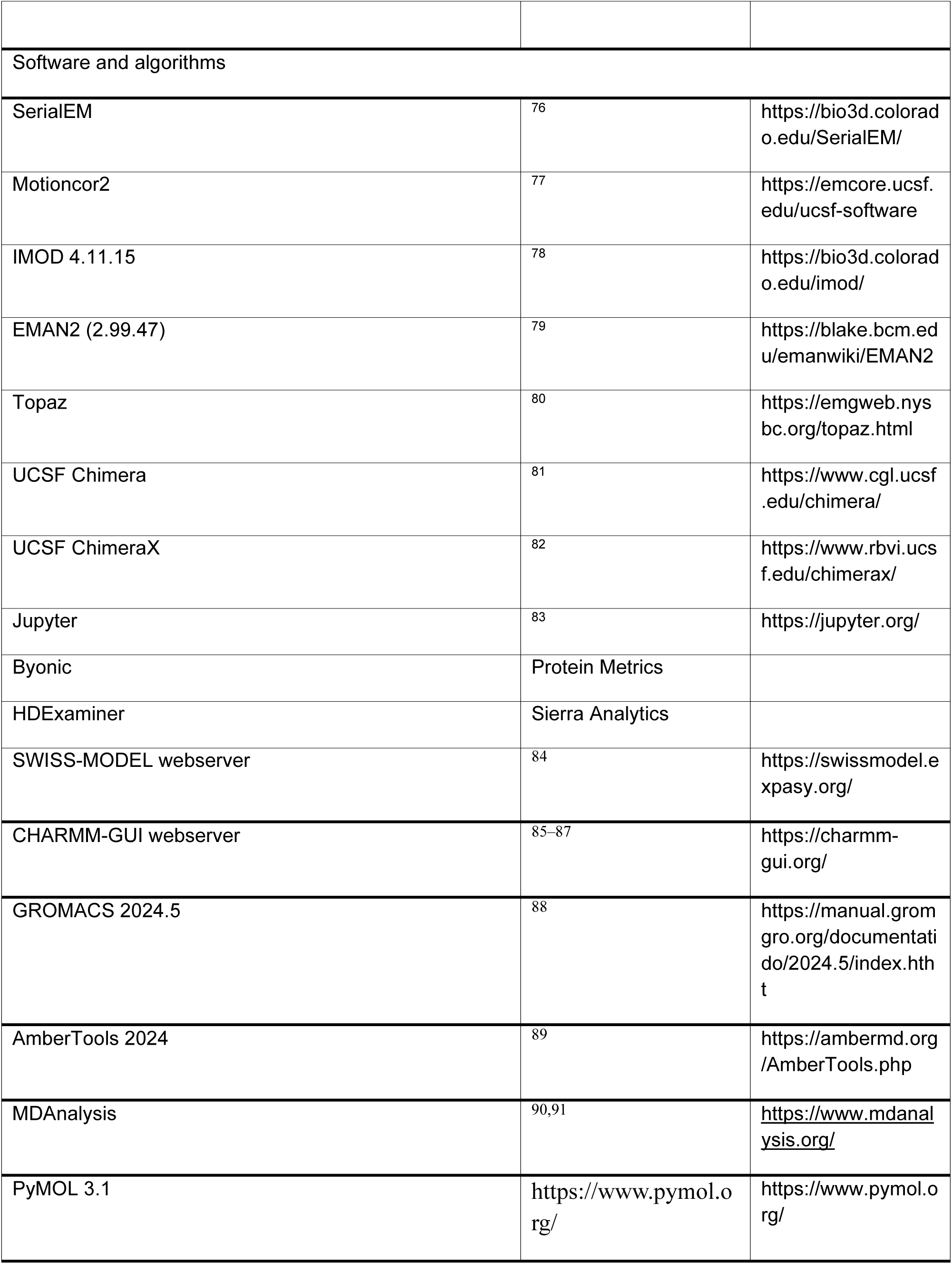
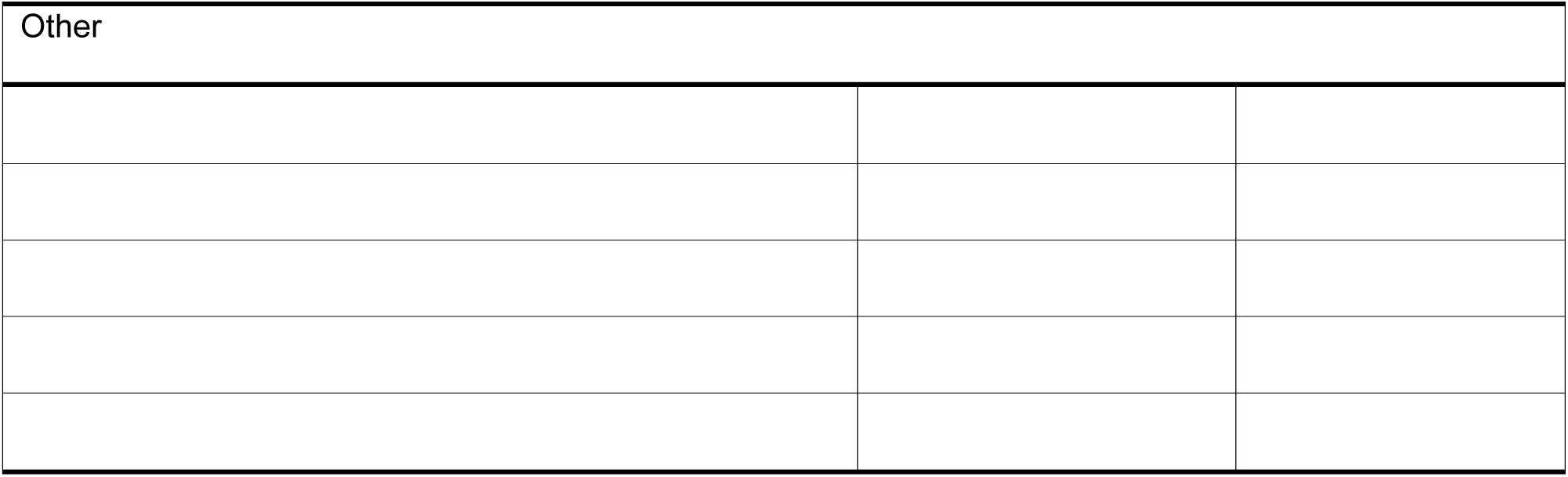

### Experimental Models and Subject Details

#### Cell lines for producing VLPs

Production of the “ADA.CM.v4” HEK293T cell line expressing high levels of HIV Env, was described previously ^68^. Cells were grown in D10 media (DMEM supplemented with 10% heat-inactivated fetal bovine serum (FBS; ThermoFisher Scientific), 20 mM L-glutamine, 100 U/mL penicillin, 100 mg/mL streptomycin, and 2.5 mg/mL puromycin (all media additives from ThermoFisher Scientific) and incubated at 37°C with 5% CO_2_). A stable cell line expressing Env CH505.N197D was generated by transduction of HEK293T cells using the lentiviral vector pLenti-III-HA following the same method as the ADA.CM.v4 cell line and has been described previously (Leaman et al., 2021). The sex of the HEK293T cell line donor was female.

### Method Details

#### VLP expression and purification

Stable cell lines expressing high levels of ADA.CM and CH505.N197D Envs have been previously described ^68,75^. Cells were plated in 15-cm tissue culture dishes in 30 ml media (DMEM containing 10% heat-inactivated fetal bovine serum, 20 mM glutamine, 100 U/ml penicillin, 100 μg/ml streptomycin, and 2.5 μg/ml puromycin) at a cell density of 2.5 x 10^5^/ml and allowed to attach overnight. Next, 30 μg of Env-deficient HIV-1 backbone plasmid pSG3ΔEnv (NIH AIDS Reagent Program) in 900 μl DMEM was mixed with 90 μg 25 kDa PEI transfection reagent (Polysciences) in 900 μl DMEM, incubated for 15 minutes at room temperature, and the PEI/DNA mixture was added to the cells. Immature VLPs were produced by adding the protease inhibitor darunavir (NIH AIDS Reagent Program) to transfected cells at the same time as the DNA and PEI to a final concentration of 2 µM. Supernatants were harvested 3 days after transfection and cleared by centrifugation at 3,000 × *g* for 15 min. VLPs were pelleted at 40,000 × *g* for 1 h and resuspended 50-fold concentrated in PBS. VLPs were separated from cellular debris using iodixanol density gradient centrifugation. Concentrated VLPs were overlaid on a 9.6 to 20.4% iodixanol (Optiprep; Sigma) gradient, formed by layering iodixanol in 1.2% increments, and centrifuged at 200,000 × *g* for 1.5 h at 4°C in an SW41Ti rotor (Beckman). The top 4 ml of the gradient were discarded and then the next 5 ml were collected and pooled. Purified VLPs were brought up to 15 ml with PBS and concentrated to ∼0.2 ml using a 100 kDa MWCO Amicon centrifugal filter (Millipore) spun at 2,000 × *g*. VLPs were inactivated by adding aldrithiol-2 (AT-2) to a final concentration of 2.5 mM and samples were incubated at RT for 2 h. The volume was again increased to 15 ml with PBS and VLPs were concentrated using a 100 kDa MWCO centrifugal filter to 250-fold the concentration in the original transfection supernatant.

#### MA mutant virus production

The infectious molecular clone (IMC) of CH505 was obtained from the NIH AIDS Reagent Program and mutants of the plasmid were generated by Genscript. Viruses were produced by transfection of HEK293T cells. Cells were plated in 6-well tissue culture plates in 2.5 ml media/well (DMEM containing 10% heat-inactivated fetal bovine serum, 20 mM glutamine, 100 U/ml penicillin, and 100 μg/ml streptomycin) at a cell density of 2 x 10^5^/ml and allowed to attach overnight. A mixture of 2 μg of IMC plasmid DNA and 6 μg 25 kDa PEI transfection reagent (Polysciences) in 100 μl DMEM was incubated for 15 minutes at room temperature, after which the PEI/DNA mixture was added to the cells. Supernatants were harvested 3 days after transfection and cellular debris was removed using a 0.45 µm syringe filter.

#### SDS-PAGE Western blot to measure Env incorporation

CH505 IMC viruses in cell culture supernatants were pelleted by centrifugation at 22,000 × *g* for 45 min and resuspended 50-fold concentrated in PBS. P24 content was determined by p24 antigen capture assay using a kit provided by the AIDS and Cancer Virus Program, Frederick National Laboratory for Cancer Research, and following the instructions provided. Virus samples, normalized for p24 content, were incubated in Laemmli Buffer (Bio-Rad) containing 50 mM dithiothreitol (DTT) for 5 min at 100°C before loading on 8-16% Tris-glycine gels (ThermoFisher). Samples were loaded in duplicate and wild-type virus was included on every gel for comparison, and two identical gels were loaded for separate Western blotting. Gels were electrophoresed at RT in running buffer (25 mM Tris, 192 mM Glycine, 0.1% SDS, pH 8.3) at 150 V for 1 h. Proteins were transferred to a PVDF membrane in transfer buffer (25 mM Tris base, 200 mM glycine, 10% methanol) at 20 V for 1 h using an XCell II Blot Module (Invitrogen), and membranes were blocked in 5% non-fat dry milk. Blots were probed overnight at 4°C with either sheep anti-p24 polyclonal Ab (Aalto) or an anti-Env Ab cocktail (1 μg/ml each 10E8, 7B2, 98-6, 2 μg/ml each HGN194, 2G12, F105) in PBS containing 5% NFDM and 0.05% Tween-20. After washing, membranes were probed for 30 min at room temperature with either donkey anti-sheep-IgG or goat anti-human-IgG HRP-conjugated Abs (Jackson), and peroxidase activity was assayed using ECL Western Blotting Substrate (Pierce). Blots were imaged using a Chemidoc instrument (BioRad) and bands were quantified using ImageLab software (BioRad). The intensity of the gp41 band on the anti-Env blot was calculated for each mutant relative to that of wild-type virus as a measure of Env incorporation. The amount of p24 loaded relative to wild-type virus was similarly calculated and the amount of Env incorporated was normalized to account for differences in p24 loaded.

#### Preparation of Samples for CryoET and Tilt-series Acquisition

Either Quantifoil R 2/2 300 or 400 mesh copper or Quantifoil R1.2/1.3 200 or 400 mesh copper grids (Electron Microscopy Sciences) were glow discharged for 30s at 25mA using an EMS100X plasma cleaner. 3 uL of purified virus was applied to each grid, which were plunge frozen on a Vitrobot Mark IV (FEI) at 100% humidity, blot force 0, and blot times of 3, 3.5 or 4 s.

For mature ADA.CM.755* VLPs, sample preparation and data collection was performed as described previously ^63^. For darunavir treated ADA.CM.755* VLPs, grids were imaged on a 300 kV Titan Krios (ThermoFisher) with a Gatan K3 direct electron detector equipped with an energy filter (slit width 20 eV). Tilt series were collected using SerialEM ^76^ using a dose-symmetric tilt scheme ranging from either +/- 48 or +/- 60 degrees with a step size of 3 and a total dose of either ∼75 or ∼94 e/Å^2^, respectively. Tilt movies were collected as 0.3 s exposures divided into 6 frames in super-resolution mode at 64kx magnification corresponding to a super-resolution pixel size of 0.6932 Å/pixel. A total of 691 tilt series were collected over 6 sessions.

For mature and darunavir treated CH505.N197D VLPs, grids were imaged on a 200 kV Glacios (ThermoFisher) equipped with Gatan K3 direct electron detector. Tilt series were collected using SerialEM with a dose-symmetric tilt scheme between +/- 60 degrees with a step size of 3 and a total dose of ∼95-100 e/Å^2^. Tilt movies were collected as 0.55 s exposures divided into 5 frames in counting mode at 24kx magnification corresponding to a pixel size of 1.86 Å/pixel. For mature CH505.N197D VLPs, a total of 327 tilt series were collected over 4 sessions. For darunavir treated CH505.N197D VLPs, a total of 248 tilt series were collected over 3 sessions.

For imaging of detergent-stripped VLPs, Triton X-100 and VLPs were to a final concentration of 0.1% incubated on ice for 15 minutes, and run over a HiPPR detergent removal spin column (Thermofisher) before plunge frozen under similar conditions to the other samples.

#### Preprocessing and Tomogram Reconstruction

Prior to tomogram reconstruction, movies were motion corrected using MotionCor2 ^77^ and combined into ordered tilt-series stacks using the IMOD ^78^ newstack command. From this point onward, all subsequent processing was performed in EMAN2 (version 2.99.47) unless otherwise specified ^92^. Tilt series stacks were imported into EMAN2, the tomograms were aligned and reconstructed, and CTF estimation was performed. Some tomograms were denoised using Topaz prior to particle picking ^80^. For all samples, particles were picked manually either in EMAN2 boxer or in IMOD slicer and imported into EMAN2.

#### Tomogram Segmentation and Visualization

Tomogram visualization for figures was prepared using the IMOD slicer tool. For automatic segmentation of tomograms, the neural-network based segmentation tools in EMAN2 were used. Segmentations were visualized with ChimeraX ^82^.

#### Immature (darunavir) ADA.CM.755* Subtomogram Averaging

The same initial model, low-pass filtered to 50 Å was used as a reference for all datasets. After preprocessing, particles were extracted at 4X binning and an initial refinement was performed with C3 symmetry, with a resolution limit of 25 Å. Then, a local refinement was performed with C1 symmetry and the output of this was input into GMM classification with 6 classes. Two classes were selected based on particle orientation and map quality and another refinement was performed with C3 symmetry and a 10 Å resolution limit. Masks were created to include only the ectodomain or the membrane, and local refinements were performed focusing on each region. For the ectodomain, processing is described in a parallel study ^71^.

For the membrane layer, GMM classification was performed to separate the data into 4 classes. One class was discarded because it did not clearly show density for an MA trimer, and the three classes with MA density were combined and a local refinement was performed with C3 symmetry. Particles were re-extracted and re-centered over MA at a binning of 2, and local refinement was performed with C3 symmetry and automasking. The processing pipeline is described in Fig S3.

#### Mature CH505.N197D Subtomogram Averaging

After preprocessing, particles were re-extracted at 4X binning and an initial refinement was performed with C3 symmetry and a 25 Å resolution limit. From here, a local refinement was performed relaxing symmetry to C1 and this was used as an input for GMM classification with five classes. Two classes were selected and combined based on manual inspection of particle orientation and map quality. Next, multireference refinement was performed with 2 references, one prefusion Env and one resembling postfusion Env. The class containing prefusion Env particles was selected and a C3 refinement was performed with a cylindrical mask, limiting resolution to 20 Å. The processing pipeline is described in Fig S2D.

#### Immature (darunavir) CH505.N197D Subtomogram Averaging

The same steps were performed for immature CH505.N197D Env, except GMM classification was performed using only two classes. After separating and refining prefusion Env particles, 3D classification was performed with five classes using a cylindrical mask. Four classes were selected and combined that appeared to have a central MA trimer. From here, both a C3 refinement and a C1 refinement were performed with cylindrical masking and 20 Å limit. The C3 refinement was used for comparison with the mature ectodomain, and local refinement with a mask of the membrane was continued from the C1. This refinement was continued further with C3 symmetry, reducing the resolution limit to 15 Å. Next, particles were re-extracted and recentered over MA and duplicate particles were excluded. Local refinement was performed of these particles with C3 symmetry and a 15 Å limit. Finally, particles were re-extracted with binning of 2 and a final local refinement was performed with C3 symmetry using a tight mask including only the Env-CT baseplate and MA features. The processing pipeline is described in Fig S2A.

#### Env Clustering Analysis

Only particles that were used in final C3 reconstructions of the prefusion ectodomain were used for clustering analysis. Particle coordinates were obtained from their positions after final refinements and clustering analysis was performed using custom python scripts. To analyze “clustering by radius” the maximum fraction of Env that could be contained within an increasing radius of x was calculated within each VLP and the average across all VLPs for each radius was plotted. To analyze “clustering by pairwise distance”, a cluster is defined by constructing a for each VLP in which each Env corresponds to a node and edges connect Env within a given pairwise distance. Thus, maximum cluster size for each VLP was defined as the largest set of connected nodes at a given distance threshold.

#### Simulations of HIV-1 Envelope Proteins in Complex with Immature Matrix Proteins

We used the SWISS-MODELLER homology modelling webserver ^84^ to model the full all atom (AA) structures of the ADA.CM.755* and CH505 HIV-1 envelope proteins starting from their amino acid sequences. We needed to model the gp120, gp41 ectodomain (ED), gp41 TM and gp41 CT domains separately in SWISS-MODELLER ^84^ before connecting them to obtain the full structures of the envelopes. We modeled the glycosylation states of the gp120 and gp41 ED of the ADA.CM.755* and CH505 HIV-1 envelope proteins based on published and unpublished experimental glycosylation profiles of the trimer envelope ^93^, in accordance with Table S1, using the CHARMM-GUI webserver ^85,87^. The palmitoylation motif (CYSP) was attached to residues C764 and C843 in the gp41 CT of CH505. The residues were numbered in accordance with the HXB2 numbering scheme of gp160 (https://www.hiv.lanl.gov/content/sequence/LOCATE/locate.html). The structure of the immature HIV-1 matrix (MA) protein was taken from the 7OVQ PDB ^72^, with only the central trimer and its three neighboring trimers kept for our simulations. The myristoylation motif (GLYM) was attached to residue G2 in each of the MA monomer in the immature MA protein. We used the CHARMM-GUI webserver ^85–87,94^ to prepare our simulation systems of ADA.CM.755* and CH505 in complex with the immature MA protein. The Envs were embedded in an asymmetric membrane composition mimicking the membrane lipid composition of HIV-1 virion ^49,95^. The outer leaflet was composed of 48% cholesterol (CHOL), 38% palmitoyl sphingomyelin (PSM), 9% dipalmitoylphosphatidylcholine (DPPC), and 5% palmitoyl-oleoyl-phosphatidylcholine (POPC). The inner leaflet was composed of 42% CHOL, 3% DPPC, 2% POPC, 4% palmitoyl-arachidonoyl phosphatidylethanolamine (PAPE), 4% palmitoyl-oleoyl phosphatidylethanolamine (POPE), 14% dioleoylphosphatidylethanolamine (DOPEE), 14% dipalmitoylphosphatidylethanolamine (DPPEE), and 17% palmitoyl-oleoyl-phosphatidylserine (POPS). The immature MA protein was initially placed 20 Å away from the gp41 CT of the envelope proteins and the membrane inner leaflet (Fig S6). The CHARMM36m force field parameter set ^96^ was used for our AA simulations. We performed pulling simulations to insert the myristoylation motifs (GLYM) of the immature MA protein into the inner leaflet of the membrane (Fig S6). The immature MA protein was iteratively pulled upwards towards the inner membrane leaflet by 5 Å at a time before energy minimization was carried out on the whole envelope-matrix-membrane complex to refine any possible clashes between the MA protein and the gp41 CT or the membrane. The pulling simulations were stopped after the myristoylation motifs were properly inserted into the inner leaflet of the membrane (Fig S9). The complexes of HIV-1 Env and immature MA proteins, with the myristoylation motifs inserted in the membrane, were solvated in 0.15 M NaCl using the GROMACS 2024.5 ^88^ simulation package. Overall, the simulation system sizes for our simulation systems of immature Envs with MA incorporated were ∼1 million atoms, with the box dimensions of 207.5 Å × 207.5 Å × 240.0 Å. The total number of lipid molecules in the upper leaflet was 864 molecules, while the total number of lipid molecules in the lower leaflet was 776 molecules in the HIV-1 membrane.

We employed the GROMACS 2024.5 ^88^ simulation package to carry out our simulations of Env proteins in complex with immature MA. We used the LINCS algorithm ^97^ to restrain all bonds containing hydrogen atoms and applied the periodic boundary conditions to our simulation systems. We used the velocity rescaling algorithm ^98^ to keep our simulation temperature constant at 310 K with a friction coefficient of 1.0 ps^-1^, and used the stochastic cell rescaling algorithm ^99^ with semi-isotropic coupling to keep pressure constant at 1.0 bar. We set the pressure coupling constant to 5 ps and the compressibility to 4.5 × 10^-5^ bar^-1^. We first performed energy minimization on our simulation systems for a maximum of 5000 steps using the steepest-descent algorithm. At this stage, we applied position restraints on the backbone atoms with a force constant of 4000 kJ.mol^-1^.nm^-2^, side chain atoms with a force constant of 2000 kJ.mol^-1^.nm^-2^, and, on the lipid and dihedral angles, with a force constant of 1000 kJ.mol^-1^.nm^-2^. We then equilibrated our simulation systems using the constant number, volume, and temperature (NVT) ensemble for a total of 375,000 steps, using a time step of 1 fs. During this stage, we gradually reduced the force constants for position restraints from 4000 to 2000 to 1000 kJ.mol^-1^.nm^-2^ for backbone atoms, from 2000 to 1000 to 500 kJ.mol^-1^.nm^-2^ for side chain atoms, from 1000 to 400 to 200 kJ.mol^-1^.nm^-2^ for dihedral angles, and from 1000 to 400 kJ.mol^-1^.nm^-2^ for lipids, after every 125,000 steps. Next, we equilibrated our simulation systems using the constant number, pressure, and temperature (NPT) ensemble for a total of 750,000 steps, using a time step of 2 fs. After the equilibration stage, we continued with two different sets of simulation setups for the immature MA proteins: one where they were completely unrestrained, and the other where the backbone atoms of the immature MA proteins were positionally restrained in the xy directions using a force constant of 1000 kJ.mol^-1^.nm^-2^. In particular, similar energies of 2,376.6 ± 78.4 kJ/mol and 2,514.8 ± 96.5 kJ/mol were applied to restraint the inter-trimer distances of the immature MA lattice in the restraint simulations of ADA.CM.755* and CH505, respectively. We performed one short 25-ns conventional MD simulation and then four different unbiased MD simulations of up to 1 µs each replica for each of the four simulation systems (including ADA.CM.755* and CH505 in complex with restraint and unrestraint MA proteins).

Density-guided simulations were also carried out after the equilibration stage on the Env proteins with restraint immature MA to explore the interactions between the gp41 CT, including the KS, of ADA.CM.755* and CH505 and the immature MA protein. Since the populated regions in the experimental cryo-ET densities did not cover the whole complexes, we only fit the gp120, gp41 ED (including MPER), gp41 TM in the inner leaflet, gp41 CT (consisting of the LLPs), and the immature MA proteins (excluding their C-terminal helices) of the ADA.CM.755* in complex with immature MA to the experimental cryo-ET density (Fig S7A). For the CH505 complex, we also did not fit the MPER and the gp41 TM to the experimental cryo-ET density (Fig S7C). We used the cross-correlation setting for the similarity measures, with atomic contribution to the experimental density set to be proportional to their masses. We set the density-guided force constant to 10^6^. Furthermore, we set the Gaussian transform spreading width to 0.2 nm and the Gaussian transform spreading range in multiples of width to 4. We set to normalize our densities and used adaptive force scaling, with an adaptive force scaling time constant set to 50 ps, to get much gentler fit in our density-guided simulations. The density-guided simulations for ADA.CM.755* and CH505 in complex with restraint immature MA were carried out in four replicas for each system, of up to 100 ns per replica. Afterwards, unbiased MD simulations were performed starting from the final checkpoints of the density-guided simulation replicas for up to 1 µs.

After the AA simulations, we used the GROMACS 2024.5 ^88^ simulation software suite to process the simulation trajectories and the MDAnalysis ^90,91^ python package to determine the important residue contacts between the gp41 CT (including the KS and LLPs) with the immature MA protein from the density-guided plus unbiased MD simulations and unbiased MD simulations of Env proteins in complex with restrained immature MA using a contact definition of ≤ 4.5 Å between any heavy atoms (Tables S2-S10). We also determined the differences in MA-MA interactions between the restrained and unrestrained simulations for both ADA.CM.755* and CH505 (Tables S11-S12).

Based on the experimental cryo-ET density of ADA.CM.755* in complex with immature MA and our density-guided simulations, we observed that the N- and C-terminals of gp120 of the envelope protein could be inserted in the membrane. Therefore, we extended the N-terminal of gp120 of ADA.CM.755* by adding the GARAEN amino acid sequence before the LWVTV amino acid sequence and the C-terminal of gp120 of ADA.CM.755* by adding the RRVVQREKR amino acid sequence after the VATKAK sequence using a combination of AlphaFold3 ^100^ and SWISS-MODELLER homology modelling webserver ^84^. Four 1µs unbiased production simulations were performed on the simulation system of ADA.CM.755* with extended gp120 N- and C-terminals in complex with restraint immature MA proteins.

The CG simulations of mature CH505 (without extended gp120 termini) and immature ADA.CM.755* Env with extended gp120 termini in complex with immature MA protein in the HIV-1 membranes, with four simulation replicas for each system, were set up to explore the lipid dynamics around the Env gp41 CT and MA protein, starting from the final AA conformations. The mature CH505 conformations were prepared by removing the restraint MA protein from the final unbiased simulation frames of CH505 with immature MA protein. It should be noted that reducing the immature MA protein reduced the simulation box dimensions to 207.5 Å × 207.5 Å × 207.5 Å. For the AA protein and membrane components of mature CH505 (without extended gp120 termini) and immature ADA.CM.755* with extended gp120 termini in complex with immature MA protein in the HIV-1 membranes, the models were converted to their CG representations ^101^ using a combination of martinize2 ^102^ for the protein components, with modifications to accommodate for the myristoylation and palmitoylation motifs in the mapping files ^103^, and backward ^104^ python script for the lipid components. No glycans were backmapped due to the absence of their force field parameters in MARTINI 3 ^105^. The CG protein components were built with the side chain corrections (-scfix) ^106^ and elastic network (-elastic) ^107^. We set the elastic bond force constant to 1500 kJ.mol^-1^.nm^-2^ (-ef 1500), with the lower and upper bound of elastic bond cutoff set to 0 and 1.2 nm (-el 0 -eu 1.2). The bond decay factor and power were set to 0 and 1.0 (-ea 0 -ep 1). The CG protein and membrane were solvated in 0.15 M NaCl solution using insane ^108^, with the box dimensions kept identical to the AA simulations (of 207.5 Å × 207.5 Å × 207.5 Å for mature CH505 and 207.5 Å × 207.5 Å × 240.0 Å for immature ADA.CM.755* with extended gp120 termini and MA protein incorporated). The MARTINI 3 force field parameter sets ^105^ were employed for the CG simulations to explore the lipid dynamics around the gp41 CT and immature MA protein. The CG simulations were performed using GROMACS 2024.5 ^88^.

The CG simulation systems of HIV-1 Env in the HIV-1 membrane were subjected to initial energy minimization, followed by equilibration in the constant number, volume, temperature (NVT) ensemble and then in the constant number, volume, pressure (NPT) ensemble. After the energy minimization and equilibration, one short 100 ns conventional MD (cMD) simulation was performed on each simulation system, before four independent 10 µs production cMD simulations were carried out on each simulation system to explore the lipid dynamics on membrane embedded gp41. We applied periodic boundary conditions to our simulation systems and used a timestep of 10 fs for the cMD simulations. During the equilibration stage, we gradually increased the timestep from 2 fs to 5 fs to 10 fs. Since we only wanted to explore the lipid dynamics in the CG simulations, we applied position restraints on the protein components using a force constant of 250 kJ.mol^-1^.nm^-2^ during the cMD simulations. We calculated the electrostatic interactions using the reaction field method ^109^ and Verlet cutoff scheme ^110^ with a cutoff distance of 1.1 nm for long-range interactions. We kept the temperature constant at 310 K using the velocity rescaling thermostat ^98^ with a friction coefficient of 1.0 ps^-1^. Meanwhile, we kept the pressure constant at 1.0 bar using the stochastic cell rescaling ^99^ with semi-isotropic coupling, with the pressure coupling constant set to 5 ps and compressibility set to 3×10^-4^ bar^-1^.

At the end of the CG simulations, we employed the GROMACS 2024.5 ^88^ simulation software suite to process the simulation trajectories and MDAnalysis ^90,91^ *python* package to explore the lipid dynamics captured during the CG simulations. We used a contact definition of 10 Å to identify neighboring lipid molecules to the gp41 CT and immature MA proteins during the CG simulations of mature CH505 and immature ADA.CM.755* with extended gp120 termini and MA protein incorporated embedded in the HIV-1 membrane, respectively.

#### Quantification and Statistical Analysis

Representative images were chosen from cryo-ET analysis for the figures. Statistical analysis of Env incorporation was performed using one way ANOVA using the scipy.stats module in Python 3.

## Notes

### Competing Interest Statement

The authors have declared no competing interest.

