## Supplemental figures and tables for "Reconstructing a Missing Link of HIV-1 Assembly: HIV-1 Envelope-Matrix Interactions in a Native Viral Context"

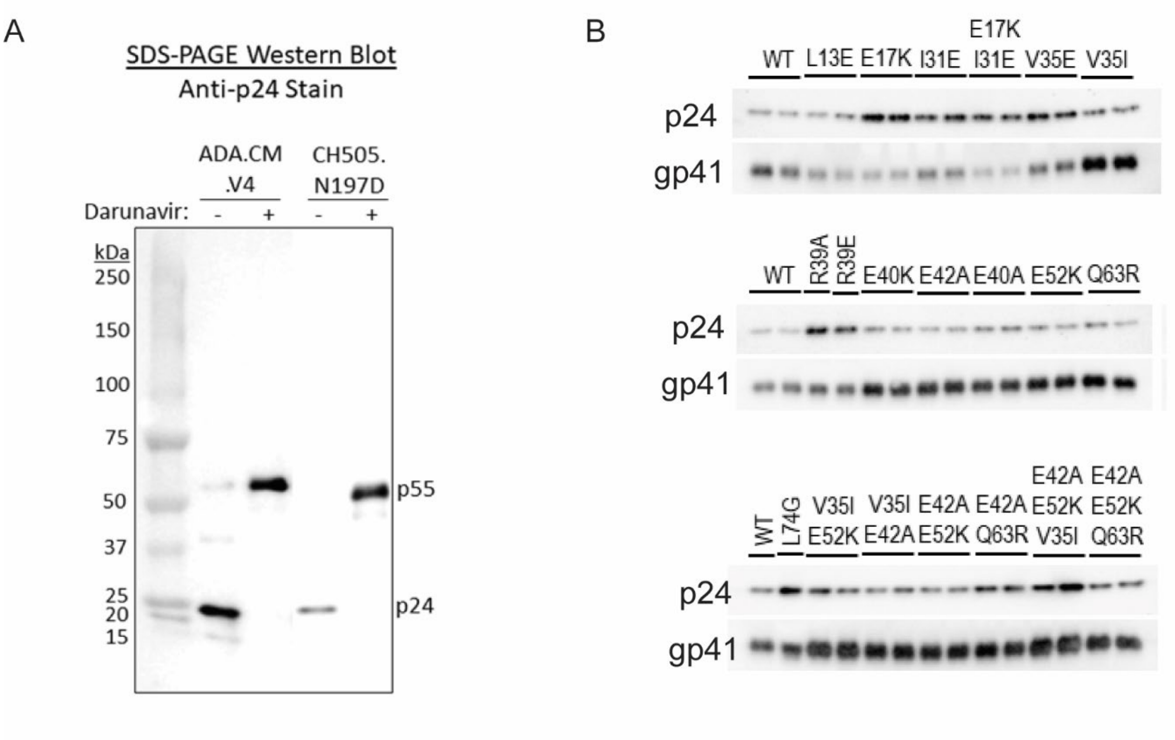


**Figure S1. Western Blots of HIV-1 VLPs.** **(A)** Western blot against p24 (Gag CA) of mature (untreated) and immature (darunavir-treated) VLP preps, confirming the presence of $PR55^{Gag}$ in immature preps and p24 Gag in mature preps. **(B)** Western blots against p24 and gp41, used to calculate Env incorporation levels plotted in Fig 5B.

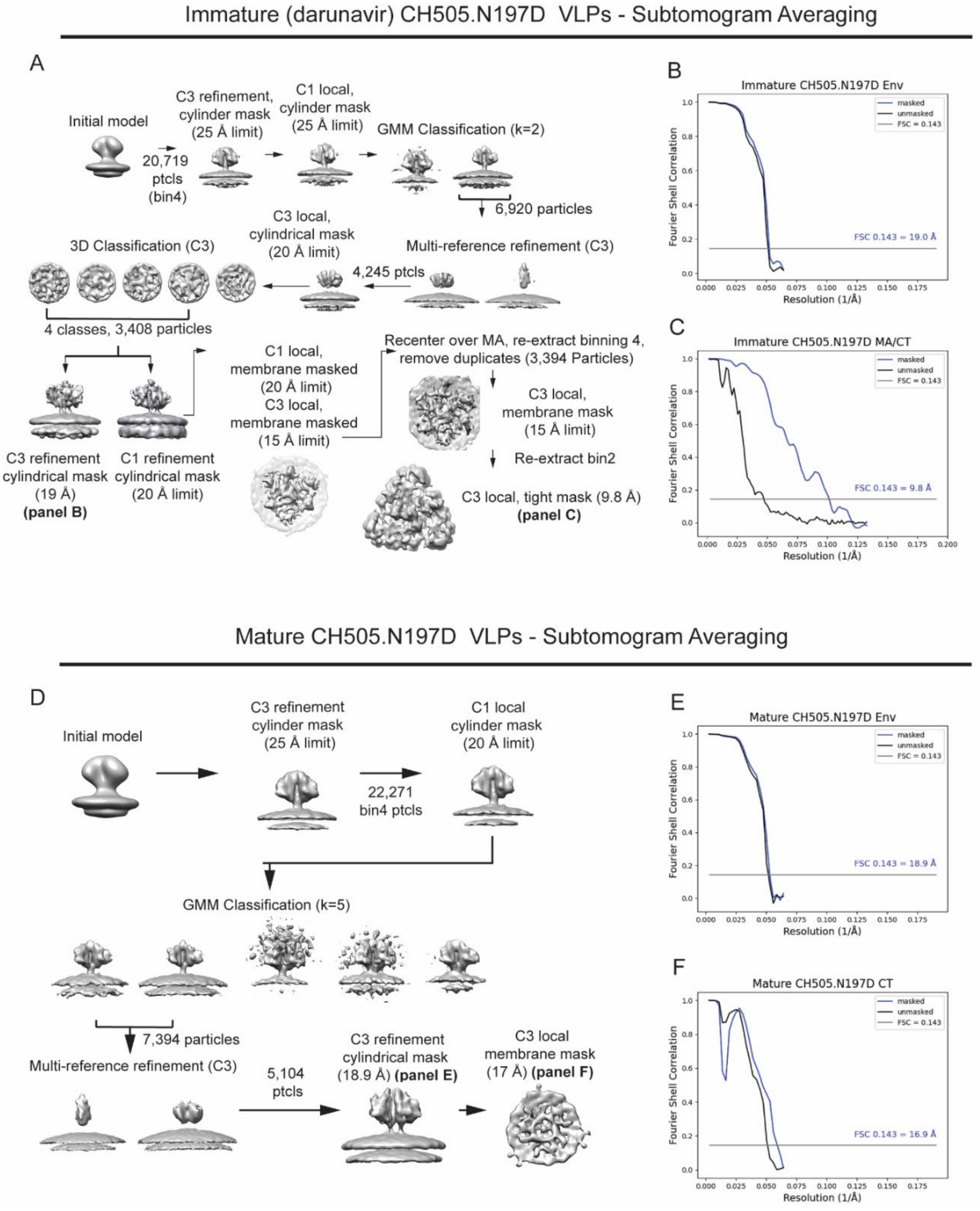


**Figure S2. CH505.N197D subtomogram averaging. (A)** Subtomogram averaging pipeline for the immature CH505 VLPs, as described in the methods. **(B)** Fourier Shell Correlation (FSC) for the immature CH505 VLP Env global refinement including the ectodomain, membranes, and MA layer with C3 symmetry. Using FSC = 0.143 criterion, we report 19.0 Å resolution. **(C)** FSC for the focused refinement of the immature CH505 VLP TM-CT/MA interaction. Using FSC = 0.143 criterion, we report 9.8 Å resolution. **(D)** Subtomogram averaging pipeline for the mature CH505 VLPs, as described in the methods. **(E)** Fourier Shell Correlation (FSC) for the mature CH505 VLP Env global refinement including the ectodomain and membranes with C3 symmetry. Using FSC = 0.143 criterion, we report 18.9 Å resolution. **(F)** FSC for the focused refinement of the Env TM-CT on mature CH505 VLPs. Using FSC = 0.143 criterion, we report 16.9 Å resolution. Throughout panels B, C, E and F the blue curve represents the masked FSC and the black curve represents the FSC calculated with unmasked maps.

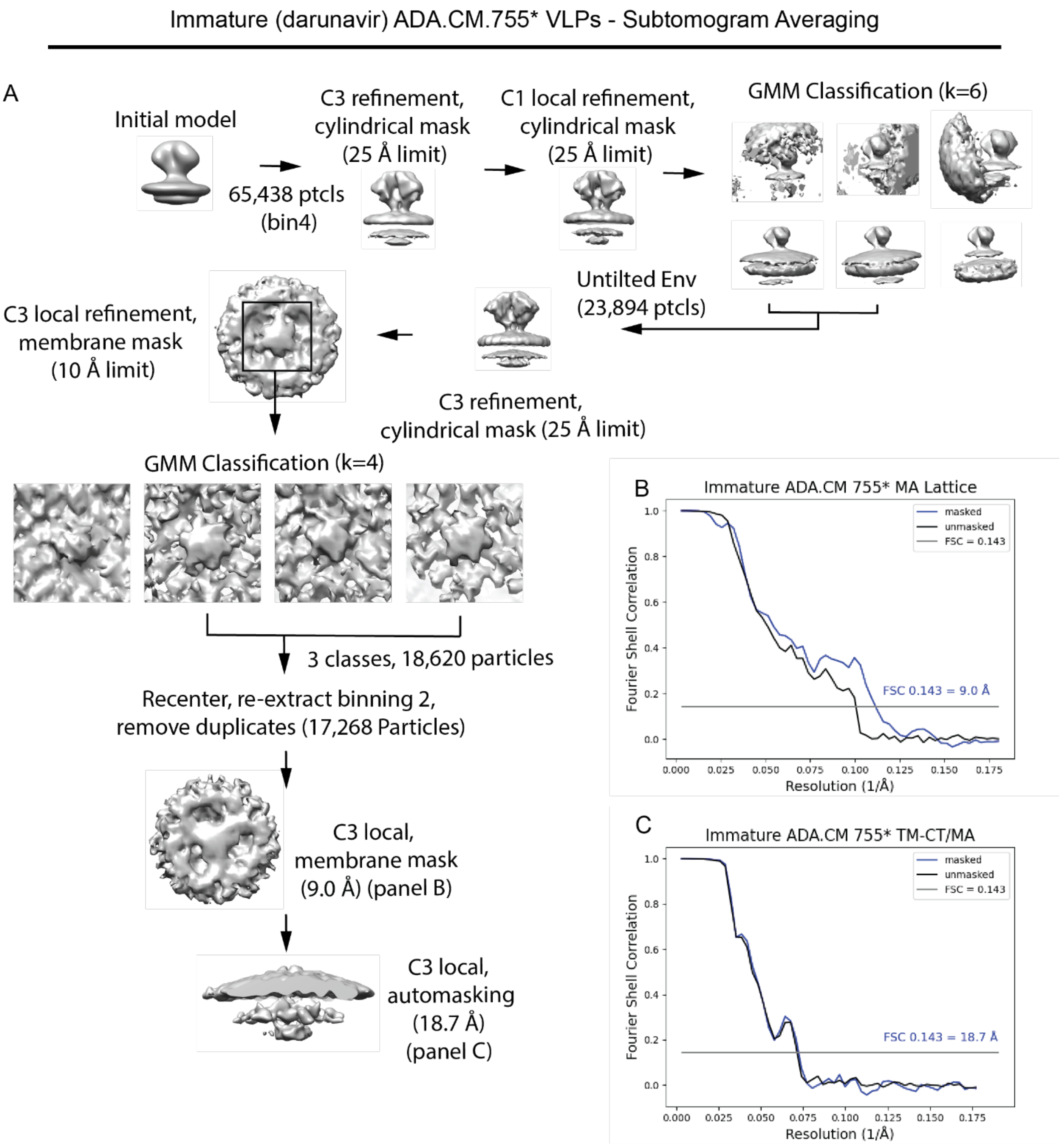


**Figure S3. Immature ADA.CM.755* VLP subtomogram averaging. (A)** Subtomogram averaging pipeline for the Env TM-CT/MA interaction on immature ADA.CM.755* VLPs, and immature MA lattice, as described in the methods. **(B)** Fourier Shell Correlation (FSC) for the focused refinement of the immature ADA.CM.755* MA lattice. Using FSC = 0.143 criterion, we report 9.0 Å resolution. **(C)** FSC for the focused refinement of the ADA.CM.755* TM-CT/MA interaction. Using FSC = 0.143 criterion, we report 18.7 Å resolution. Throughout panels B-

DC, the blue curve represents the masked FSC and the black curve represents the FSC calculated with unmasked maps.

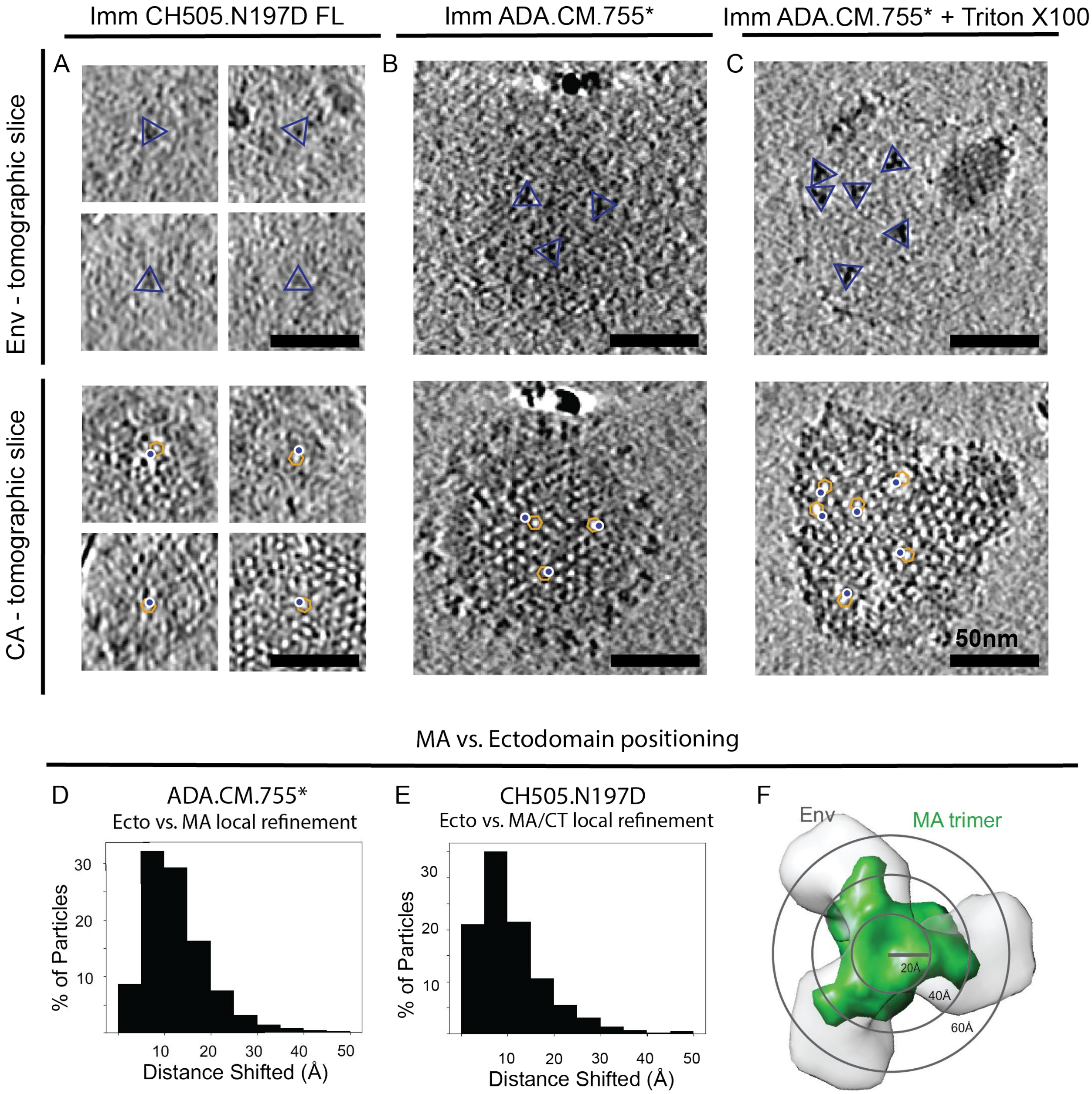


**Figure S4. Positioning of Env relative to CA lattice in cryo-electron tomograms.** In panels A-C, representative 10 nm thick tomographic slices are shown. Images on the top row correspond to Z-slices containing Env, indicated by blue triangles. Images on the bottom row correspond to Z-slices further into the virion, in which the PR55$^{Gag}$ CA lattice is visible. CA hexamers are annotated with orange hexagons, with the positions of Env in the above slices indicated by blue circles. In all three samples, including immature CH505.N197D VLPs **(A)**, immature ADA.CM.755* VLPs **(B)**, and Triton X-100 treated ADA.CM.755* VLPs **(C)** Env appears located over the sides of the CA lattice. Scale bars 50nm. **(D, E)** Histogram plotting distance shifted between corresponding subtomograms between refinements focused on the Env ectodomain and the underlying MA layer in ADA.CM.755* VLPs **(D)** and CH505.N197D VLPs **(E)**. In panel **(F)** concentric circles represent 20 Å radii from the center of the particles for scale. MA trimer density is colored green, Env ectodomain is colored transparent grey.

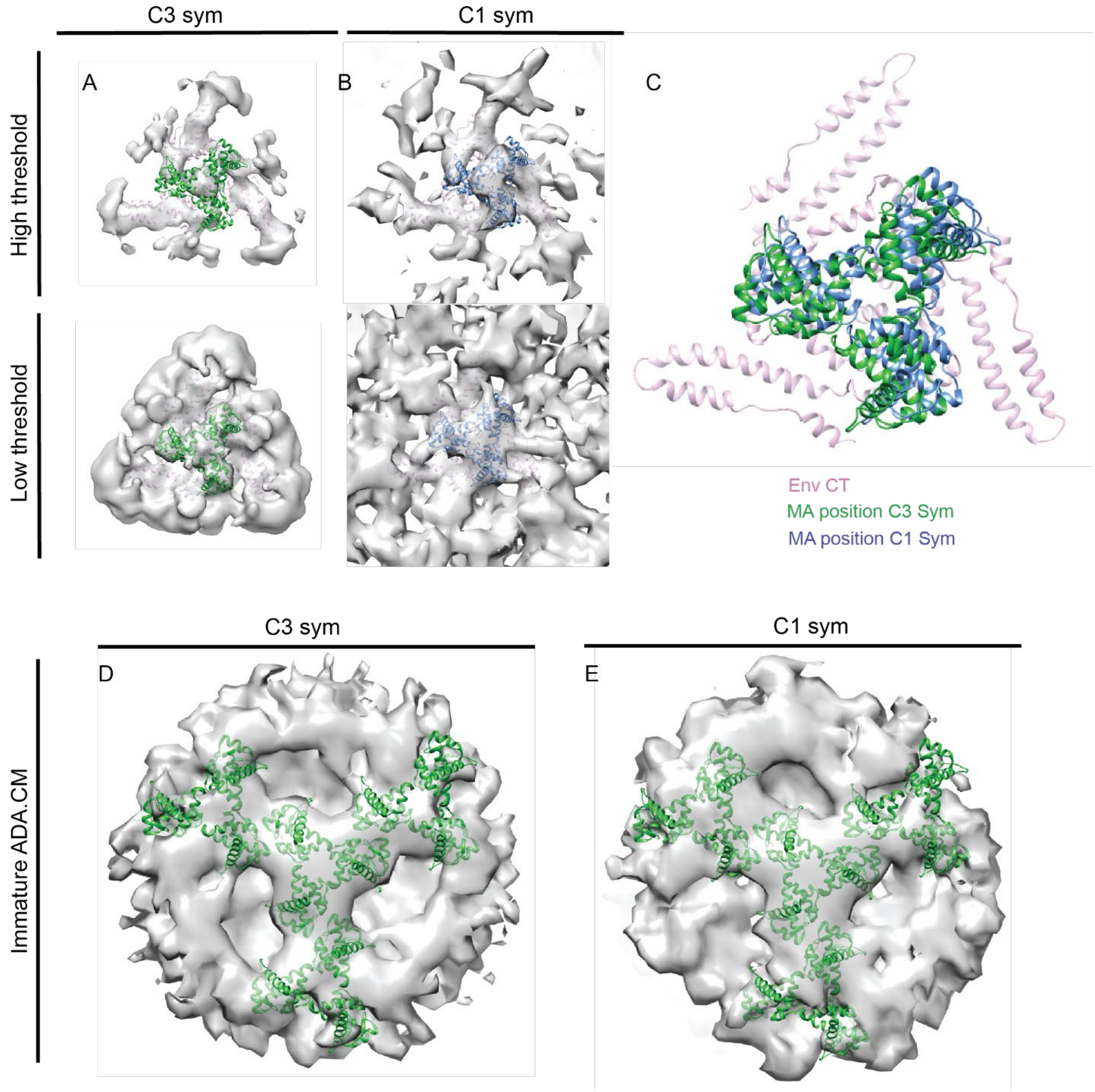


**Figure S5. Comparison of C3 and C1-relaxed symmetric reconstructions of the CH505.N197D Env CT-MA interaction. (A)** Subtomogram average of the C3 symmetric reconstruction of the CH505.N197D Env CT-MA interaction described in the main text (Fig 2) at high density threshold (top) showing the fit of the Env CT model to the map, and low density threshold (bottom) showing the fit of the MA trimer to the map (green). **(B)** Using the C3 refinement, symmetry relaxation was performed. Map is shown at a high density threshold (top) showing the fit of the Env CT model to the map (pink), and low density threshold (bottom) showing the fit of the MA trimer to the map (blue). **(C)** Superposition of the fitted MA trimers in the C3 reconstruction (green MA trimer) and C1-relaxed reconstruction (blue). The center of the MA trimer is shifted 4.6 Å between the maps, which at the resolution of the maps is not a significant difference. **(D)** Subtomogram average of the C3 symmetric reconstruction of the immature ADA.CM.755* MA lattice interaction described in the main text (Fig 3G). **(E)** From the C3 map, symmetry relaxation to C1was performed. The fit of the immature MA lattice does not change between the C3 symmetric and asymmetric maps. For panels **D** and **E**, the immature MA lattice model (PDB: 7OVQ) was used.

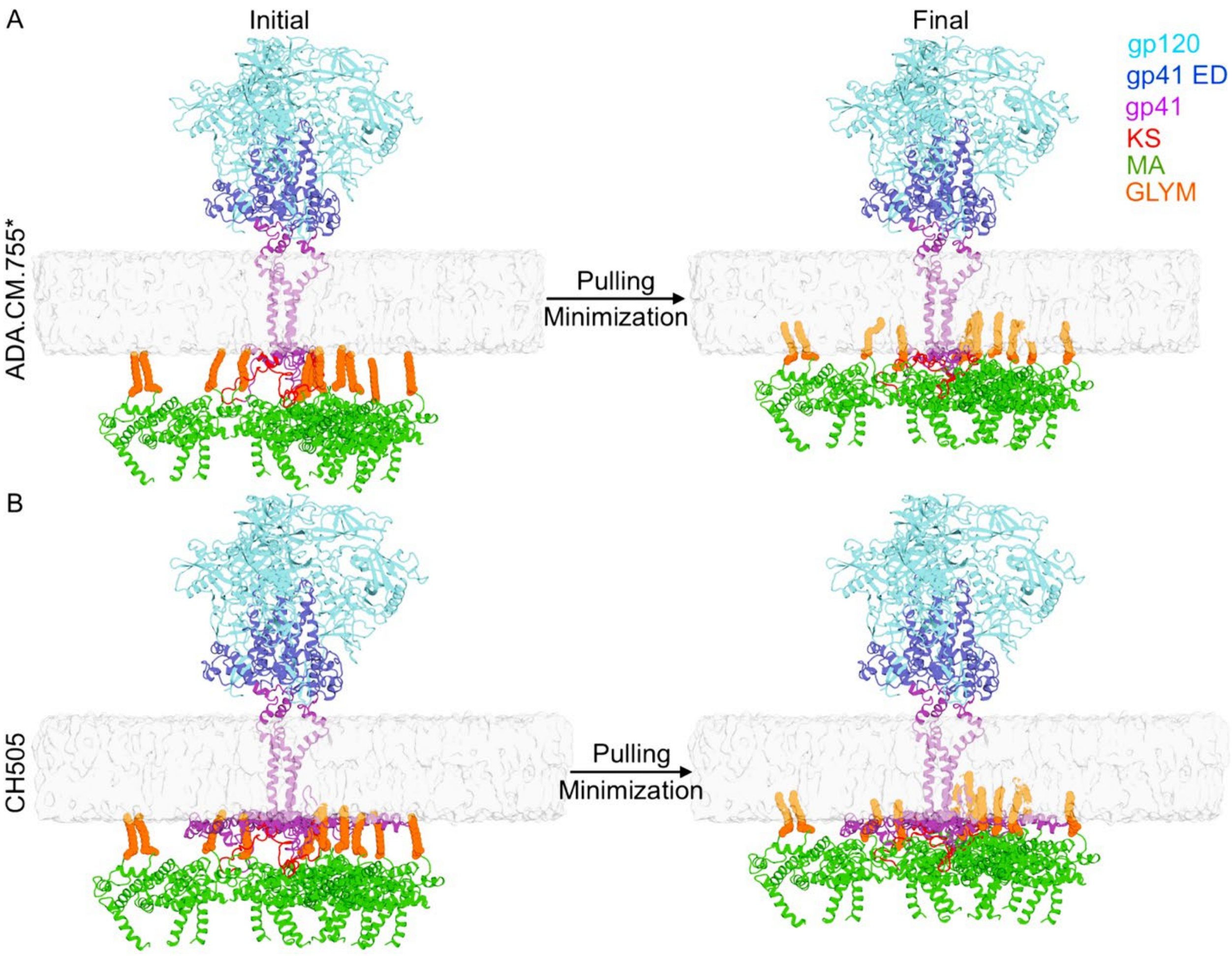


**Figure S6. Insertion of the myristoylation motifs (GLYM) of the immature MA protein to obtain the initial complexes of ADA.CM.755* and CH505 with immature MA by pulling and energy minimization simulations.** The MA was pulled upwards 5 Å at a time before the whole system was energetically minimized. The gp120 domain is colored cyan, gp41 ectodomain (ED) is colored cornflower blue, gp41 MPER, TM, and CT domains are colored magenta, KSs are colored red, myristoylation motifs (GLYM) are colored orange, the MA is colored green, and the membrane is colored gray.

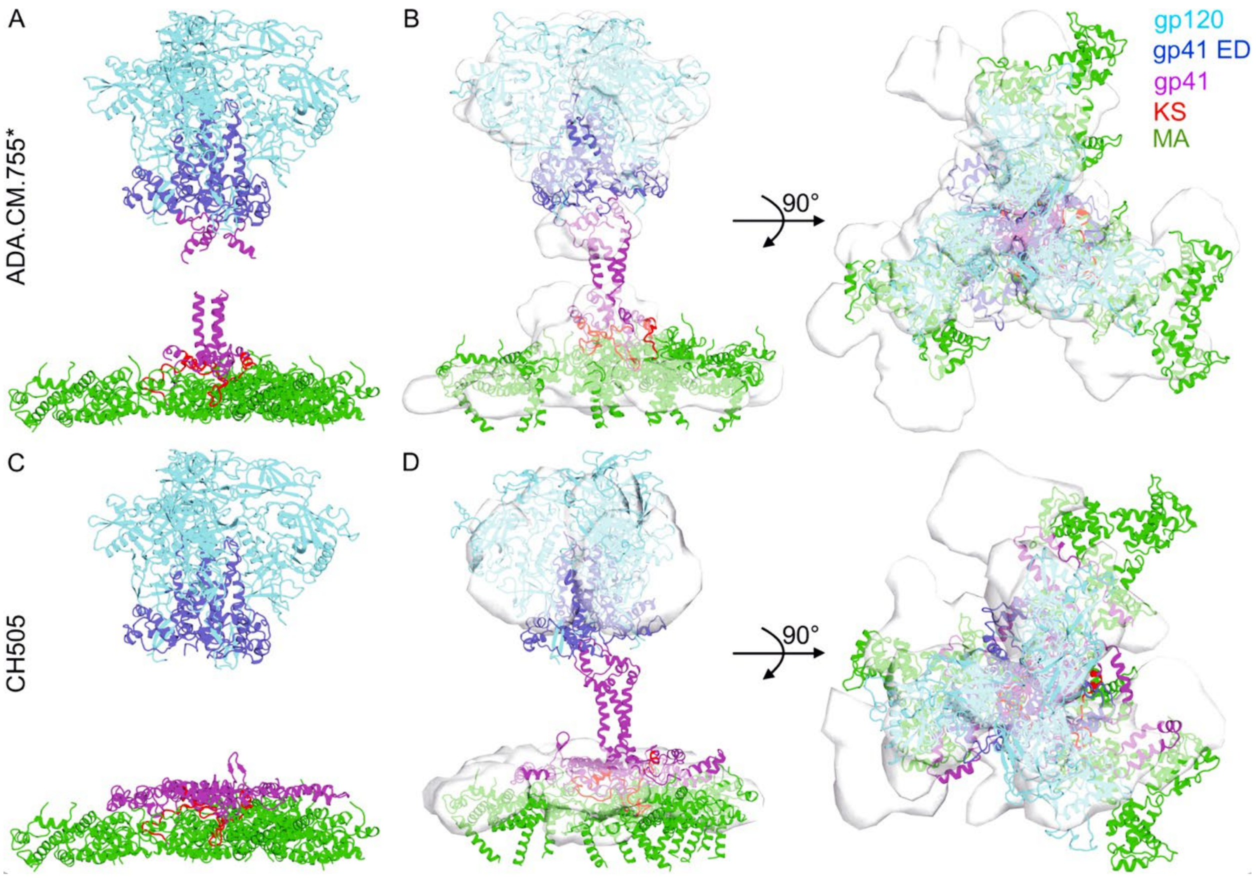


**Figure S7. Fitting of the ADA.CM.755* and CH505 Env in complex with immature MA protein incorporated to the experimental cryo-ET density by density-guided MD simulations.** **(A)** Regions of ADA.CM.755* in complex with immature MA protein that were fit to the experimental cryo-ET density. **(B)** Final density-guided conformation of ADA.CM.755* in complex with the restrained immature MA protein. **(C)** Regions of CH505 in complex with immature MA protein that were fit to the experimental cryo-ET density. **(D)** Final density-guided conformation of CH505 in complex with the restrained immature MA protein. The gp120 domain is colored cyan, gp41 ectodomain (ED) is colored cornflower blue, gp41 MPER, TM, and CT domains are colored magenta, Kennedy's sequences (KSs) are colored red, the MA is colored green, and the cryo-ET density is colored gray

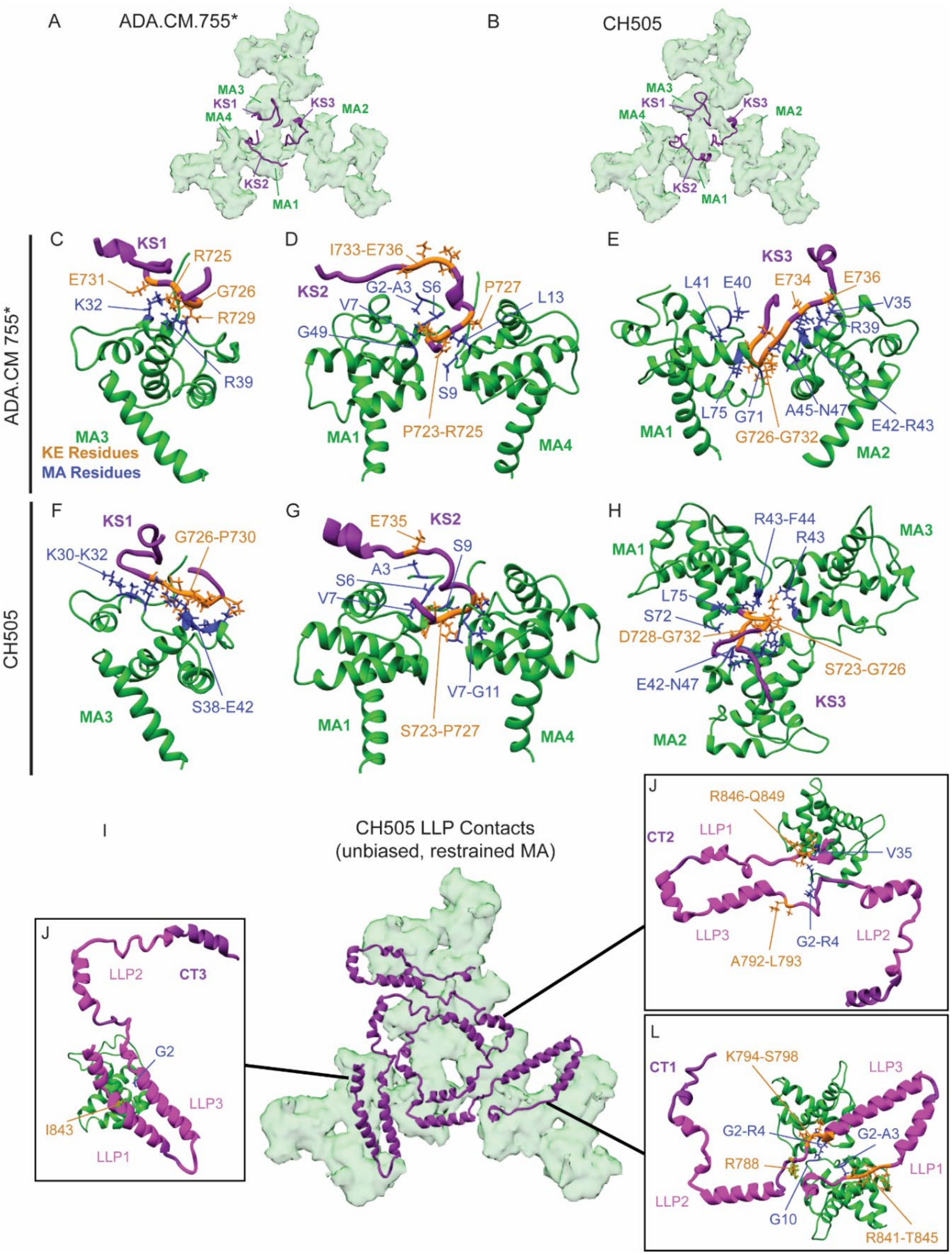


**Figure S8. Interactions between Env CT and the immature SG3 Gag MA protein captured from unbiased MD simulations of ADA.CM.755* and CH505 in complex with the restrained immature MA. (A)** Representative KS positions relative to the restrained immature MA in the unbiased MD simulations of ADA.CM.755*. **(B)** Representative KS positions relative to the restrained immature MA in the unbiased MD simulations of CH505. **(C)** Important residues involved in the interactions between KS1 of ADA.CM.755* and MA3 of the central MA trimer. **(D)** Important residues involved in the interactions between KS2 of ADA.CM.755* MA1 of the central MA trimer and MA4 of the perimeter MA trimer. Important residues involved in the interactions between KS3 of ADA.CM.755* and MA1 and MA2 of the central MA trimer. **(F)** Important residues involved in the interactions between KS1 of CH505 and MA3 of the central MA trimer. **(G)** Important residues involved in the interactions between KS2 of CH505 and MA1 of the central MA trimer and MA4 of the perimeter MA trimer. **(H)** Important residues involved in the interactions between KS3 of CH505 and MA1, MA2, and MA3 of the central MA trimer. **(I)** Representative gp41 CT positions relative to the restrained immature MA in the unbiased MD simulations. **(J)** Important residues involved in the interactions between the CT3 and MA3 of the central MA trimer. **(K)** Important residues involved in the interactions between CT2 and MA12 of the perimeter MA trimer. **(L)** Important residues involved in the interactions between CT1 and MA2 of the central trimer as well as MA8 of the perimeter MA trimer. The residue contacts shown in the figure had contact frequencies ≥ ~0.5 during the MD simulations. Throughout the figure KSs are colored red, the gp41 CTs are colored pink, the LLPs are colored magenta, the MA protein is colored green, the important residues of the CTs involved in the CT-MA interactions are colored orange, and the important residues of the MA involved in the CT-MA interactions are colored blue. The residue numbers were in accordance with the HXB2 numbering scheme.

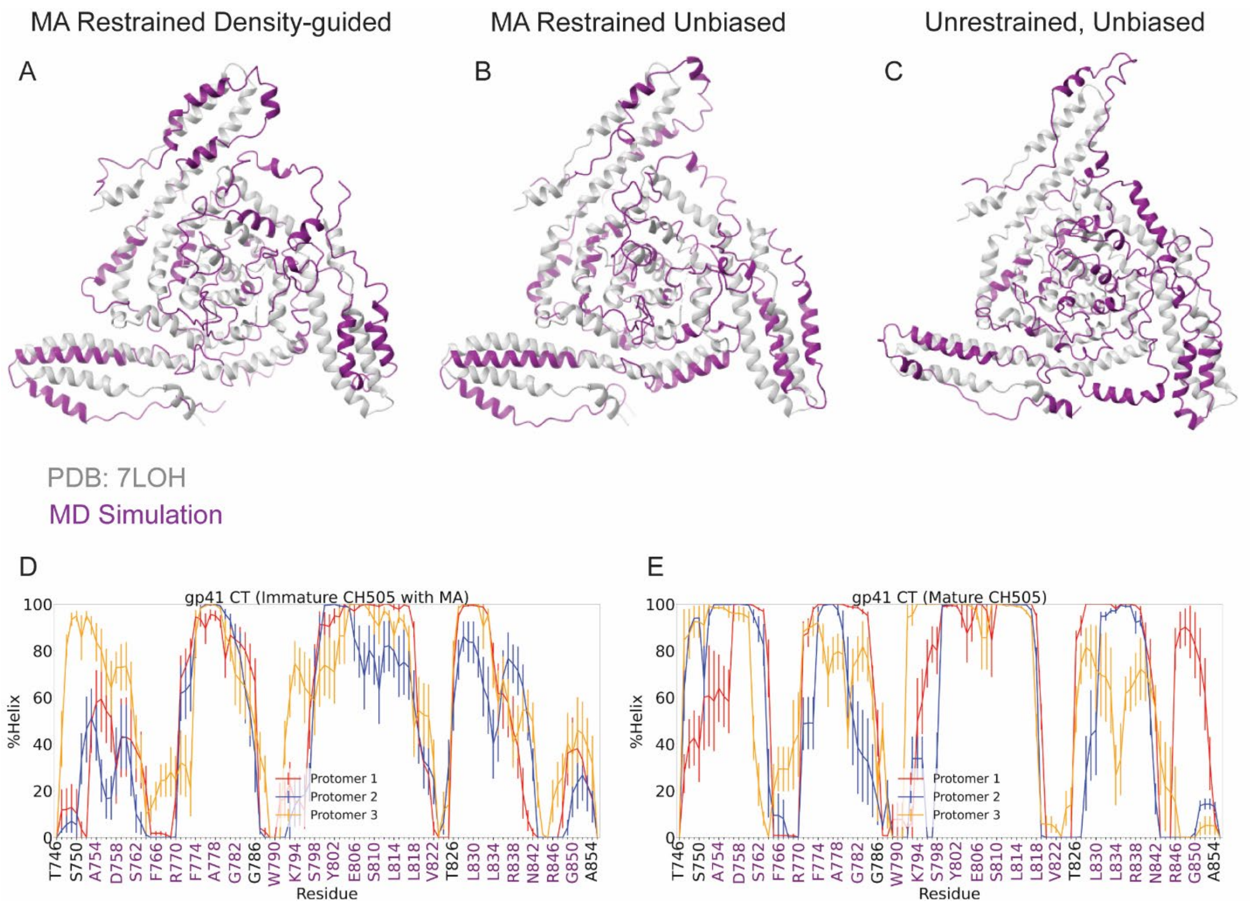


**Figure S9. Comparison of Env CT NMR Model to Final Conformations from MD simulations.** Env CT structure previously determined by NMR (PDB: 7LOH) (1) compared to the final conformations from the MD simulations. **(A)** Density-guided with MA lattice restrained **(B)** Unbiased with MA lattice restrained. **(C)** Unbiased with no MA lattice restraints. **(D-E)** Secondary structure of gp41 CT sampled from the MD simulations of CH505 with immature MA **(D)** and without immature MA protein **(E)**. Here, residues constituting the lentivirus lytic peptides (LLP), including HXB2 residues 753-785 (LLP-2), 790-823 (LLP-3), as well as 827-842 and 846-853 (LLP-1), are colored purple. The error bars represent the standard errors of means (S.E.m.) from different simulation replicas.

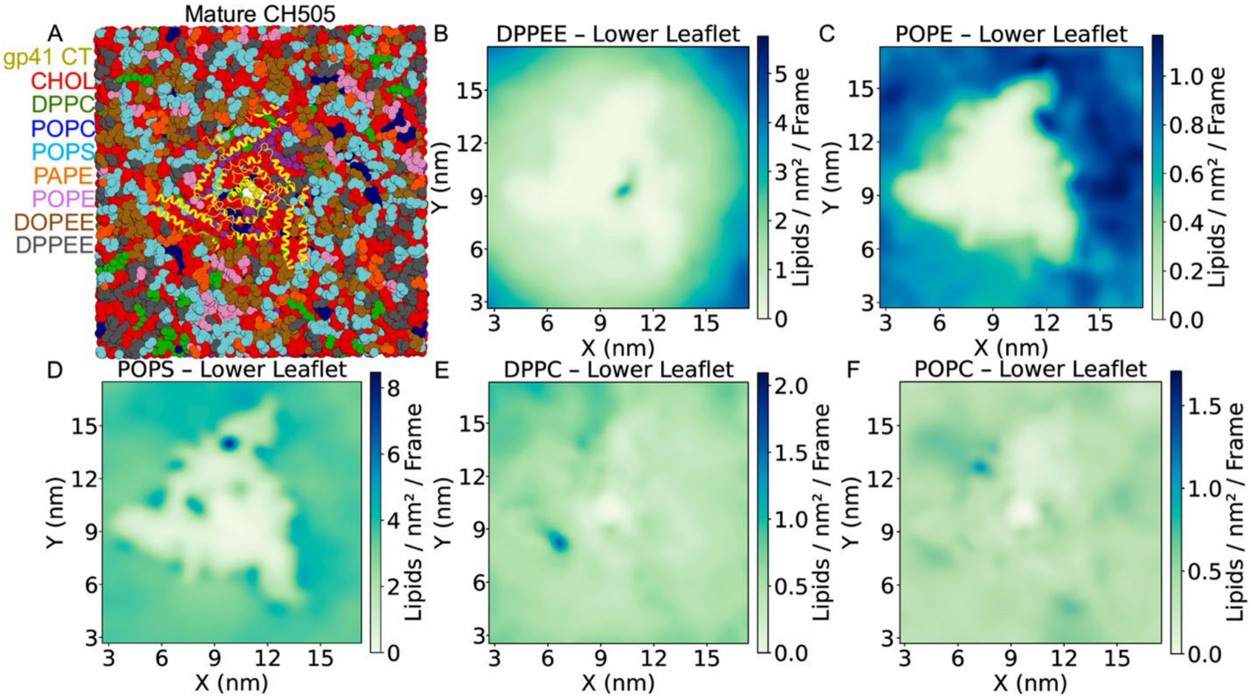


**Figure S10. Distributions of lipid molecules in the lower leaflet in the last 5µs of the CG simulations of mature CH505. (A)** Representative conformation of the CH505 gp41 CT in the CG simulated HIV-1 membrane taken from one last simulation frame of the CG simulations. Since the HIV-1 proteins were completely restrained during the CG simulations, the atomistic conformations of the HIV-1 proteins before the CG simulations were used for better visualizations. The gp41 CT is colored yellow, CHOL lipid molecules are colored red, DPPC are colored green, POPC are colored blue, POPS are colored cyan, PAPE are colored orange, POPE are colored light pink, DOPEE are colored brown, and DPPEE are colored gray. **(B-F)** Distributions of DPPEE **(B)**, POPE **(C)**, POPS **(D)**, DPPC **(E)**, and POPC **(F)** in the lower leaflet in the last 5µs of the CG simulations of mature CH505. A color scheme of green – blue was used to illustrate the lowest – highest average density of the lipid molecules in the lower leaflet in the simulations.

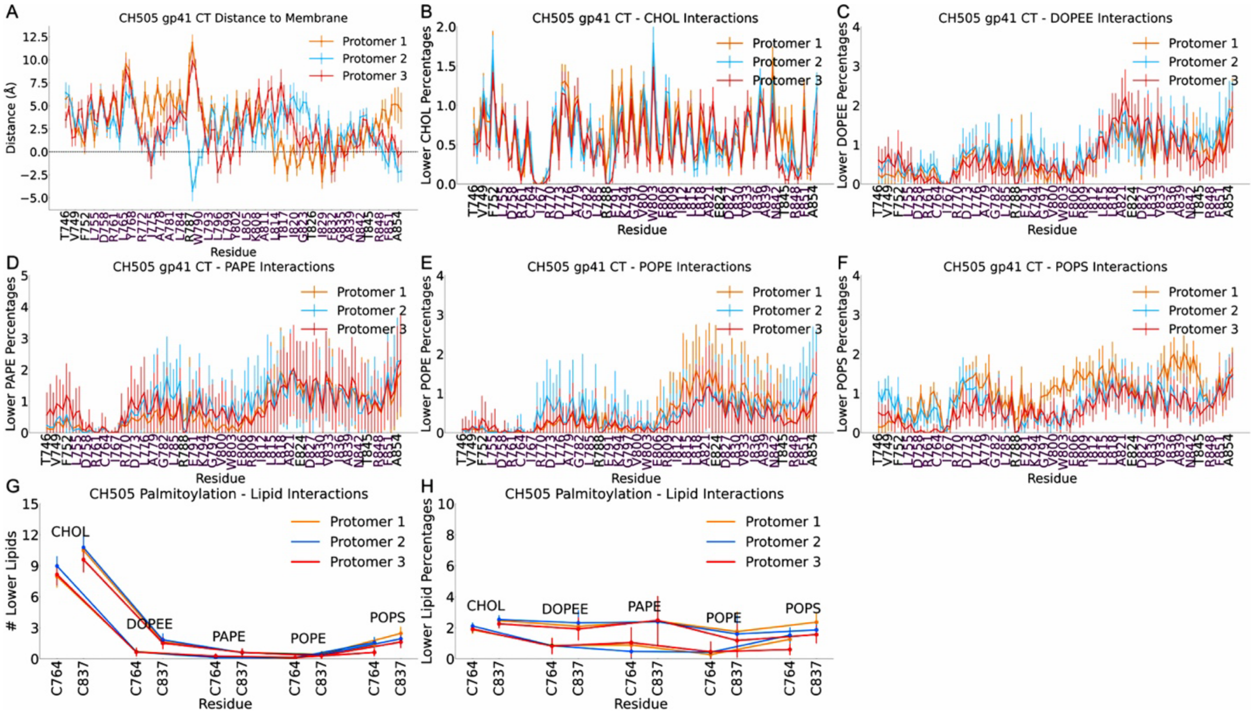


**Figure S11. Interactions of the gp41 CT residues of CH505 with lipid molecules in the lower leaflet of the HIV-1 membrane. (A)** Average distances of the CH505 gp41 cytoplasmic tails (CT) relative to the phosphate lipid head groups in the membrane lower leaflet sampled from the AA MD simulations of CH505 in complex with immature MA protein in the model HIV-1 membrane. **(B-F)** Interactions of the gp41 CT residues of CH505 with CHOL **(B)**, DOPEE **(C)**, PAPE **(D)**, POPE **(E)**, and POPS **(F)** determined from the last 5μs of CG simulations of mature CH505 in the model HIV-1 membrane. **(G-H)** Interactions of the palmitoylation motifs of CH505 gp41 CT (C764 and C837) with CHOL, DOPEE, PAPE, POPE, and POPS shown as numbers **(G)** and percentages **(H)** of the lipid types in the lower leaflet determined from the last 5μs of CG simulations of mature CH505 in the model HIV-1 membrane. A cutoff distance of 10 Å was used to determine the numbers and percentages of neighboring lipid molecules to the protein beads of gp41 CT residues. Here, residues constituting the lentivirus lytic peptides (LLP), including HXB2 residues 753-785 (LLP-2), 790-823 (LLP-3), as well as 827-842 and 846-853 (LLP-1), are colored purple. The error bars represent the standard errors of means (S.E.m.) from different simulation replicas.

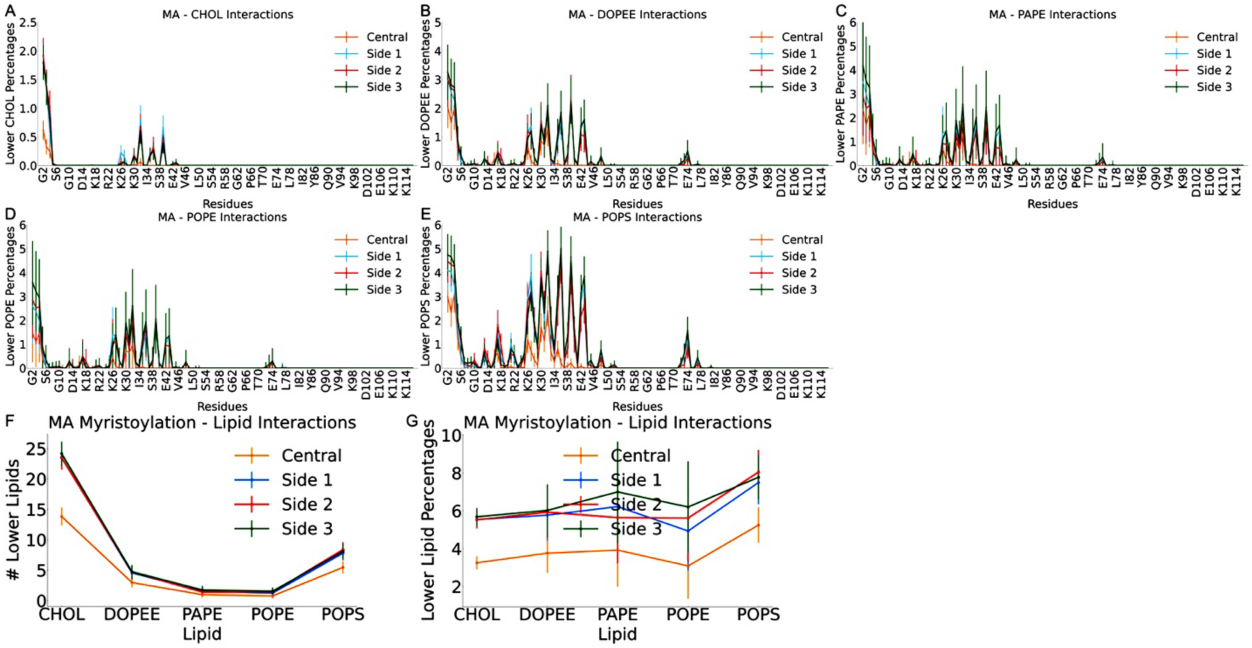


**Figure S12. Interactions of the MA residues with lipid molecules in the lower leaflet captured from the last 5μs of CG simulations of ADA.CM.755* with immature MA. (A-E)** Interactions of MA residues with CHOL **(A)**, DOPEE **(B)**, PAPE **(C)**, POPE **(D)**, and POPS **(E)**. **(F-G)** Interactions of the myristoylation motifs of MA protein (G2) with CHOL, DOPEE, PAPE, POPE, and POPS shown as numbers **(F)** and percentages **(G)** of the lipid types in the lower leaflet. A cutoff distance of 10 Å was used to determine the numbers and percentages of neighboring lipid molecules to the protein beads of MA residues. The error bars represent the S.E.m. from different simulation replicas.

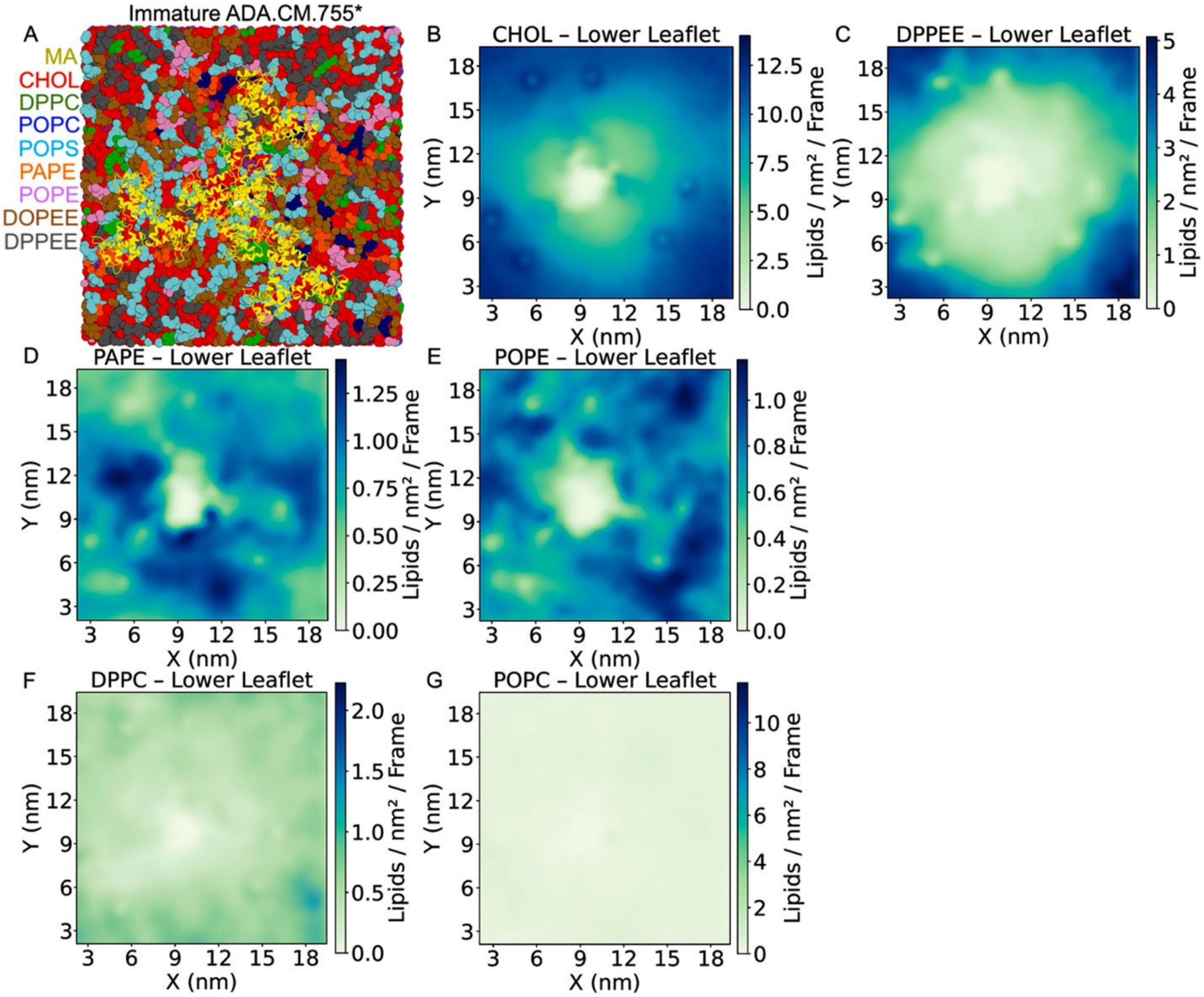


**Figure S13. Distributions of lipid molecules in the lower leaflet in the last 5μs of the CG simulations of ADA.CM.755* with immature MA. (A)** Representative conformation of the immature MA protein in the CG simulated HIV-1 membrane taken from one last simulation frame of the CG simulations. Since the HIV-1 proteins were completely restrained during the CG simulations, the atomistic conformations of the HIV-1 proteins before the CG simulations were used for better visualizations. The MA is colored yellow, CHOL lipid molecules are colored red, DPPC are colored green, POPC are colored blue, POPS are colored cyan, PAPE are colored orange, POPE are colored light pink, DOPEE are colored brown, and DPPEE are colored gray. **(B-G)** Distributions of CHOL **(B)**, DPPEE **(C)**, PAPE **(D),** POPE **(E),** DPPC **(F)**, and POPC **(G)** in the lower leaflet in the last 5μs of the CG simulations of ADA.CM.755* with immature MA. A color scheme of green – blue was used to illustrate the lowest – highest average density of lipid molecules in the lower leaflet in the simulations.

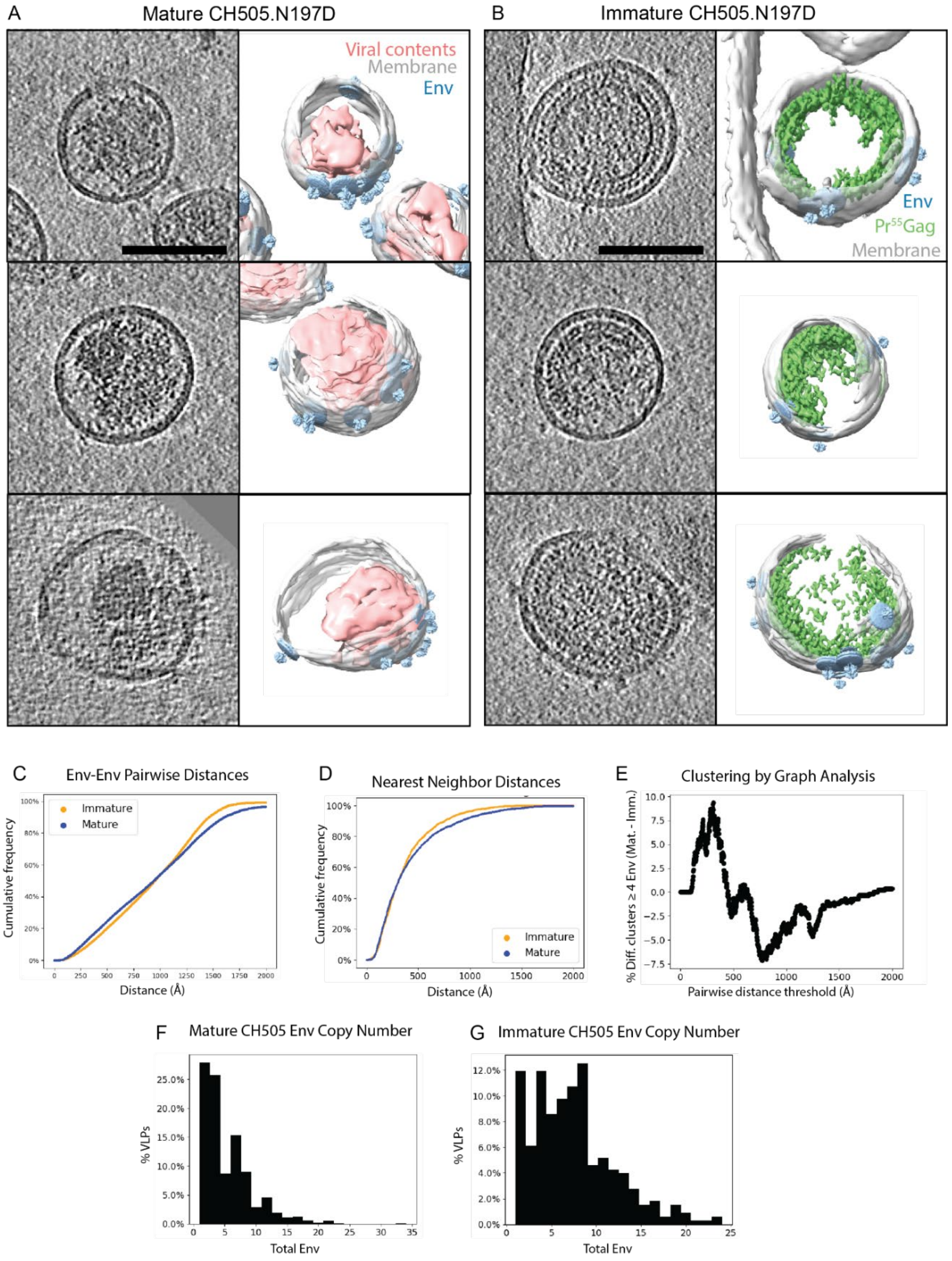


**Figure S14. CH505.N197D Env Clustering Analysis. (A)** Representative 10 nm thick tomographic slices (left) and segmentations (right) of mature HIV-1 VLPs bearing CH505.N197D Env. In the segmentations, the viral membrane is colored grey, refined Env subtomograms are projected back into their context and colored blue, and the dense viral contents are colored red. Due to AT-2 treatment as a safety precaution, the mature conical CA core is not visible. Scale bar 100 nm. **(B)** Representative 10 nm thick tomographic slices (left) and segmentations (right) of immature HIV-1 VLPs bearing CH505.N197D Env. In the segmentations, the viral membrane is colored grey, refined Env subtomograms are projected back into their context and colored blue, and the immature $PR55^{Gag}$ lattice is colored green. Scale bar 100 nm. **(C)** Cumulative frequency of Env-Env pairwise distances (Å) for mature (blue) and immature (orange) samples. **(D)** Cumulative frequency of Env-Env nearest neighbor distances (Å) for mature (blue) and immature (orange) samples. **(E)** Env clustering was analyzed by graph analysis, where Env within a given distance threshold formed connected nodes on a graph. A connected set of nodes was considered clustered. Difference plot (mature – immature) of percentage of VLPs with a largest cluster of at least 4 Env is plotted against distance threshold. At small distances, mature virions displayed greater Env clustering. At larger distances mature virions displayed lesser Env clustering, which we reasoned was due to lower numbers of Env per virion. **(F, G)** Histograms showing Env copy number on mature and immature VLPs, respectively.

| Env | Glycan Type | Env Residues |
|---|---|---|
| ADA.CM.755* | MAN-9 | N130, N136, N139, N156, N160, N234, N241, N262, N295, N332, N339, N356, N362, N386, N392, N401, N411, N448 |
| | FA2 | N88, N188, N197, N276, N301, N397, N406, N461, N611, N616, N625, N637 |
| CH505 | MAN-9 | N130, N133, N141, N156, N160, N230, N241, N262, N289, N334, N339, N358, N386, N392, N398, N404, N409, N448 |
| | FA2 | N88, N187, N197, N276, N301, N442, N461, N611, N616, N625, N637 |

**Table S1. Glycosylation profile of the ADA.CM.755* and CH505 envelope (Env) proteins modelled based on the experimental glycosylation profile of BG505 SOSSIP.664 trimer.** The residue numbers were in accordance with the HXB2 numbering scheme.

| Kennedy Sequence Residues | | MA Residues | | KS-MA Residue Contacts |
|---|---|---|---|---|
| **KS1** | R725, G726, D728, R729, P730, E731 | **MA3** | A3, K32, V35, R39, E40 | G726 – R39 (0.70), P730 – A3 (0.57), R725 – R39 (0.56), E731 – K32 (0.46), R729 – K32 (0.34), R725 – E40 (0.25), D728 – V35 (0.25) |
| **KS2** | T723, P724, R725, G726, P727, I733, E734, E735, E736 | **MA1** | G2, S6, V7, G49 | P724 – G49 (0.90), R725 – S6 (0.66), E736 – G2 (0.59), E735 – G2 (0.56), T723 – G49 (0.51), E734 – G2 (0.43), G726 – S6 (0.42), P724 – S6 (0.39), P724 – V7 (0.35), I733 – G2 (0.32) |
| | | **MA4** | A5, S9, G10, G11, L13 | G726 – A5 (0.50), R725 – S9 (0.44), T723 – G11 (0.40), P727 – L13 (0.39), R725 – G10 (0.24) |
| **KS3** | P724, R725, G726, P727, D728, R729, P730, E731, G732, I733, E734, E736 | **MA1** | E40, L41, R43, G71, E74, L75 | G726 – E40 (0.97), P727 – L75 (0.96), G726 – L41 (0.81), P724 – R43 (0.73), D728 – L75 (0.70), P727 – E74 (0.70), R729 – G71 (0.42), D728 – G71 (0.37), P730 – G71 (0.31), R725 – R43 (0.23) |
| | | **MA2** | V35, R39, E42, R43, F44, A45, V46, N47 | G732 – E42 (1.0), P730 – A45 (0.85), E731 – R43 (0.84), R729 – A45 (0.68), E731 – E42 (0.67), G732 – R43 (0.55), P730 – V46 (0.52), P730 – R43 (0.51), P730 – N47 (0.50), E734 – R39 (0.48), G732 – N47 (0.48), E731 – N47 (0.46), P730 – F44 (0.43), I733 – E42 (0.37), E736 – V35 (0.32), E734 – R43 (0.27), G732 – N47 (0.23), E731 – N47 (0.22) |

**Table S2. Residue contacts between the KSs of ADA.CM.755* and the immature SG3 Gag MA captured from the density-guided MD simulations of ADA.CM.755* in complex with the restrained immature MA fit to the experimental cryo-ET density.** The residue contacts listed in the table had contact frequencies (listed in parentheses) ≥ 0.2 during the MD simulations. The residue numbers were in accordance with the HXB2 numbering scheme.

| Kennedy Sequence Residues | MA Residues | KS-MA Residue Contacts |
|---|---|---|

| | | | | |
|---|---|---|---|---|
| **KS1** | P724, R725, G726, P727, D728, P730, G731 | **MA3** | A3, L31, S38, R39, E40, L41, E42 | G726 – L31 (0.80), G726 – E42 (0.78), G726 – L41 (0.72), R725 – E42 (0.68), D728 – R39 (0.68), G726 – S38 (0.62), P730 – A3 (0.56), P727 – S38 (0.53), R725 – E40 (0.53), P727 – R39 (0.52), R725 – R39 (0.52), P724 – E42 (0.40), D728 – S38 (0.39), G726 – E40 (0.38), G731 – L31 (0.30) |
| **KS2** | S723, P724, R725, G726, P727, D728, P730, I733, E734, E735, E736, E739 | **MA1** | G2, A3, R4, S6, V7, K32 | R725 – S6 (0.99), E735 – G2 (0.99), E734 – G2 (0.98), I733 – G2 (0.97), E736 – G2 (0.96), E735 – A3 (0.42), R725 – V7 (0.41), I733 – A3 (0.39), E735 – R4 (0.36), P724 – V7 (0.35), E736 – A3 (0.35), E739 – K32 (0.22), E734 – A3 (0.20) |
| | | **MA4** | A5, S6, V7, L8, S9, G10, G11, E12, L13 | R725 – S9 (1.0), R725 – L8 (1.0), G726 – V7 (1.0), P727 – G10 (0.99), G726 – S6 (0.99), G726 – A5 (0.98), P724 – S9 (0.97), G726 – G10 (0.97), G726 – S9 (0.96), S723 – G10 (0.95), R725 – S6 (0.89), G726 – L8 (0.87), D728 – L13 (0.85), P730 – L13 (0.85), S723 – S9 (0.84), R725 – V7 (0.81), S723 – G11 (0.79), R725 – G10 (0.64), P730 – E12 (0.49), P724 – G10 (0.47), G731 – L13 (0.44) |
| **KS3** | P724, R725, G726, P727, D728, R729, P730, G731, G732, E734 | **MA1** | R43, F44, L64, G71, S72, L75 | P730 – G71 (0.68), P730 – S72 (0.64), P724 – F44 (0.58), D728 – L75 (0.52), P724 – R43 (0.51), D728 – L64 (0.47), P727 – L75 (0.20) |
| | | **MA2** | R39, E42, R43, F44, A45, V46, N47, P48 | G731 – E42 (0.99), P730 – N47 (0.94), E734 – R39 (0.88), R725 – R43 (0.76), G731 – N47 (0.76), R729 – F44 (0.68), R729 – A45 (0.65), G732 – E42 (0.57), G726 – R43 (0.55), G731 – R43 (0.47), P730 – V46 (0.42), P730 – A45 (0.38), G731 – P48 (0.29), G731 – V46 (0.29), R729 – R43 (0.29), G732 – P48 (0.23), E734 – E42 (0.22) |
| | | **MA3** | R43 | P724 – R43 (0.82), R725 – R43 (0.21) |

**Table S3. Residue contacts between the KSs of CH505 and the immature SG3 Gag MA captured from the density-guided MD simulations of CH505 in complex with the restrained**

**immature MA fit to the experimental cryo-ET density.** The residue contacts listed in the table had contact frequencies (listed in parentheses) ≥ 0.2 during the MD simulations. The residue numbers were in accordance with the HXB2 numbering scheme.

| GP41 C-Terminal Residues | | MA Residues | | CT-MA Residue Contacts |
|---|---|---|---|---|
| **CT1** | F766$^{LLP2}$, I767$^{LLP2}$, R788, L793$^{LLP3}$, K794$^{LLP3}$, Y795$^{LLP3}$, L796$^{LLP3}$, G797$^{LLP3}$, S798$^{LLP3}$, L799$^{LLP3}$, I843, P844, T845, R846$^{LLP1}$, I847$^{LLP1}$, G850$^{LLP1}$, T853$^{LLP1}$ | **MA1** | R4, L31 | F766 – R4 (0.48), I767 – L31 (0.21), I767 – R4 (0.21) |
| | | **MA2** | G2, A3, R4 | K794 – G2 (0.99), G797 – G2 (0.94), Y795 – G2 (0.68), S798 – G2 (0.68), S798 – A3 (0.67), L796 – G2 (0.46), Y795 – A3 (0.40), S798 – R4 (0.36), L799 – A3 (0.24), L793 – G2 (0.21) |
| | | **MA8** | G2, A3, G10, E17, K30, L31 | T845 – G2 (0.99), R846 – A3 (0.86), R846 – G2 (0.76), T845 – A3 (0.51), P844 – G2 (0.45), R788 – G10 (0.44), G850 – K30 (0.41), P844 – A3 (0.40), T853 – E17 (0.39), G850 – L31 (0.32), I847 – A3 (0.27), I843 – G2 (0.26) |
| **CT2** | A792$^{LLP3}$, L793$^{LLP3}$, K794$^{LLP3}$, S813$^{LLP3}$, D816$^{LLP3}$, G825, R846$^{LLP1}$, I847$^{LLP1}$, R848$^{LLP1}$, Q849$^{LLP1}$ | **MA11** | G25, K26, Q28 | G825 – Q28 (0.26), S813 – G25 (0.22), D816 – K26 (0.21) |
| | | **MA12** | G2, A3, R4, K32, V35 | L793 – A3 (0.94), A792 – A3 (0.83), A792 – G2 (0.78), A792 – R4 (0.74), L793 – R4 (0.66), R846 – V35 (0.55), K794 – R4 (0.55), L793 – G2 (0.49), K794 – R4 (0.48), I847 – G2 (0.44), R848 – G2 (0.42), I847 – A3 (0.29), Q849 – K32 (0.24) |
| **CT3** | S750, Q801$^{LLP3}$, Y802$^{LLP3}$ | **MA3** | G2 | S750 – G2 (0.43) |
| | | **MA4** | G2 | Q801 – G2 (0.19), Y802 – G2 (0.19) |

**Table S4. Residue contacts between the gp41 CTs of CH505 and the immature SG3 Gag MA captured from the density-guided MD simulations of CH505 in complex with the restrained immature MA fit to the experimental cryo-ET density.** The residue contacts listed in the table had contact frequencies (listed in parentheses) ≥ 0.2 during the MD simulations. The residue numbers were in accordance with the HXB2 numbering scheme.

| Kennedy Sequence Residues | | MA Residues | | KS-MA Residue Contacts |
|---|---|---|---|---|
| **KS1** | R725, G726, P727, R729, P730, E731, G732 | **MA3** | A3, L31, K32, R39, E40 | G726 – R39 (0.81), R725 – R39 (0.58), R729 – K32 (0.54), E731 – K32 (0.52), P727 – R39 (0.39), E731 – L31 (0.39), G726 – E40 (0.37), P730 – A3 (0.27), G732 – L31 (0.26), R725 – E40 (0.21) |
| **KS2** | T723, P724, R725, G726, P727, I733, E734, E735, E736, G737 | **MA1** | G2, A3, R4, S6, V7, L31, K32, G49 | R725 – S6 (0.96), P724 – G49 (0.87), E735 – G2 (0.81), T723 – G49 (0.74), P724 – S6 (0.70), P724 – V7 (0.70), E734 – G2 (0.67), E736 – G2 (0.61), I733 – G2 (0.58), E735 – A3 (0.49), E735 – R4 (0.35), G737 – K32 (0.34), E734 – R4 (0.32), E736 – A3 (0.27), G737 – L31 (0.21) |
| | | **MA4** | A5, S9, L13 | P727 – L13 (0.53), R725 – S9 (0.43), G726 – A5 (0.40) |
| **KS3** | P724, G726, P727, D728, R729, P730, E731, G732, I733, E734, E736 | **MA1** | E40, L41, R43, S67, T70, G71, S72, E74, L75 | P727 – L75 (0.97), R729 – G71 (0.87), G726 – E40 (0.73), P730 – G71 (0.65), G726 – L41 (0.60), D728 – G71 (0.50), D728 – L75 (0.47), P727 – E74 (0.38), P724 – R43 (0.34), D728 – S67 (0.33), P730 – S72 (0.27), R729 – T70 (0.24) |
| | | **MA2** | G2, V35, W36, R39, E42, R43, A45, V46, N47, P48 | R729 – A45 (1.0), G732 – E42 (1.0), P730 – V46 (1.0), G732 – N47 (1.0), P730 – A45 (1.0), E731 – N47 (0.98), E731 – E42 (0.94), P730 – N47 (0.88), E731 – R43 (0.82), E731 – V46 (0.69), E734 – R39 (0.61), E736 – V35 (0.43), I733 – E42 (0.41), E736 – W36 (0.37), E736 – G2 (0.37), G737 – G2 (0.30), E734 – R43 (0.26), E731 – A45 (0.25), G732 – P48 (0.21), E734 – E42 (0.21) |

**Table S5. Residue contacts between the KSs of ADA.CM.755* and the immature SG3 Gag MA captured from the unbiased MD simulations of ADA.CM.755* in complex with the restrained immature MA.** The residue contacts listed in the table had contact frequencies (listed in parentheses) ≥ 0.2 during the MD simulations. The residue numbers were in accordance with the HXB2 numbering scheme.

| Kennedy Sequence Residues | | MA Residues | | KS-MA Residue Contacts |
|---|---|---|---|---|
| **KS1** | P724, R725, G726, P727, D728, R729, P730, G731, G732, E736, G737 | **MA3** | G2, A3, K30, L31, K32, S38, R39, E40, L41, E42, R43 | G731 – L31 (0.94), G731 – K32 (0.92), P730 – A3 (0.86), G726 – R39 (0.82), R729 – A3 (0.82), R725 – R39 (0.79), R729 – K32 (0.77), G726 – L41 (0.76), G726 – E42 (0.73), R725 – E40 (0.67), G726 – S38 (0.62), G726 – E40 (0.60), P724 – E42 (0.58), R725 – E42 (0.56), P727 – R39 (0.56), P730 – K32 (0.54), D728 – A3 (0.53), G731 – K30 (0.49), P727 – S38 (0.48), E736 – A3 (0.46), G732 – K32 (0.43), P724 – R43 (0.40), E736 – G2 (0.34), G732 – K30 (0.34), G737 – A3 (0.34) |
| **KS2** | S723, P724, R725, G726, P727, D728, P730, E734, E735, E736, E739 | **MA1** | G2, A3, S6, V7, S9, K32, G49, L50 | R725 – S6 (0.99), P724 – S6 (0.96), R725 – S9 (0.82), P724 – V7 (0.72), E735 – A3 (0.65), E734 – G2 (0.43), S723 – L50 (0.43), E735 – G2 (0.42), E739 – K32 (0.38), E736 – G2 (0.32), S723 – G49 (0.27), E734 – A3 (0.26), E736 – A3 (0.26) |
| | | **MA4** | A5, S6, V7, L8, S9, G10, G11, L13 | R725 – S9 (1.0), P727 – G10 (1.0), R725 – G10 (1.0), R725 – L8 (1.0), G726 – G10 (1.0), G726 – V7 (0.99), P724 – G10 (0.92), G726 – S9 (0.87), S723 – G11 (0.75), G726 – L8 (0.70), S723 – G10 (0.57), P724 – S9 (0.52), S723 – S9 (0.33), D728 – L13 (0.33), P727 – L13 (0.27), G726 – A5 (0.26), P730 – L13 (0.20), G726 – S6 (0.20) |
| **KS3** | S723, P724, R725, G726, D728, R729, P730, G731, G732, I733, E734 | **MA1** | R43, F44, G71, S72, L75 | P724 – F44 (0.94), P724 – R43 (0.84), P730 – S72 (0.82), D728 – L75 (0.72), P730 – G71 (0.51), R725 – F44 (0.44) |
| | | **MA2** | R39, E40, E42, R43, F44, A45, V46, N47 | G731 – E42 (1.0), P730 – A45 (0.99), R729 – A45 (0.93), P730 – V46 (0.88), G732 – E42 (0.84), P730 – N47 (0.83), G726 – R43 (0.83), G731 – N47 (0.81), G731 – R43 (0.67), G726 – F44 (0.64), R729 – F44 (0.64), G731 – V46 (0.63), R725 – F44 (0.61), S723 – R43 (0.56), |

| | | | | |
|---|---|---|---|---|
| | | | | E734 – R39 (0.39), P730 – F44 (0.37), I733 – E42 (0.32), E734 – E40 (0.26), R725 – R43 (0.25), G731 – F44 (0.20) |
| | | **MA3** | R43 | P724 – R43 (0.72) |

**Table S6. Residue contacts between the KSs of CH505 and the immature SG3 Gag MA captured from the unbiased MD simulations of CH505 in complex with the restrained immature MA.** The residue contacts listed in the table had contact frequencies (listed in parentheses) ≥ 0.2 during the MD simulations. The residue numbers were in accordance with the HXB2 numbering scheme.

| GP41 C-Terminal Residues | | MA Residues | | CT-MA Residue Contacts |
|---|---|---|---|---|
| **CT1** | F766$^{\text{LLP2}}$, I767$^{\text{LLP2}}$, R788, G789, K794$^{\text{LLP3}}$, Y795$^{\text{LLP3}}$, L796$^{\text{LLP3}}$, G797$^{\text{LLP3}}$, S798$^{\text{LLP3}}$, R841$^{\text{LLP1}}$, N842$^{\text{LLP1}}$, I843, P844, T845, R846$^{\text{LLP1}}$ | **MA1** | A3, L31 | F766 – A3 (0.33), I767 – L31 (0.26) |
| | | **MA2** | G2, A3, R4 | G797 – G2 (1.0), S798 – G2 (1.0), Y795 – G2 (1.0), S798 – A3 (0.99), S798 – R4 (0.98), L796 – G2 (0.90), Y795 – A3 (0.85), K794 – G2 (0.69), L796 – A3 (0.54) |
| | | **MA8** | G2, A3, G10, G11 | T845 – G2 (0.82), P844 – A3 (0.80), I843 – G2 (0.62), P844 – G2 (0.57), R841 – G2 (0.55), N842 – A3 (0.55), R788 – G10 (0.52), I843 – A3 (0.51), G798 – G10 (0.41), T845 – A3 (0.39), N842 – G2 (0.39), R846 – A3 (0.37), R846 – G2 (0.35), R788 – G11 (0.20) |
| **CT2** | A792$^{\text{LLP3}}$, L793$^{\text{LLP3}}$, I829$^{\text{LLP1}}$, T845, R846$^{\text{LLP1}}$, I847$^{\text{LLP1}}$, R848$^{\text{LLP1}}$, Q849$^{\text{LLP1}}$, G850$^{\text{LLP1}}$ | **MA11** | K27 | I829 – K27 (0.19) |
| | | **MA12** | G2, A3, R4, K32, V35, R39 | R848 – G2 (0.96), I847 – G2 (0.83), R848 – A3 (0.64), I847 – A3 (0.62), L793 – A3 (0.55), Q849 – G2 (0.49), R846 – V35 (0.47), L793 – G2 (0.47), A792 – R4 (0.47), A792 – G2 (0.46), R846 – A3 (0.45), A792 – A3 (0.41), G850 – G2 (0.41), L793 – R4 (0.38), R846 – R39 (0.30), T845 – A3 (0.26), R848 – K32 (0.20) |
| **CT3** | A754$^{\text{LLP2}}$, I843, T845, R846$^{\text{LLP1}}$ | **MA3** | G2 | A754 – G2 (0.31) |
| | | **MA4** | G2, A3 | I843 – G2 (0.40), R846 – A3 (0.30), T845 – G2 (0.28) |

**Table S7. Residue contacts between the gp41 CTs of CH505 and the immature SG3 Gag MA captured from the unbiased MD simulations of CH505 in complex with the restrained immature MA.** The residue contacts listed in the table had contact frequencies (listed in parentheses) ≥ 0.2 during the MD simulations. The residue numbers were in accordance with the HXB2 numbering scheme.

| Kennedy Sequence Residues | | MA Residues | | KS-MA Residue Contacts |
|---|---|---|---|---|
| **KS1** | P724, R725, G726, P727, D728, R729, P730, E736 | **MA3** | A3, R4, A5, S6, K32, V35, S38, R39 | R725 – R39 (0.36), P730 – A3 (0.31), G726 – R39 (0.31), G726 – S38 (0.29), R729 – R4 (0.29), R729 – A3 (0.26), G726 – V35 (0.26), P724 – R39 (0.26), E736 – K32 (0.23), R729 – A5 (0.22), D728 – A3 (0.21), D728 – S6 (0.21), P727 – S6 (0.21), R725 – A5 (0.21) |
| **KS2** | P724, R725, G726, P727, P730, E731, G732, I733, E734, E735, E736 | **MA1** | G2, A3, A5, S6, G49, L50, E52, T53, | P724 – G49 (0.65), E734 – G2 (0.50), E735 – G2 (0.50), E736 – A3 (0.36), E736 – G2 (0.32), P724 – S6 (0.28), I733 – G2 (0.28), E735 – A3 (0.25), P724 – L50 (0.24), P724 – E52 (0.24), P724 – T53 (0.23), E734 – A3 (0.23), P724 – A5 (0.23), P730 – G2 (0.22), E731 – G2 (0.22), G732 – A2 (0.21) |
| | | **MA4** | V7, L8, S9, G10, G11, E12, L13 | G726 – L8 (0.42), R725 – L8 (0.31), G726 – V7 (0.29), G726 – G11 (0.29), G726 – E12 (0.28), R725 – S9 (0.27), P727 – L13 (0.26), R725 – V7 (0.25), G726 – G10 (0.22) |
| **KS3** | P724, D728, R729, P730, E731, G732, I733 | **MA1** | R43, S67, G71, S72 | D728 – S67 (0.38), D728 – G71 (0.33), P730 – G71 (0.31), R729 – G71 (0.30), P724 – R43 (0.29), P730 – S72 (0.26) |
| | | **MA2** | E42, R43, A45 | G732 – A45 (0.64), G732 – R43 (0.36), I733 – R43 (0.32), I733 – E42 (0.29), G732 – F44 (0.29), P730 – A45 (0.28), I733 – A45 (0.27), G732 – E42 (0.24), E731 – A45 (0.21) |

**Table S8. Residue contacts between the KSs of ADA.CM.755* and the immature SG3 Gag MA captured from the unbiased MD simulations of ADA.CM.755* in complex with the unrestrained immature MA.** The residue contacts listed in the table had contact frequencies (listed in parentheses) ≥ 0.2 during the MD simulations. The residue numbers were in accordance with the HXB2 numbering scheme.

| Kennedy Sequence Residues | | MA Residues | | KS-MA Residue Contacts |
|---|---|---|---|---|
| **KS1** | P724, R725, D728, R729, G731 | **MA3** | L31, K32, I34, R39, R43 | R729 – K32 (0.38), R725 – R39 (0.28), P724 – R43 (0.28), D728 – I34 (0.23), G731 – L31 (0.22) |
| **KS2** | S723, P724, R725, G726, P727, G732, I733, E734, E735, E736, G738 | **MA1** | G2, A3, W36 | E735 – G2 (0.49), E736 – G2 (0.49), E735 – A3 (0.43), E736 – A3 (0.42), G738 – W36 (0.28), I733 – G2 (0.27), G732 – G2 (0.21), E734 – G2 (0.20) |
| | | **MA4** | V7, L8, S9, G10, G11, E12 | G726 – S9 (0.74), R725 – S9 (0.72), P727 – G10 (0.56), G726 – G10 (0.53), G726 – V7 (0.37), G726 – L8 (0.37), R725 – L8 (0.34), P727 – S9 (0.33), R725 – G10 (0.26), S723 – G11 (0.22), P724 – S9 (0.20), S723 – E12 (0.20) |
| **KS3** | S723, P724, R725, G726, D728, R729, P730, G731, G732, I733, E734 | **MA1** | R43, S67, T70, G71 | D728 – S67 (0.32), D728 – G71 (0.24), P724 – R43 (0.22), D728 – T70 (0.21) |
| | | **MA2** | R39, E42, R43, V46 | G732 – R43 (0.44), G731 – R43 (0.34), G731 – E42 (0.32), G732 – E42 (0.26), E734 – R39 (0.25), P730 – V46 (0.20) |

**Table S9. Residue contacts between the KSs of CH505 and the immature SG3 Gag MA captured from the unbiased MD simulations of CH505 in complex with the unrestrained immature MA.** The residue contacts listed in the table had contact frequencies (listed in parentheses) ≥ 0.2 during the MD simulations. The residue numbers were in accordance with the HXB2 numbering scheme.

| GP41 C-Terminal Residues | | MA Residues | | CT-MA Residue Contacts |
|---|---|---|---|---|
| **CT1** | L793$^{LLP3}$, K794$^{LLP3}$, G797$^{LLP3}$, S798$^{LLP3}$, Q801$^{LLP3}$, D816$^{LLP3}$, A819$^{LLP3}$, I820$^{LLP3}$, R841$^{LLP1}$, N842$^{LLP1}$, I843, P844, T845, G850$^{LLP1}$, A854 | **MA2** | G2, A3, R4, A5 | G797 – G2 (0.91), K794 – A3 (0.56), G797 – A3 (0.49), K794 – G2 (0.43), L793 – G2 (0.36), S798 – A3 (0.26), S798 – R4 (0.24), K794 – R4 (0.21), G797 – R4 (0.21), K794 – A5 (0.20) |
| | | **MA7** | G24, G25, K26 | D816 – G25 (0.43), I820 – G25 (0.41), A819 – G25 (0.32), I820 – K26 (0.29), D816 – G24 (0.26) |
| | | **MA8** | G2, A3, R4, Q28, L31, K32 | G850 – L31 (0.43), P844 – A3 (0.40), N842 – A3 (0.40), R841 – G2 (0.35), I843 – A3 (0.33), R841 – A3 (0.30), T845 – G2 (0.30), I843 – G2 (0.26), A854 – E17 (0.23), G850 – K32 (0.23), P844 – A5 (0.22), A854 – Q28 (0.21), P844 – G2 (0.21), Q801 – R4 (0.20) |
| **CT2** | L785$^{LLP2}$, A792$^{LLP3}$, L793$^{LLP3}$, T845, R846$^{LLP1}$, I847$^{LLP1}$, R848$^{LLP1}$, Q849$^{LLP1}$ | **MA3** | G2 | L785 – G2 (0.25) |
| | | **MA12** | G2, A3, R4, S6, V35 | R846 – A3 (0.77), R848 – G2 (0.67), I847 – G2 (0.64), A792 – G2 (0.49), I847 – A3 (0.47), R848 – A3 (0.42), T845 – A3 (0.35), L793 – R4 (0.32), L793 – A3 (0.32), A792 – R4 (0.30), A792 – A3 (0.30), R848 – V35 (0.29), T845 – G2 (0.25), L793 – G2 (0.25), R846 – G2 (0.23), R846 – S6 (0.23), Q849 – G2 (0.20) |
| **CT3** | V749, S750, I820$^{LLP3}$, A821$^{LLP3}$, V822$^{LLP3}$, G823$^{LLP3}$, E824 | **MA3** | G2 | S750 – G2 (0.44), V749 – G2 (0.24) |
| | | **MA6** | W22, P23, G24, G25, K26, K27, K30, K32, H33, W36 | A821 – K32 (0.45), A821 – H33 (0.40), G823 – K30 (0.30), G823 – G25 (0.24), A821 – G25 (0.24), G823 – G24 (0.24), V822 – G25 (0.24), A821 – W36 (0.23), I820 – G25 (0.23), G823 – P23 (0.23), E824 – P23 (0.23), G823 – W22 (0.22), A821 – K26 (0.22), I820 – K30 (0.20), V822 – K27 (0.20) |

**Table S10. Residue contacts between the gp41 CTs of CH505 and the immature SG3 Gag MA captured from the unbiased MD simulations of CH505 in complex with the unrestrained immature MA.** The residue contacts listed in the table had contact frequencies (listed in parentheses) ≥ 0.2 during the MD simulations. The residue numbers were in accordance with the HXB2 numbering scheme.

| MA Residues | | | | Changes in MA-MA Residue Contacts |
|---|---|---|---|---|
| **MA1** | A45, Q59, Q108, S111, K112 | **MA3** | Q65, P66, T70, G71, S72 | A45 – S72 (-0.92), A45 – G71 (-0.70), Q59 – T70 (-0.62) |
| | | | | K112 – P66 (0.23), S111 – P66 (0.22), Q108 – P66 (0.21), K112 – Q65 (0.20) |
| **MA1** | A5, S9, G10 | **MA4** | S6 | S9 – S6 (-0.90), A5 – S6 (-0.51), G10 – S6 (-0.23) |
| **MA2** | S67, T70, G71, S72 | **MA3** | A45, V46, N47 | S72 – N47 (-0.84), T70 – V46 (-0.77), G71 – N47 (-0.76), G71 – V46 (-0.76), G71 – A45 (-0.72), T70 – N47 (-0.24), S67 – A45 (-0.23) |
| **MA2** | G2, A3, A5, S6, S9, G10, G11, G49 | **MA8** | G2, A3, R4, S6, V7, S9, G10, G11, G49 | G10 – S6 (-0.98), A5 – S6 (-0.96), S6 – S9 (-0.91), S9 – S6 (-0.67), S6 – G10 (-0.52), G49 – G11 (-0.51), G11 – G49 (-0.49), S9 – V7 (-0.49) |
| | | | | G2 – G2 (0.44), G2 – A3 (0.38), A3 – A3 (0.33), A3 – R4 (0.24), G2 – R4 (0.20) |
| **MA3** | G2, A3, R4, A5, S6, V7, S9, G10 | **MA12** | A3, R4, S6, V7, L8, S9, E52, T53, H89, Q90 | S6 – S6 (-0.92), S9 – E52 (-0.88), V7 – S9 (-0.80), A5 – S6 (-0.57), V7 – S6 (-0.55), V7 – L8 (-0.28) |
| | | | | A5 – S9 (0.56), R4 – S9 (0.51), G2 – A3 (0.35), G2 – R4 (0.33), A3 – S9 (0.30), A3 – S6 (0.25), A3 – V7 (0.23), A3 – L8 (0.22), G10 – T53 (0.21), R4 – S6 (0.21), G10 – H89 (0.20), G10 – Q90 (0.20) |
| **MA4** | T70, G71, S72 | **MA5** | A45, V46, N47, Q59, I60 | T70 – I60 (-0.31), S72 – V46 (-0.29) |
| | | | | G71 – N47 (0.73), G71 – V46 (0.70), T70 – V46 (0.67), S72 – A45 (0.64), G71 – A45 (0.56), T70 – N47 (0.35), G71 – Q59 (0.21) |
| **MA4** | V46, N47 | **MA6** | T70 | N47 – T70 (0.37), V46 – T70 (0.20) |
| **MA5** | F44, A45, T70, S72 | **MA6** | R43, A45, V46, N47 | S72 – A45 (-0.31), S72 – V46 (-0.30) |
| | | | | F44 – R43 (0.30), A45 – R43 (0.25), F44 – A45 (0.23), T70 – N47 (0.21) |
| **MA7** | T70, G71, S72 | **MA8** | A45, N47 | S72 – A45 (0.83), G71 – A45 (0.40), T70 – N47 (0.25) |
| **MA7** | A45, V46, N47 | **MA9** | T70, G71, S72 | N47 – G71 (0.33), N47 – T70 (0.30), A45 – S72 (0.29), V46 – G71 (0.25), A45 – G71 (0.24) |

| | | | | |
|---|---|---|---|---|
| **MA8** | T70, G71, S72 | **MA9** | A45, V46, N47 | T70 – V46 (0.55), G71 – N47 (0.49), T70 – N47 (0.35), G71 – V46 (0.31), S72 – A45 (0.25), G71 – A45 (0.22) |
| **MA10** | T70, G71 | **MA11** | A45, V46, N47, Q59, I60 | T70 – Q59 (-0.51), T70 – I60 (-0.29) |
| | | | | G71 – N47 (0.65), T70 – V46 (0.39), T70 – N47 (0.39), G71 – A45 (0.25) |
| **MA10** | A45, V46, N47, Q59 | **MA12** | T70, G71, S72 | Q59 – T70 (-0.42) |
| | | | | N47 – G71 (0.71), V46 – T70 (0.57), A45 – G71 (0.38), N47 – T70 (0.37), V46 – G71 (0.24), N47 – S72 (0.20) |
| **MA11** | T70 | **MA12** | N47 | T70 – N47 (0.31) |

**Table S11. Changes in MA lattice contacts from the MD simulations (both density-guided and unbiased) of ADA.CM.755* in complex with restrained vs unrestrained MA protein (unbiased).** The differences in residue contact frequencies were calculated by subtracting the residue contact frequencies determined from the simulations with restrained MA protein from those determined from the simulations with unrestrained MA. Here, we included residue contacts with absolute differences in contact frequencies ≥ 0.2 (i.e., differences in residue contact frequencies ≤ -0.2 (decreased contacts) or ≥ 0.2 (increased contacts)). The residue numbers were in accordance with the HXB2 numbering scheme.

| MA Residues | | | | Changes in MA-MA Residue Contacts |
|---|---|---|---|---|
| **MA1** | F44, A45, V46, N47, G56 | **MA3** | R43, F44, P66, S67, T70, G71, S72 | V46 – S72 (-0.64), N47 – S72 (-0.64), A45 – S72 (-0.48), V46 – G71 (-0.36) |
| | | | | G56 – P66 (0.37), V46 – S67 (0.29), N47 – T70 (0.28), F44 – R43 (0.23), A45 – F44 (0.20), G56 – T70 (0.20) |
| **MA1** | A5, S6, S9, G10, G11 | **MA4** | A3, R4, S6, V7, E52, T53 | S9 – S6 (-0.85), A5 – S6 (-0.58), S6 – S6 (-0.50), S9 – V7 (-0.35) |
| | | | | G11 – T53 (0.32), G11 – E52 (0.27), G10 – E52 (0.22), S6 – R4 (0.22), S6 – A3 (0.21) |
| **MA2** | T70, G71, S72 | **MA3** | A45, V46, N47 | S72 – N47 (-1.0), G71 – V46 (-1.0), G71 – N47 (-1.0), S72 – V46 (-0.99), T70 – V46 (-0.97), G71 – A45 (-0.49), S72 – A45 (-0.49), T70 – N47 (-0.30) |
| **MA2** | G2, A3, A5, S6, S9, G10, G11, G49 | **MA8** | A5, S6, V7, S9, G11, E52, T53 | G49 – G11 (-0.85), G10 – S6 (-0.84), S9 – E52 (-0.77), S9 – S6 (-0.72), S6 – S9 (-0.70), S9 – V7 (-0.52), A5 – S6 (-0.46), S6 – S6 (-0.43) |
| | | | | G11 – T53 (0.51), G10 – E52 (0.41), G10 – T53 (0.33), A3 – S6 (0.26), G11 – E52 (0.26), G2 – A5 (0.23), A3 – A5 (0.21) |
| **MA3** | A3, R4, A5, S6, V7, L8, S9, G10, G11, T53 | **MA12** | S6, V7, L8, S9, G10, G49, L50, E52, T53, Q90 | G10 – G49 (-0.98), S9 – T53 (-0.97), S6 – E52 (-0.96), R4 – V7 (-0.93), G11 – L50 (-0.92), S6 – L8 (-0.91), A5 – V7 (-0.90), L8 – T53 (-0.89), S9 – E52 (-0.88), V7 – T53 (-0.79), R4 – L8 (-0.79), R4 – S9 (-0.75), G11 – G49 (-0.71), L8 – E52 (-0.69), S6 – V7 (-0.66), S6 – T53 (-0.58), S9 – G49 (-0.57), A5 – E52 (-0.49), V7 – E52 (-0.48), G10 – L50 (-0.43), T53 – Q90 (-0.41), R4 – S6 (-0.33), S9 – L50 (-0.32), A5 – L8 (-0.26), R4 – A5 (-0.21) |
| | | | | A3 – G10 (0.27) |
| **MA4** | T70, G71, S72 | **MA5** | V46, N47, Q59 | S72 – V46 (-0.51), T70 – Q59 (-0.27), G71 – V46 (-0.24), S72 – N47 (-0.23) |
| | | | | G71 – N47 (0.40), T70 – V46 (0.20) |

| | | | | |
|---|---|---|---|---|
| **MA4** | A45, V46, N47 | **MA6** | T70, G71, S72 | V46 – S72 (-0.43), A45 – S72 (-0.32), N47 – S72 (-0.24), V46 – G71 (-0.22) |
| | | | | N47 – T70 (0.26) |
| **MA5** | T70, S72 | **MA6** | V46, N47 | S72 – V46 (-0.46), S72 – N47 (-0.20) |
| | | | | T70 – N47 (0.25) |
| **MA7** | T70, G71 | **MA8** | V46, N47 | T70 – V46 (0.43), G71 – V46 (0.38), G71 – N47 (0.35), T70 – N47 (0.21) |
| **MA7** | V46, N47 | **MA9** | T70, G71, S72 | V46 – S72 (-0.20) |
| | | | | N47 – T70 (0.28), N47 – G71 (0.23) |
| **MA8** | F44, T70, G71, S72 | **MA9** | A45, V46, N47, G49, L50, Q59 | S72 – V46 (-0.50), S72 – N47 (-0.33) |
| | | | | G71 – N47 (0.37), T70 – V46 (0.36), F44 – A45 (0.29), G71 – L50 (0.22), T70 – Q59 (0.22), S72 – G49 (0.22), G71 – G49 (0.22), T70 – G56 (0.21) |
| **MA10** | Q69, T70, G71, S72 | **MA11** | A45, V46, N47, G49, L50, G56 | S72 – V46 (-0.94), S72 – N47 (-0.84), G71 – A45 (-0.83), G71 – V46 (-0.82), S72 – A45 (-0.80), T70 – V46 (-0.65), G71 – N47 (-0.55) |
| | | | | T70 – G56 (0.32), Q69 – G56 (0.25), T70 – L50 (0.22), T70 – G49 (0.20) |
| **MA10** | A45, V46, N47, Q59 | **MA12** | T70, G71, S72 | Q59 – T70 (-0.50) |
| | | | | N47 – G71 (0.83), V46 – T70 (0.73), A45 – G71 (0.41), N47 – T70 (0.39), V46 – G71 (0.38), A45 – S72 (0.36) |
| **MA11** | P66, T70, G71, S72 | **MA12** | A45, V46, N47, Q59 | S72 – V46 (-0.91), S72 – A45 (-0.90), S72 – N47 (-0.52), G71 – A45 (-0.46), G71 – V46 (-0.43), G71 – N47 (-0.42), T70 – V46 (-0.42) |
| | | | | P66 – Q59 (0.21) |

**Table S12. Changes in MA lattice contacts from the MD simulations (both density-guided and unbiased) of Ch505 in complex with restrained vs unrestrained MA protein (unbiased).** The differences in residue contact frequencies were calculated by subtracting the residue contact frequencies determined from the simulations with restrained MA protein from those determined from the simulations with unrestrained MA. Here, we included residue contacts with absolute differences in contact frequencies ≥ 0.2 (i.e., differences in residue contact frequencies ≤ -0.2 (decreased contacts) or ≥ 0.2 (increased contacts)). The residue numbers were in accordance with the HXB2 numbering scheme.
